# Synergy of culture-dependent molecular identification and whole-community metabarcode sequencing for characterizing the microbiota of arable crop residues

**DOI:** 10.1101/2021.03.23.436452

**Authors:** Valérie Laval, Lydie Kerdraon, Matthieu Barret, Anne-Lise Liabot, Coralie Marais, Benjamin Boudier, Marie-Hélène Balesdent, Marion Fischer-Le Saux, Frédéric Suffert

**Author notes:** equal contributions (Valérie Laval, Lydie Kerdraon).

## Abstract

This study is the first to compare culture-dependent (strain isolation plus molecular identification) and culture-independent (whole-community metabarcode sequencing) approaches for characterizing the microbiota of crop residues. We investigated the diversity of fungal and bacterial communities in wheat and oilseed rape residues, using two different culture-dependent strategies to cover the maximum diversity for each kingdom: broad substrate sampling coupled with low-throughput isolation and diversity analysis for fungi, and reduced substrate sampling coupled with high-throughput isolation and diversity analysis for bacteria. The proportion of cultivable microorganisms was neither as low as the ‘1%’ paradigm long associated with the rhizosphere microflora, nor as high as the 50% sometimes reported for the phyllosphere microflora. It was, thus, intermediate between the values for soil and plants. This finding is consistent with residues being considered to constitute an ecotone, at the interface between soil and phyllosphere. Isolation and metabarcoding provided consistent complementary information: they revealed close community profiles, leading to the identification of several common and specific amplicon sequence variants (ASVs). The power of the culture-independent approach was thus confirmed. By contrast the culture-dependent approach was less weak than anticipated. Firstly, it provided complementary information about microbial diversity, with several ASVs not retrieved by metabarcoding being identified after isolation in the community-based culture collection. Secondly, this approach made it possible to preserve and test different taxa either individually or after the creation of synthetic communities, for deciphering the ecological functions of communities beyond merely descriptive aspects.

## Introduction

Crop residues — the decaying parts of the crop plant that are not harvested — were recently described as an ‘ecotone’: a transient half-plant/half-soil compartment constituting a key fully fledged microbial ecosystem worthy of the attention of microbiologists and plant disease epidemiologists (Smil *et al*., 1999; Kerdraon *et al*., 2019a). Crop residues have both positive and negative effects on crop health and productivity. They are the main support for the pathogenic agents responsible for residue-borne plant diseases, but they also host beneficial microorganisms and make a significant contribution to agrosystem stability (Pascault *et al*., 2010; Cobo-Díaz *et al*., 2019a; Kerdraon *et al*., 2019b). The interactions between members of the microbial communities associated with plant residues are a potential lever for innovative disease management strategies involving microbiome-based biocontrol.

In recent years, our knowledge of the diversity of microbial communities has increased considerably, through the development of whole-community metagenomic sequencing methods, which have fuelled many studies on the microbial ecology of the phyllosphere (Yang *et al*., 2003), rhizosphere (Øvreås *et al*., 1998; Zhang *et al*., 2021), and, more recently, the ‘residue sphere’ (Kerdraon *et al*., 2019b). However, these ecological studies cannot completely dispense with culture-dependent approaches. The isolation and culture of fungal and bacterial species are indispensable as they complement taxonomic databases through a single-strain approach (from isolation to molecular characterization), making it possible to test whether the interactions between micro-organisms predicted by metagenomic sequencing (e.g. through the analysis of interaction networks; Fath *et al*., 2007) actually occur. They are also essential for assessing the efficacy of strains as potential biocontrol agents (e.g. Daniel *et al*., 2004; Comby *et al*., 2017; Tuan Hamzah *et al*., 2018), with such tests often limited by the availability of representative microbial culture collections (Lewis *et al*., 2020).

A major bias is introduced if only the cultivable fraction is considered, because this precludes an exhaustive description of the microbial community (Wintzingerode *et al*., 1997; Armanhi *et al*. 2016). For example, the fraction of cultivable bacteria in the phyllosphere may be much lower than that in other environments (Müller and Ruppel, 2014; Ritz, 2007), depending on the media used, the growth conditions, and the plant species (e.g. Rastogi *et al*., 2010; Stiefel *et al*., 2013). It has also long been thought that only a very small fraction of soil micro-organisms — about 1% — can be isolated *in vitro* (Daniel *et al*., 2004; Ritz, 2007). However, recent advances in culture techniques have led to suggestions that this “1% paradigm” may no longer be correct (Martiny, 2019), particularly for leaf- and root-derived bacteria (e.g. in *Arabidopsis thaliana,* the overlap between culture-dependent community profiles and those obtained by metagenomic sequencing is over 50%; Bai *et al*., 2015). To our knowledge, no equivalent information is available for the ‘residue sphere’.

The combination of metabarcoding and culture techniques is thought to provide a more complete view of the microbiota. Direct comparisons have been made between the microbial population compositions obtained by culture-dependent (with or without single-strain molecular identification) and culture-independent (whole-community metagenomic or metabarcoding sequencing) methods, mostly for bacterial communities, in the contexts of human health (gut microflora; e.g. Abayasekara *et al*., 2017; Hilton *et al*., 2016; Zapka *et al*., 2017), food (cheese; Carraro *et al*., 2011), and plants (roots, leaves, and seeds; Armahni *et al*., 2013; 2016; 2018; Jackson *et al*., 2013). Comparative studies focusing on the fungal fraction (e.g. the endophytic flora; Ben Chobba *et al*., 2013; Comby *et al*., 2016; Dissanayake *et al*., 2018), or even on both the bacterial and fungal fractions (e.g. root symbionts; Thiergart *et al*. 2019) are much rarer. Crop residues, in particular, are very much the “poor relation” in analyses of this type.

The structure of the microbiota associated with arable crop residues has recently been characterized by metabarcode sequencing in a wheat-oilseed rape rotation system (Kerdraon *et al*., 2019b). The impact of plant species, seasonality and crop rotation on the residue microbiome was highlighted for both fungal and bacterial communities, with the replacement of plant-specific taxa — including pathogens and endophytes — by more generalist taxa originating from the soil. The effect of the presence of *Zymoseptoria tritici* and *Leptosphaeria maculans* — two important residue-borne fungal pathogens of wheat and oilseed rape, respectively (Suffert and Sache, 2011; Fitt *et al*., 2006) — on the residue microbiome was then assessed by combining linear discriminant analyses and ecological network analyses (Kerdraon *et al*., 2019c; 2020). The bacterial and fungal communities associated with residues were compared between wheat with and without preliminary *Z. tritici* inoculation, between wheat in contact with the soil and without soil contact (Kerdraon *et al*., 2019c) and between two isogenic oilseed rape lines with and without a resistance gene against *L. maculans* (Kerdraon *et al*., 2020). Correlative approaches of this type provide original information about the impact of phytopathogenic microorganisms on a whole microbial community, and may facilitate the identification of beneficial keystone taxa and assemblages beneficial to host plant health. Indeed, several species already described as biocontrol agents were found to be affected by the presence of the two pathogens studied, consistent with recent results for *Fusarium* sp. on maize residues (Cobo-Diaz *et al*., 2019b; 2019c). However, it is necessary to go beyond correlative approaches, by using methods for determining causal relationships through the testing of direct interactions. Such approaches require the prior isolation, typing and storage of representative microbial strains, before their impact on the development of plant pathogens can be tested, using single species or more complex microbial assemblages, involving the reintroduction of some of the complexity of the natural system through the creation of ‘synthetic communities’, for example (Großkopf and Soyer, 2014).

Assessments of the effectiveness of culture-dependent and culture-independent approaches, in terms of microbial representativeness, are essential to highlight the key role of crop residues in agrosystems. In this study, we investigated the diversity of fungal and bacterial communities in wheat and oilseed rape residues, with these two types of approach. We compared the ability of these two approaches — whole-community metabarcode sequencing and the differential isolation of fungal and bacterial strains followed by molecular identification — to describe the microbial diversity of this habitat.

## Materials and methods

### Overall strategy

Wheat (W) and oilseed rape (O) residues were collected from three field plots after the first two cropping seasons (2015-2016 and 2016-2017) of a three-year wheat-oilseed rape rotation at the Grignon experimental station (Yvelines, France; 48° 51’N, 1° 58’ E; Kerdraon *et al*., 2019b). Each of the residues sampled was cut in two lengthwise, with one half used for metabarcoding and the other half for microbial isolation (**Fig. 1**). Whole-community metabarcoding (*sensu stricto* high-throughput amplicon sequencing) was implemented identically for the residues of both crops, targeting the gene encoding the 16S rRNA for bacteria and the ITS1 region for fungi, respectively, as described by Kerdraon *et al*. (2019b). For the analysis of microbial diversity by a culture-dependent approach, bacterial and fungal strains were treated separately, according to two strategies: for bacteria, reduced substrate sampling was coupled with high-throughput isolation, whereas, for fungi, broad substrate sampling was coupled with low-throughput isolation. The individual strains were identified molecularly, by barcoding targeting the v4 region of the gene encoding 16S rRNA in a single MiSeq Illumina run for bacteria, and Sanger sequencing of the ITS1-5.8S-ITS2 region with ITS1F-ITS4 primers for fungi (**Fig. 2**).

**Figure 1.**
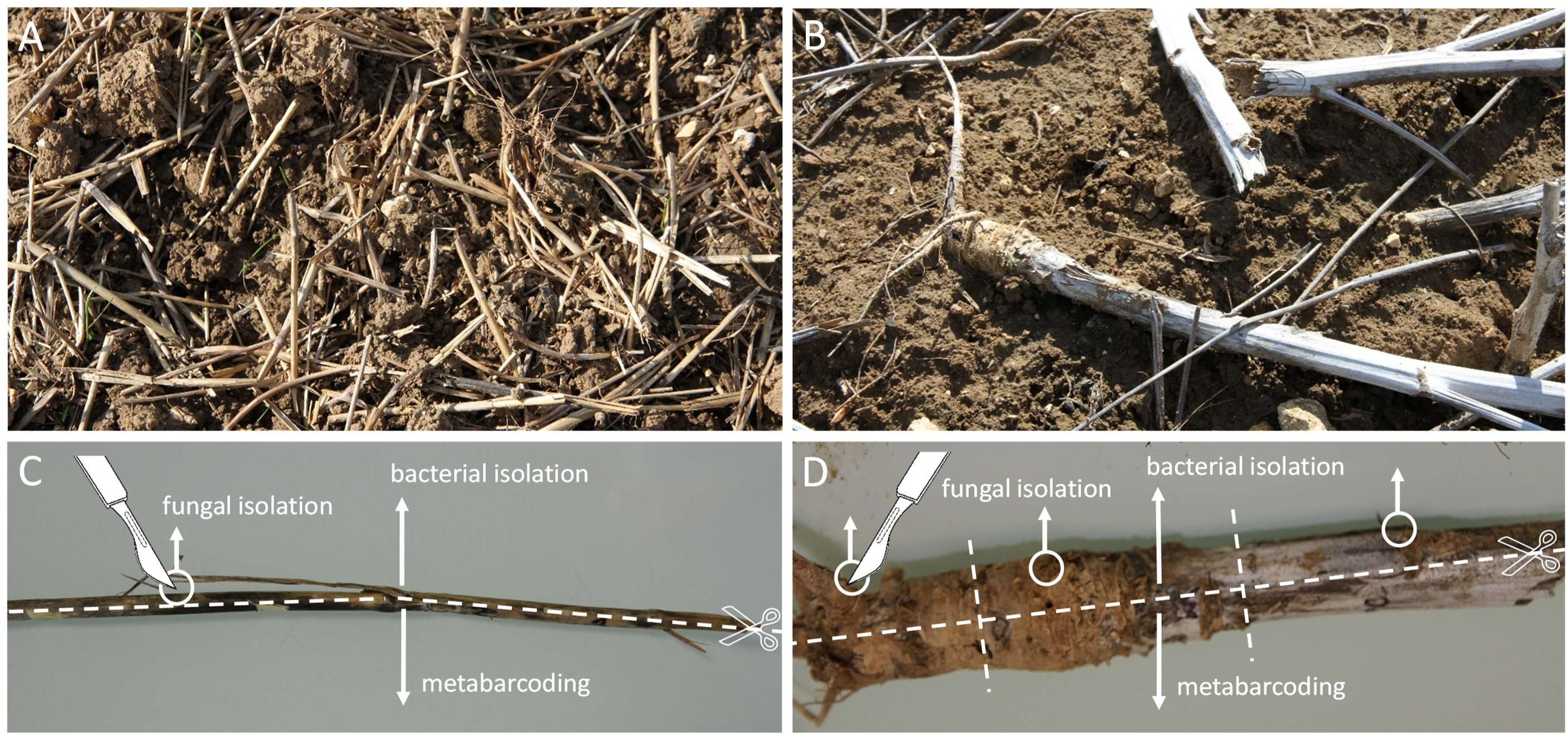
Sampling and preparation of crop residues. Wheat (**A**) and oilseed rape residues (**B**) on the ground in field plots. Wheat (**C**) and oilseed rape residues (**D**) cut lengthwise and selection of pieces of residue. For fungal isolation, we used one 1 mm^2^ piece of residue cut from the first half section of each of 12 wheat residues and 3 × 1 mm^2^ pieces of residue cut from the first half section of each of four oilseed rape residues (one piece from the stem base, one from the start of the pivotal root, and one piece from the collar); for bacterial isolation, the rest of the first half section of each residue was used for maceration, if required. All the second half sections of the residues were used for metabarcoding (Kerdraon *et al*., 2019b).

**Figure 2.**
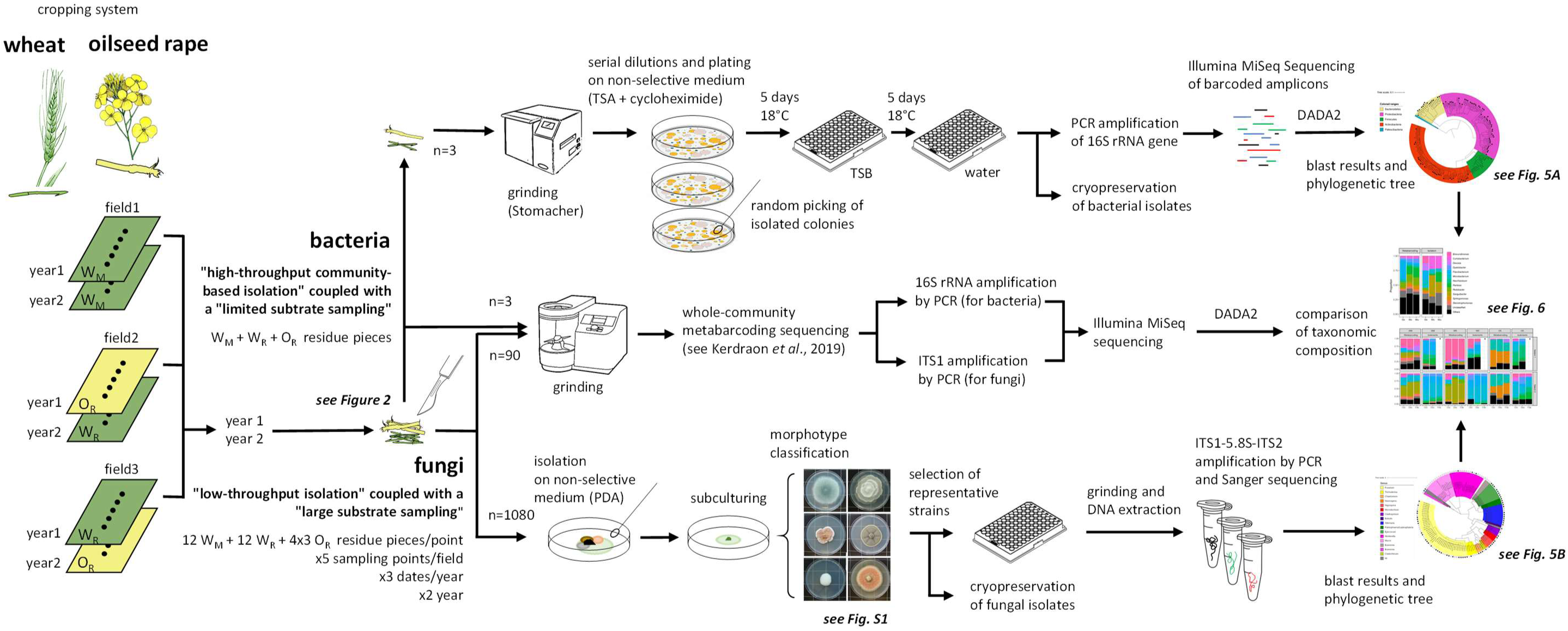
Overview of the culture-dependent molecular identification and whole-community metabarcoding strategies for characterizing and comparing the bacterial and fungal communities of wheat (W) and oilseed rape (O) residues.

### Culture-dependent molecular identification of residue-associated bacteria

#### Residue sampling

For bacteria, we focused on recovering a large number of bacterial strains from a small number of residue samples (*n*=3) after the first cropping season: two sets of five pieces of wheat residue from a wheat monoculture plot (W_M_) and from a first plot under the wheat-rapeseed rotation (W_R_), and a set of five pieces of oilseed rape residue O_R_ (from a second plot under the wheat-oilseed rape rotation) (**Fig. 1 and 2**). Crop residue-associated bacteria were isolated by spreading the macerates obtained with each set of residue pieces onto a primary plate and picking bacterial colonies at random.

#### Isolation of the bacterial fraction

The pieces of residue (each weighing approximately 5 g) were soaked in 2 mL of phosphate-buffered saline (PBS, Sigma-Aldrich P4417) supplemented with Tween 20 (0.05% v/v) per gram of fresh material. Residues were crushed and homogenized with a Stomacher blender (Mixwel, Alliance Bio Expertise) for two minutes. Serial dilutions of the resulting suspensions were plated on 1/10 strength tryptic soy agar (TSA; 17 g.L^-1^ tryptone, 3 g.L^-1^ soybean peptone, 2.5 g.L^-1^ glucose, 5 g.L^-1^ NaCl, 5 g.L^-1^ K_2_HPO_4_, and 15 g.L^-1^ agar) supplemented with cycloheximide (50 µg.mL^-1^). The plates were incubated for five days at 18°C, and 2232 colonies (1395 for W_M_, 744 for W_R_ and 93 for O_R_) were picked (from two plates for each set of residue pieces) and subcultured in 96-well plates containing 200 µL of tryptic soy broth (TSB) per well. One negative control (TSB with no added colony, as a control for sterility), one positive control (*Agrobacterium tumefaciens* CFBP 2413, control for growth and barcoding) and a water control (negative control for subsequent PCR assays) were included on all plates. The plates were incubated for five days at 18°C with continual shaking (70 rpm), and we then transferred 10 µL of bacterial suspension to 90 µL of sterile water for PCR amplification. We added 190 µL of 80% glycerol to the remaining bacterial suspension in each well, which was then stored at −80°C.

#### High-throughput partial sequencing of the 16S rRNA gene

Molecular typing was performed on each bacterial isolate with the 515f/806r primer pair (5’-GTGCCAGCMGCCGCGGTAA and 5’-GGACTACHVHHHTWTCTAAT; Caporaso *et al*., 2011) targeting the v4 region of 16S rRNA gene, according to the procedure described by Armanhi *et al*. (2016). Briefly, each primer was labeled with a unique plate barcode. PCR amplification was performed in 25 µL reaction mixtures containing 2.5 µL boiled bacterial suspension (99°C, 5 min), 0.25 µL GoTaq flexi DNA Polymerase (Promega), 5 µL Greenflexi buffer (Promega), 2 µL dNTP (Promega U151B, 2.5 mM), 1.5 µL MgCl_2_ (25 mM), 0.5 µL of each primer (10 mM) and 12.75 µL water. The following conditions were used for amplification: 35 cycles of amplification at 94°C (30 s), 50°C (45 s) and 68°C (90 s), followed by a final elongation at 68°C for 10 min. Amplicons were purified with magnetic beads (Sera-Mag^TM^, Merck) at a beads-to-PCR product ratio of 1.2. Amplicons with plate barcodes from this first PCR were pooled on a single plate and a second PCR amplification was performed to incorporate Illumina adapters and unique well barcodes. PCR was performed with 10 µL Greenflexi buffer, 2 µL dNTP, 4 µL MgCl_2_, 0.2 µL GoTaq flexi DNA polymerase, 26.8 µL water, 2 µL of each primer and 5 µL purified PCR1 product. PCR conditions were as follows: initial denaturation at 94°C (1 min), followed by 12 cycles of 94°C (1 min), 55°C (1 min) and 68°C (1 min), and a final elongation at 68°C for 10 minutes. PCR products were purified with Sera-Mag^TM^, pooled in a single tube with 10% PhiX and sequenced with MiSeq reagent kit v2 (500 cycles).

#### Sequence analyses

Fastq files were demultiplexed and plate-specific primers were trimmed with cutadapt version 1.8 (Martin, 2011). Fastq files were processed with DADA2 1.6.0 (Callahan *et al*., 2016), using the following parameters: truncLen=c(180,120), maxN=0, maxEE=c(1,1), truncQ=5. Chimeric sequences were identified and removed with the removeBimeraDenovo function of DADA2. The taxonomic affiliations of amplicon sequence variants (ASVs) were determined with a naive Bayesian classifier (Wang *et al*., 2007) implemented in DADA2. ASVs derived from the 16S rRNA gene were classified with the Silva 132 taxonomic training data (silva_nr_v132_train_set.fa.gz). ASVs that were unclassified at phylum level or affiliated to chloroplasts and mitochondria were removed. Sequence contaminants were identified with decontam package version 1.4 (Davis *et al*., 2018), using the “prevalence” method at a threshold of 0.01. Finally, ASVs were filtered on the basis of their relative abundances per well. Only ASVs with a relative abundance greater than 1% per well were retained. A neighbor-joining phylogenetic tree was built with the phangorn R package v 2.5.5 (Schliep, 2011), using a multiple sequence alignment obtained with the DECIPHER R package v. 2.12.0 (Wright, 2015).

### Culture-dependent molecular identification of residue-associated fungi

#### Residue sampling

Fungi were isolated by selecting a small number of fungal colonies from a large number of crop residue samples (*n*=1080) collected during three periods (October, December and February) of the second and third cropping seasons: 12 wheat (W) and 4 oilseed rape (O) residue samples collected at five sampling points in the wheat monoculture plot (W_M_) and in the two wheat-oilseed rape rotation plots (W_R_ and O_R_). Several 1 mm^2^ pieces of the wheat and oilseed rape residue samples were removed with a scalpel for fungal isolation (**Figs. 1 and 2**).

#### Isolation of the fungal fraction

Each piece of residue was deposited on a PDA (potato dextrose agar, 39 gL^−1^) plate and kept at 18°C in the dark for 5-8 days. Dominant and non-dominant strains were purified by successive subcultures on PDA. Only one fungal morphotype was chosen per sample, with the aim of collecting 20 to 25 strains per sampling point × period combination × plot. Fungal strains were kept at −80°C in glycerol-water solution (20% v/v) for long-term storage.

#### Molecular characterization of the fungal collection by Sanger sequencing

The strains stored at −80°C were subcultured on PDA plates for molecular characterization. The mycelium of each plate was scraped off and placed in the well of a 96-well plate. The mycelia were then lyophilized and ground with a tungsten bead in 50 μL lysis buffer (Qiagen DNeasy-Plant AP1 buffer) with a Retsch™-MM400 (Retsch, France) mill operating at 20 Hz for 30 seconds. The DNA was extracted with the DNeasy Plant Mini DNA extraction kit (Qiagen, France), according to the simplified instructions provided by the manufacturer, as the DNA columns were not used. The DNA of all strains was quantified with a NanoDrop ND-1000 UV-Vis spectrophotometer (NanoDrop, Wilmington, DE, USA), adjusted to 10 to 20 ng.mL^-1^ and stored at −20°C until use.

For molecular characterization, a fragment encompassing the ITS regions (ITS1 and ITS2) and the small 5.8S subunit of rDNA was amplified from each DNA with the ITS1F (Gardes *et al*., 1993) and ITS4 (White *et al*., 1993) primers. The PCR mixture contained 1x master mix for the Type-it Microsatellite PCR Kit (Qiagen, France), 3 µL of Q-Solution and 0.25 µM of each primer in a total volume of 30 µL. PCR was performed with an Applied Biosystems 9700 thermal cycler. The amplification parameters were as follows: initial denaturation at 95°C for 5 min, 35 cycles of 95°C for 60 s, 60°C for 30 s, and 72°C for 60 s, and a final elongation at 72°C for 10 min. We checked the quality of the PCR products by electrophoresis in 2% agarose gels. The PCR products were then subjected to single-strand sequencing by the Sanger method (Eurofins, France). Chromatograms were checked with DNASTAR lasergene sequence analysis software (Burland, 2000) to generate the sequences used for molecular identification. The taxonomic assignment of each sequence was performed with a naïve Bayes classifier on the UNITE7.1 database, as for the culture-independent approach (metabarcoding).

#### Phylogenetic tree for the ITS region of fungal strains

The ITS sequences of each of the 424 fungal strains were blasted against each other to prevent redundancy in subsequent analyses. We obtained 131 unique sequences, which were considered to be ‘representative ASVs’ of the fungal collection. A phylogenetic tree of these ASVs was built with the ‘build’ function of ETE3 v3.1.1 (Huerta-Cepas *et al*., 2016), as implemented on GenomeNet (https://www.genome.jp/tools/ete/). An alignment was generated with MAFFT v6.861b, using the default options (Katoh and Standley, 2013). The tree was constructed with FastTree v2.1.8, using the default parameters (Price *et al*., 2009). The values at the nodes were SH-like local support. Data were visualized with Interactive Tree Of Life (iTOL) v4 (Letunic and Bork, 2019).

## Results

### Principal results for bacteria

#### Filtering collection data

The analysis of the cultivable bacterial fraction was based on 2232 bacterial colonies (1395 for W_M_, 744 for W_R_ and 93 for O_R_) obtained after maceration of the three sets of residues. Sequencing was successful for 2201 wells, with a median of 3709 reads per well (min 27, max 18774; **Fig. 3A**), and the detection of 753 ASVs. The number of ASVs per well ranged from 1 to 19, with a median value of 2. The high diversity of ASVs in each well often resulted from the presence of sequences presenting only one single nucleotide polymorphisms (SNPs). For instance, two ASVs were systematically detected in the wells corresponding the positive PCR control (*A. tumefaciens* CFBP 2413). However, these two ASVs had different relative abundances. The highly abundant ASV (>95% of the reads) was 100% identical to the genomic sequence of *A. tumefaciens* CFBP 2413, and the lower abundance ASV (<5% reads) presented one SNP relative to the genomic sequence. The observed ASV diversity within each well therefore probably reflects amplification and/or sequencing artifacts. We corrected for these errors by applying a conservative filter to the read set retained, in which we retained only ASVs with a relative abundance exceeding 5% in the well concerned. This filtering procedure reduced the number of ASVs to 158, with a median of one ASV per well (1-5 ASVs; **Fig. 3B**).

**Figure 3.**
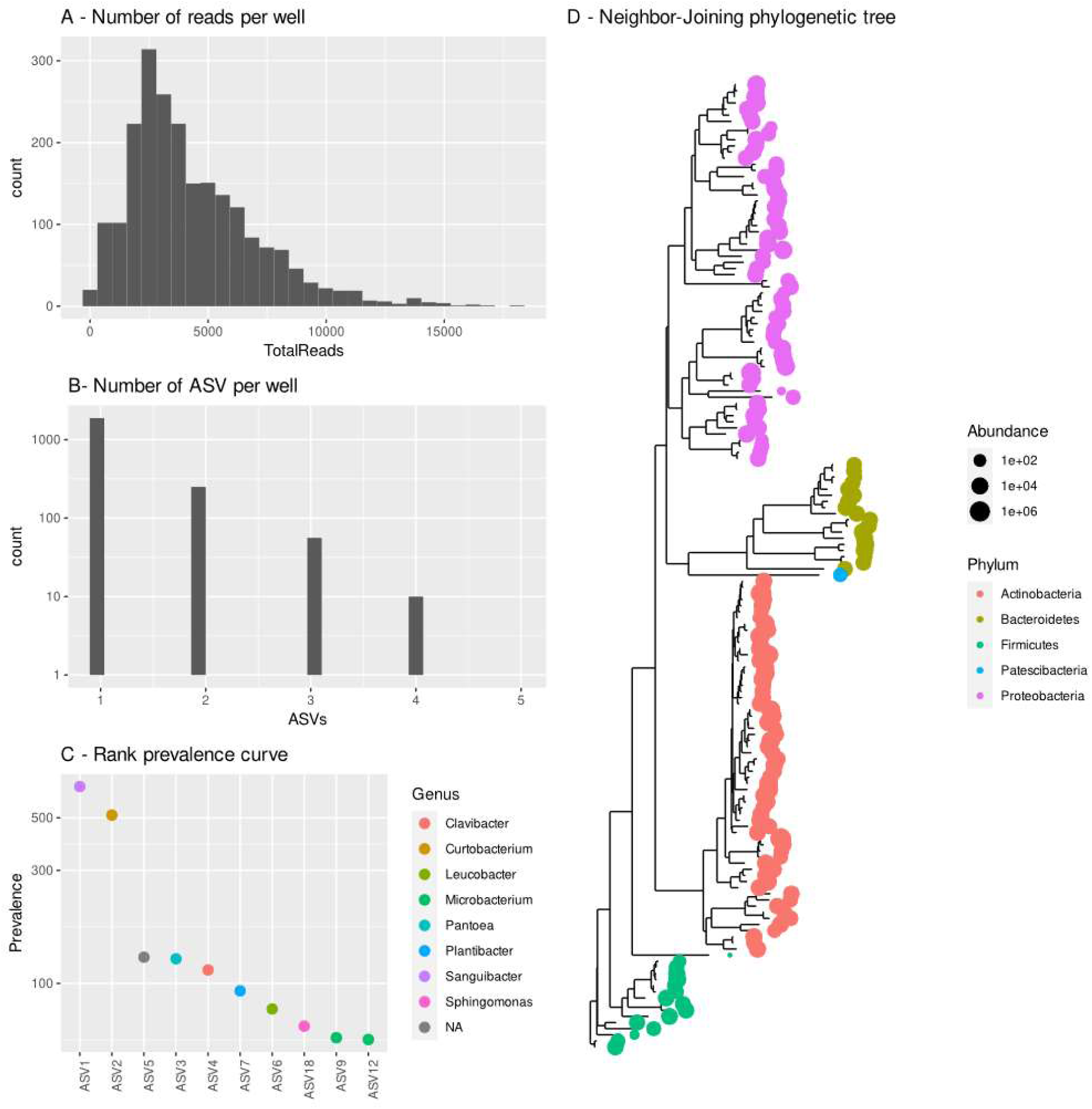
Community-based culture collection of bacterial strains isolated from three wheat and oilseed rape residue samples. Distribution of sequences (**A**) and ASVs (**B**) per well after abundance filtering (>5%). Prevalence of the 10 most frequent bacterial ASVs isolated in the community-based culture collection (**C**). Neighbor-joining phylogenetic tree based on the v4 region of the 16S rRNA gene of all bacterial strains found in the community-based culture collection (**D**).

#### Description of the collection

These 158 ASVs were affiliated to five phyla (Actinobacteria, Bacteroidetes, Firmicutes, Patescibacteria and Proteobacteria) and 36 bacterial genera (**Fig. 3D** and *Table S1*). The most prevalent bacterial strains recovered with the culture-dependent approach were related to *Sanguibacter* (*n*=677) and *Curtobacterium* (*n*=519), two genera from Actinobacteria (**Fig. 3C**).

#### Comparison of metabarcoding and culture-dependent molecular identification methods

The metabarcoding analysis (Kerdraon *et al*., 2019b) identified 1129 ASVs corresponding to 16 phyla and 231 bacterial genera. The most prevalent ASVs were affiliated to the phyla Actinobacteria, Bacteroidetes and Proteobacteria (**Fig. 4A**) and the genera *Sphingomonas* and *Neorhizobium* (**Fig. 4B**). The culture-dependent approach resulted in the recovery of 158 ASVs after filtering, 89 of which were 100% identical to ASVs detected with metabarcoding (**Fig. 5** and *Table S1*). The most abundant taxa identified by metabarcoding were successfully isolated, because there were representative strains in the community-based culture collection for eight of the 10 most abundant taxa (**Fig. 4B**). Most of the collected strains not previously detected by metabarcoding belonged to Firmicutes or Actinobacteria (**Fig. 5A**). The culture collection was particularly enriched in members of the families Microbacteriaceae and Paenibacillaceae (*Table S1*). Finally, we compared the overall profiles of residue-associated bacterial communities characterized by metabarcoding sequencing and by culture-dependent molecular identification, focusing on the 12 most abundant bacterial taxonomic groups. Marked differences were noted in the proportions of certain genera: ASVs affiliated to Actinobacteria (*Microbacterium*, *Curtobacterium*, *Sanguibacter*) were more frequently represented in the culture-dependent molecular identification data than in the metabarcoding data (**Fig. 6A**). Conversely, ASVs affiliated to the genera *Dyadobacter*, *Pedobacter* and *Flavobacteria* were more frequently represented in the metabarcoding data. Nevertheless, several differences in profiles were detected between wheat (W_M_ and W_R_) and oilseed rape (O_R_) residues with both approaches, highlighting their consistency. For example, ASVs affiliated to *Flavobacterium* and *Devosia* were less frequently represented on wheat residues than in oilseed rape residues, whereas ASVs affiliated to *Curtobacterium* were almost entirely absent from oilseed rape residues.

**Figure 4.**
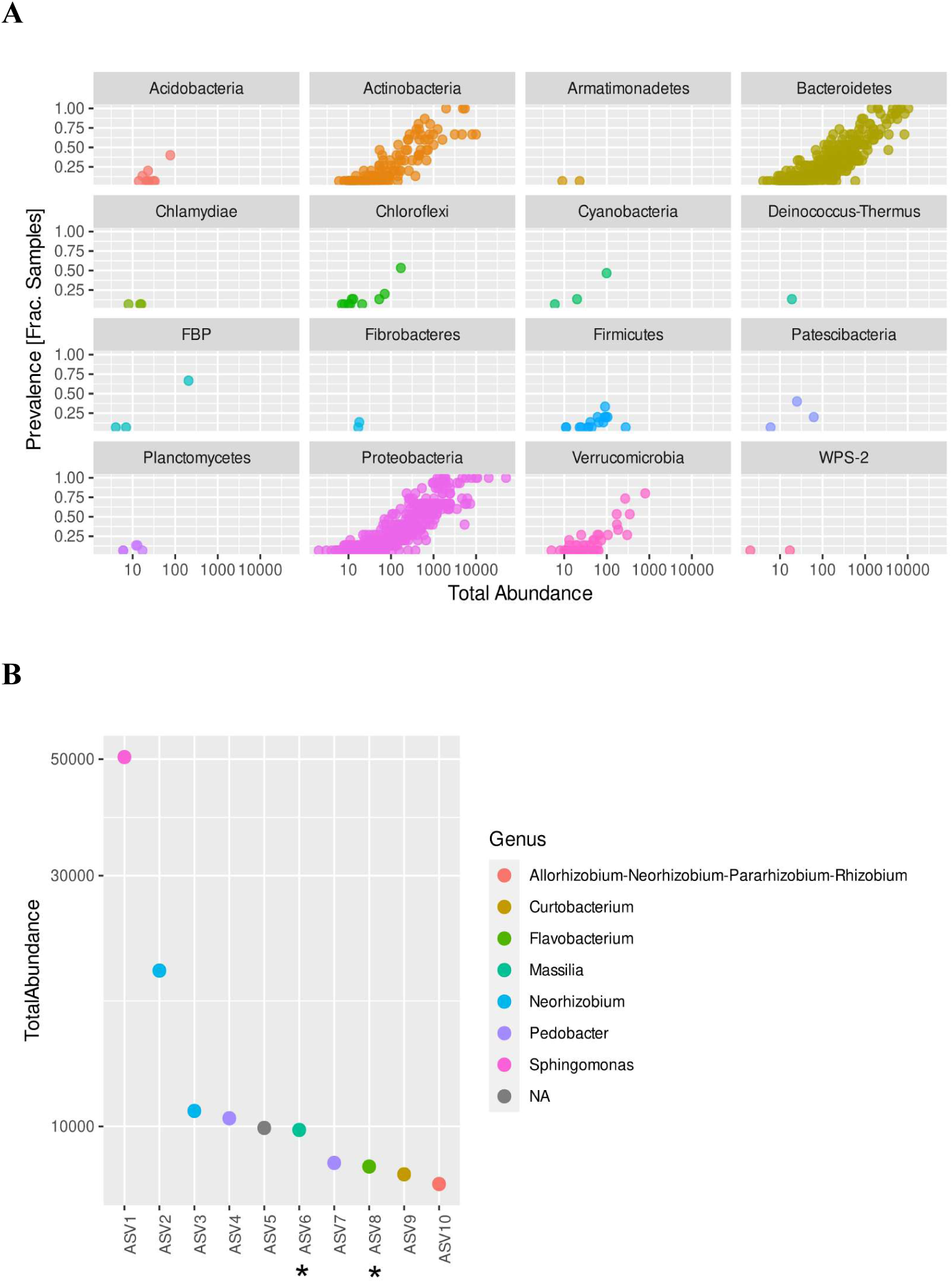
Analysis of the bacterial diversity of three wheat and oilseed rape residue samples characterized by metabarcoding (data from Kerdraon et al., 2019b). (**A**) Abundance and prevalence of the 1129 ASVs detected by metabarcoding. (**B**) Rank abundance curve of the 10 most abundant ASVs identified by metabarcoding. Asterisks indicate that no representative strains were isolated in the community-based culture collection.

**Figure 5.**
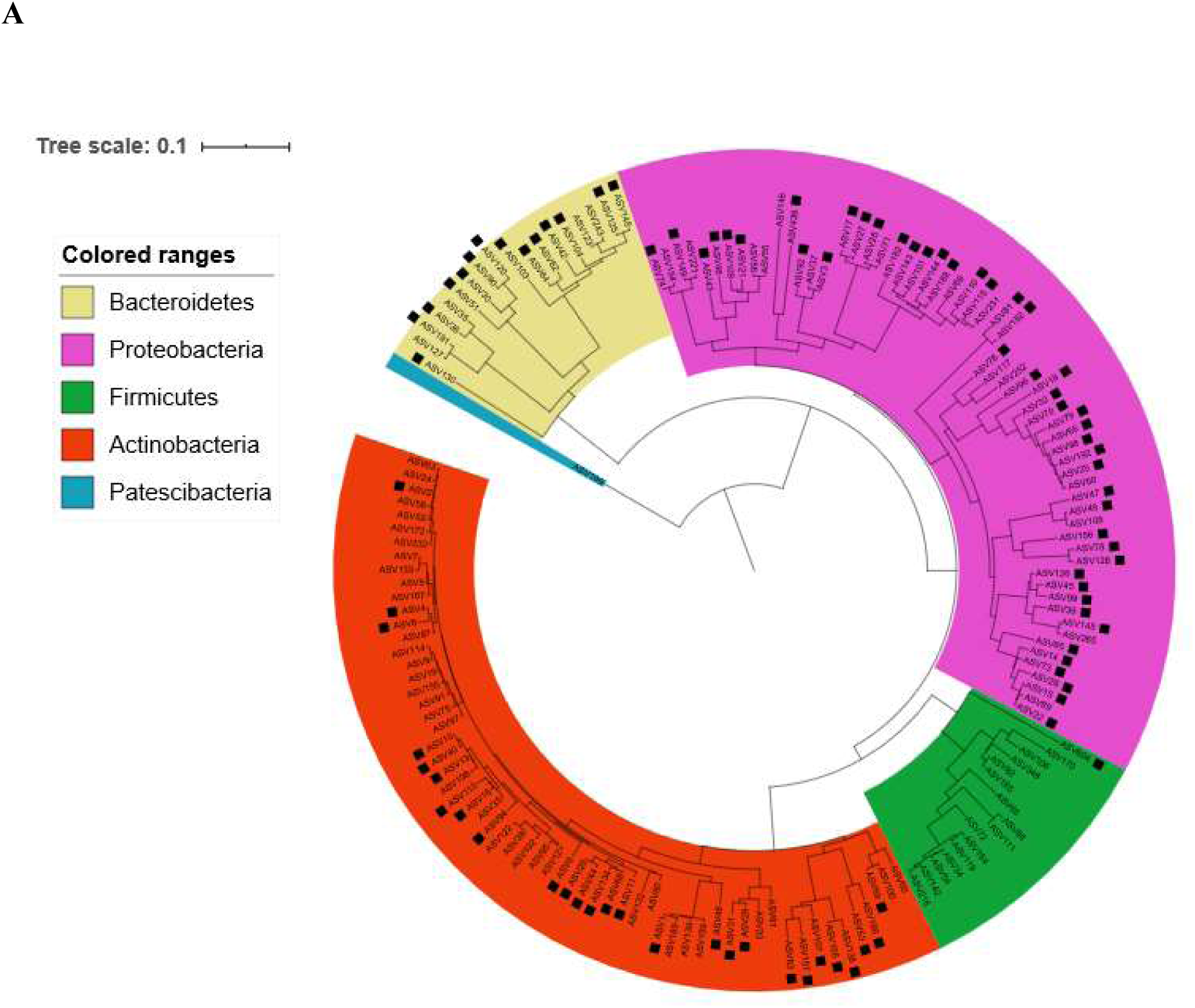
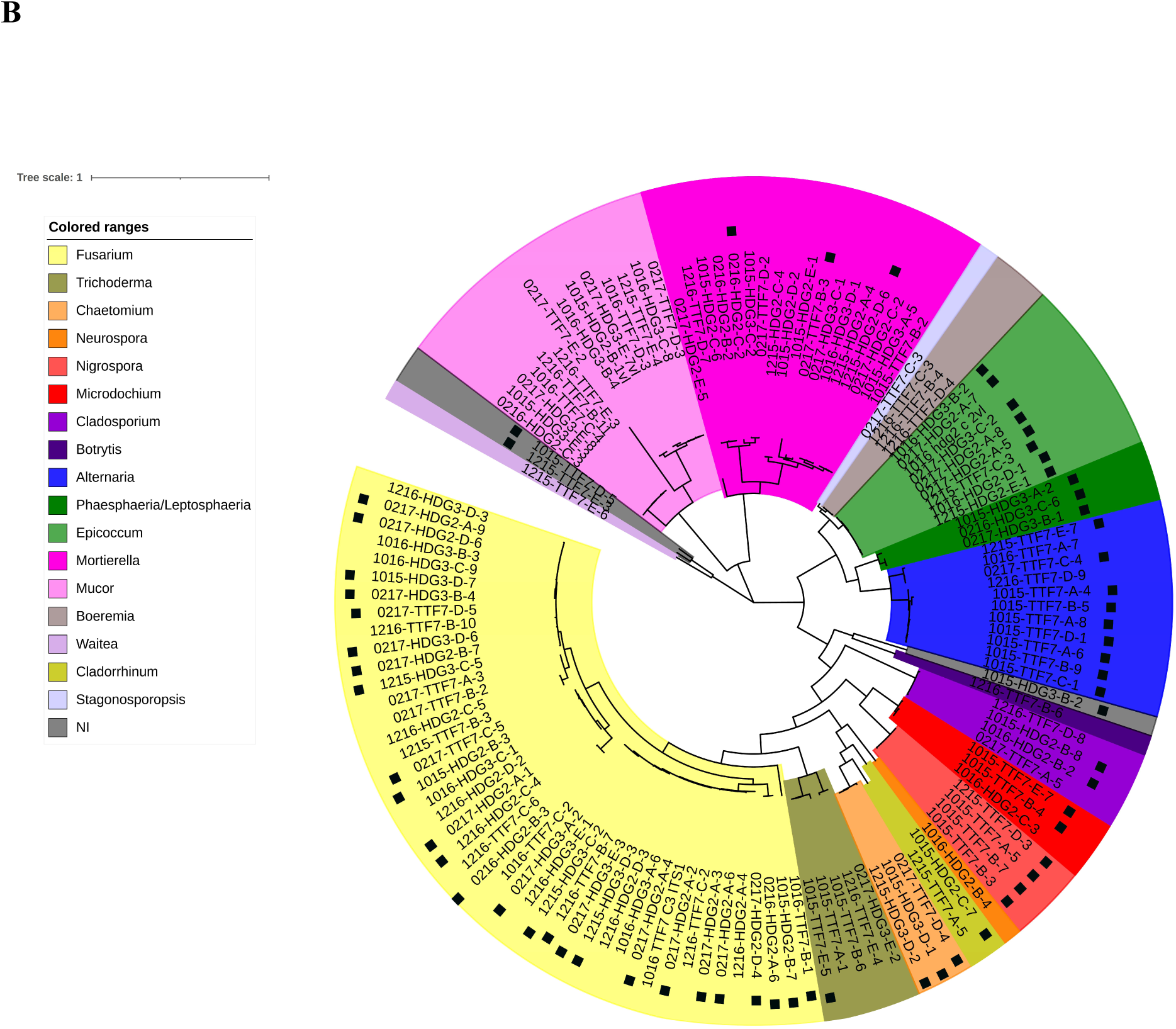
Phylogenetic distribution of the bacterial strains (from three residue samples) and fungal strains (from 18 residue samples) recovered through culture collection. A Neighbor-joining phylogenetic tree was constructed (**A**) with the v4 region of 16S rRNA gene of all unique bacterial ASVs (*n*=158) and (**B**) the ITS1-5.8S-ITS2 region of all unique fungal ASVs (*n*=131) detected in each culture collection. The presence of an ASV displaying 100% identity to the ITS1 region in the metabarcoding dataset (Kerdraon *et al*., 2019b) is indicated by a black square.

**Figure 6.**
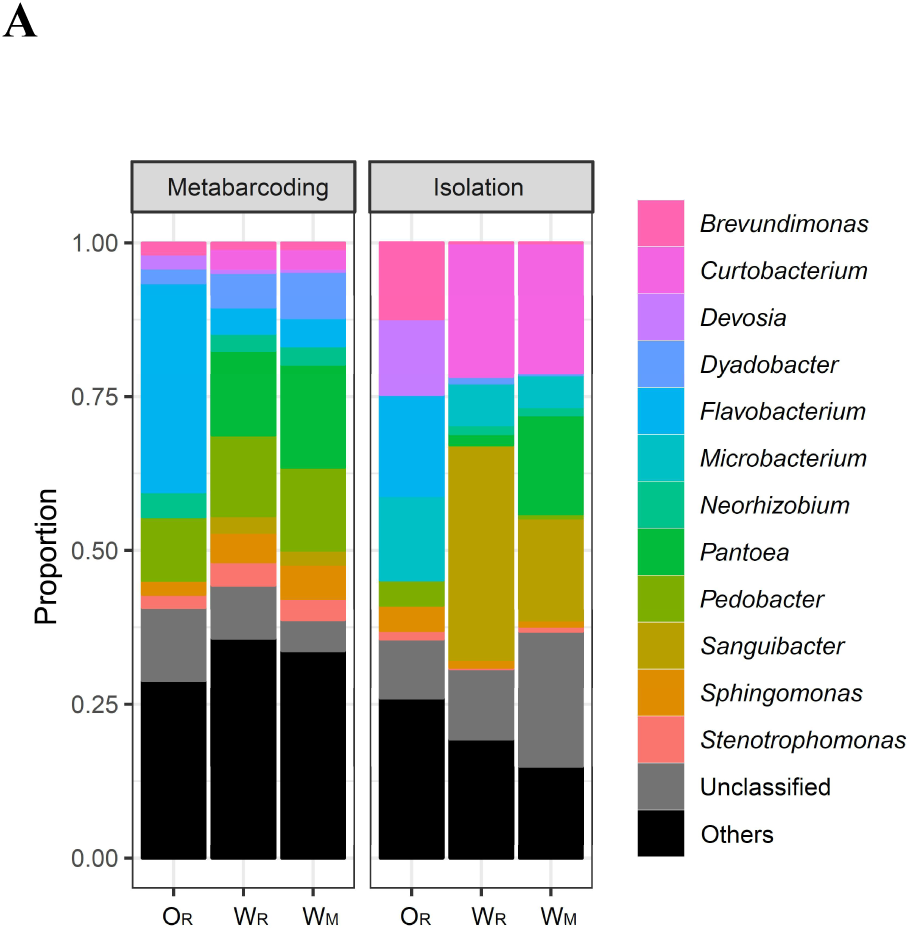
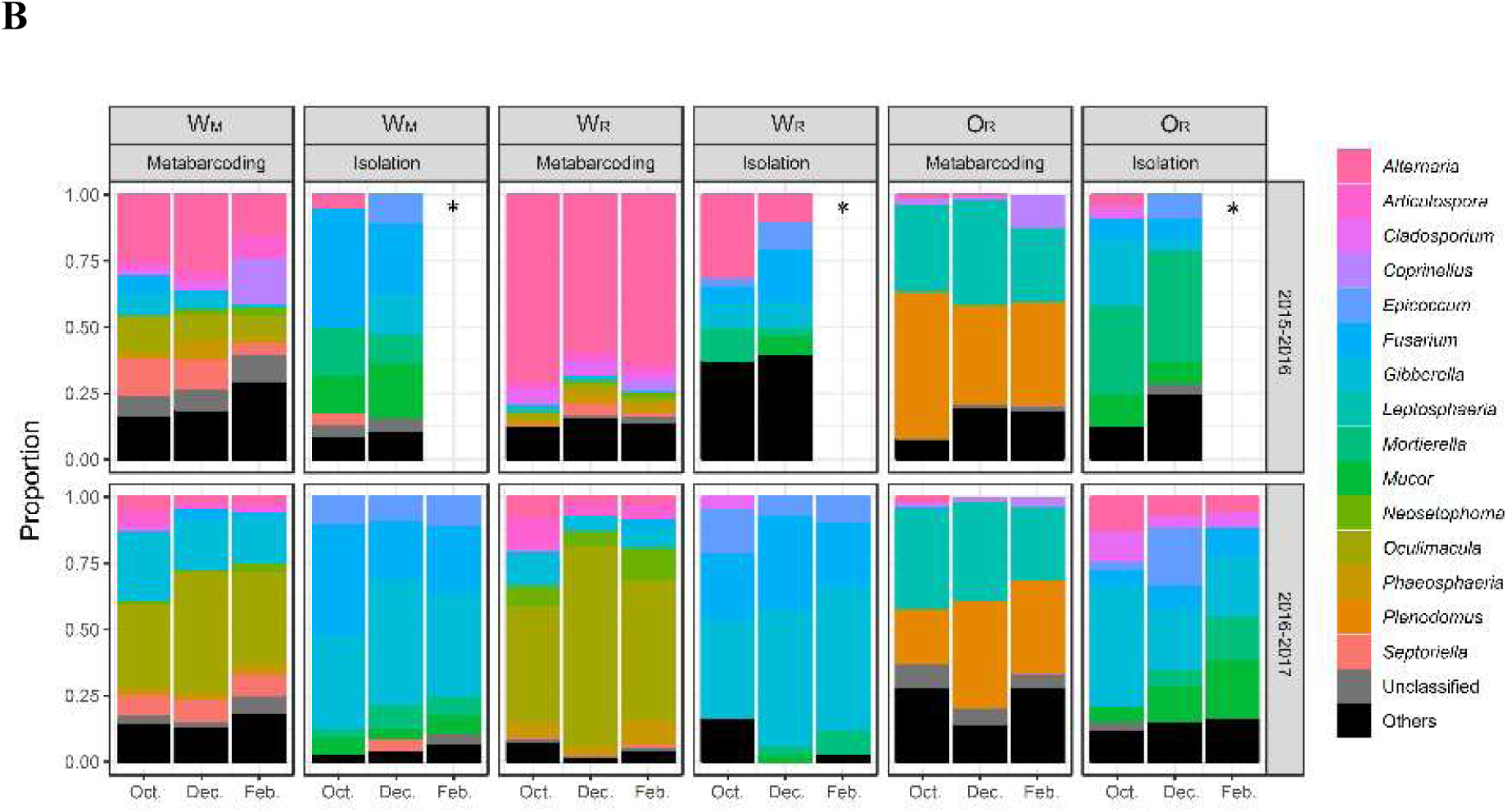
Comparison of the overall profiles of the residue-associated bacterial and fungal communities obtained by metabarcode sequencing (‘metabarcoding’) and by culture-dependent molecular identification (‘isolation’). (**A**) For the bacterial community, comparisons were based on the same three wheat (W) and oilseed rape (O) residue samples collected from the wheat monoculture plot (W_M_) and the two wheat-oilseed rape rotation plots (W_R_ and O_R_) in October 2015. (**B**) For the fungal community, comparisons were based on the same 18 wheat (W) and oilseed rape (O) residue samples collected from the wheat monoculture plot (W_M_) and the two wheat-oilseed rape rotation plots (W_R_ and O_R_) during three sampling periods (October, December, February) in the 2015-2016 and 2016-2017 cropping seasons. Only the 12 most abundant bacterial genera and the 16 most abundant fungal genera are presented. The composition of the three sets marked by * is missing due to unsuccessful isolation.

### Principal results for fungi

#### Filtering of the collection data

The analysis of the cultivable fungal fraction led to the isolation and sequencing of 424 fungal strains; morphotypes were described from photographs for 171 of these strains (*Fig. S1*) and full ITS regions (ITS1-5.8S-ITS2) were sequenced to assess the taxonomic diversity of the collection (*Table S2*). A dataset of 131 unique ASVs was obtained by blasting the 424 sequences against each other.

#### Description of the collection

The 131 ASVs were affiliated to 17 genera, with some left unclassified (**Table S2, Fig. 5B**). The most prevalent strains isolated were affiliated to the genus *Fusarium* (50% of the strains, 37% of the ASVs).

#### Comparison of metabarcoding and culture-dependent molecular identification

The metabarcoding analysis identified 894 ASVs belonging to 148 fungal genera (Kerdraon *et al*., 2019b). The microbial community profiles obtained by metabarcoding and culture-dependent molecular identification were quite different (**Fig. 6**). However, the two approaches clearly distinguished between the fungal communities of the three residue samples (W_M_, W_R_ and O_R_) and between the two cropping seasons, consistent with the findings of Kerdraon *et al*. (2019b). The ASVs corresponding to the most frequently isolated fungal genera were also detected by metabarcoding, except for the ASVs affiliated to the genera *Mucor* and *Mortierella*, represented by 34 and 46 strains, respectively (*Table S3*); this result is notable because it was unexpected. Strains from some other genera, such as *Actinomucor*, *Boeremia*, *Botrytis*, *Neurospora*, and *Stagonosporopsis,* were isolated at a very low frequency and were not detected at all by metabarcoding. Several other genera, such as *Epicoccum* and *Fusarium* (known to be highly competitive *in vitro* due to their high growth rates relative to other fungi) were, unsurprisingly, overrepresented in analyses based on isolation relative to those based on metabarcoding, particularly for wheat residues (W_M_ and W_R_) during the second cropping season (2016-2017) (**Fig. 6**). Conversely, the ASVs affiliated to several of the most frequent fungal genera identified by metabarcoding were obtained rarely, if at all, by isolation techniques; this was the case for *Leptosphaeria* and *Plenodomus* (probably *L. maculans* and *P. biglobosus*, known to be pathogens of oilseed rape; Kerdraon *et al*., 2020) in O_R_ and *Oculimacula* (probably *O. yallundae,* known to be pathogen of wheat that survives on straw; Vera and Murray, 2016) in W_M_ and W_R_.

For technical reasons, the comparison of the results obtained by the two approaches was less straightforward than for bacteria, and fully comparable data would have been obtained had ITS1 been extracted alone from the long fragment. Indeed, the size of the genomic region amplified and sequenced for fungal strain identification in the culture-dependent approach (ITS1-5.8S-ITS2 region) was larger and included the genomic region targeted in the culture-independent approach (ITS1 region only). Thus, the ITS1 region of one ASV obtained by metabarcoding could potentially display 100% identity with the IST1 of the sequenced ITS1-5.8S-ITS2 region of several isolated strains. Overall, 33 of the 894 ASVs identified by metabarcoding were 100% identical to one or several ASVs from the dataset of the collection (**Fig. 5B**). As an example of equivocal matches, one ASV detected by metabarcoding displayed 100% identity to eight ASVs of the genus *Epicoccum* from the collection. Conversely, several ASVs detected by metabarcoding identified displayed 100% identity to one ASV from one genus (e.g. *Alternaria* and *Fusarium*). The culture-dependent approach is complementary to metabarcoding, as three unidentified ASVs of the metabarcoding dataset were 100% identical to three ASVs corresponding to three strains from the collection.

## Discussion

### Complementarity of the two approaches

This study was the first to compare culture-dependent and culture-independent approaches for characterizing the microbiota of arable crop residues, using two different isolation strategies to generate community-based culture collections adapted to the diversity of the bacterial and fungal kingdoms. The strategy developed to ensure that a maximum of fungal diversity was covered involved broad substrate sampling (*n*=1080) coupled with a low-throughput isolation and diversity analysis. By contrast, that developed to ensure maximum coverage of bacterial diversity involved a very low level of substrate sampling (*n*=3) coupled with a high-throughput isolation and diversity analysis. The use of these two approaches was methodologically satisfactory and thought-provoking. We identified a number of ASVs by both the metabarcoding and isolation approaches, but also ASVs specific to each approach. Our findings confirm the power of culture-independent strategies, which make it possible to detect much larger numbers of ASVs — six to seven times more for bacteria and fungi — than in the culture-dependent strategy, whether involving high- or low-throughput community-based isolation. However, importantly, they also show that culture-dependent approaches may not be as weak as previously assumed, for at least two reasons.

### The two approaches yield consistent results, particularly for bacteria

The high-throughput isolation strategy employed for bacteria yielded a community profile similar to that obtained with the metabarcoding approach. Bacterial ASV richness was lower with the culture-dependent approach, but most of the dominant ASVs were recovered on a single synthetic medium. Thus, the differences in richness between metabarcoding and culture-dependent approaches reflected differences in sampling depth, with some rare taxa difficult to isolate by culture. However, this difference may result from the strict filter applied, which would have decreased richness. This filter was required to remove PCR and/or sequencing artifacts, which were readily identified in the culture-dependent approach. Indeed, the systematic co-occurrence of ASVs within wells with only one single nucleotide polymorphism reflected sequencing artifacts rather than the isolation of phylogenetically close strains from the same well. It is, therefore, highly probable that certain rare taxa not found by the culture-dependent approach are actually sequencing errors.

As stated above, the use of a single synthetic medium in a single set of growing conditions was sufficient for isolation of the most abundant taxa of the residue microbiota. For example, some abundant taxa affiliated to *Massilia* and *Flavobacteria* were not recovered because the dilution of TSA (1/10) used was not suitable for their isolation (Baldani *et al*., 2014; Nishioka *et al*., 2016). However, one strain from the recently identified superphylum Patescibacteria (Rinke *et al*., 2013), the members of which are not easy to isolate in pure culture (Lewis *et al*., 2020), was successfully obtained. The conditions used therefore made it possible to isolate diverse bacterial strains.

About 40% of the bacterial strains obtained by culture methods were not detected with the metabarcoding approach. Interestingly, most of these strains belonged to Actinobacteria and Firmicutes. The reasons for the underestimation of the presence of these bacterial phyla within the residue microbiome by the culture-independent approach remain unclear. Biases introduced during DNA extraction procedures for endospore-forming Firmicutes may be responsible (Filippidou *et al*., 2015). However, other parameters must also underlie these differences because (i) the family of Actinobacteria underestimated in our study (Microbacteriaceae) does not form spores (Evtushenko and Takeuchi, 2017) and (ii) some spore-forming Firmicutes were detected by metabarcoding but not in culture-dependent approaches.

#### The two approaches provide complementary information, particularly for fungi

A strategy combining broad substrate sampling with low-throughput isolation revealed differences between the fungal profiles obtained by isolation and metabarcoding. With the exception of *Alternaria*, some of the most abundant taxa in the metabarcoding approach were absent from the fungi identified by culture-based methods. This was the case for *Lesptosphaeria* (*L. maculans*), *Plenodomus* (*P. biglobosus*) and *Oculimacula* (*O. yallundae*), all of which are known to be pathogenic. However, a very large number of strains from different genera (e.g. *Fusarium, Epicoccum*, *Boeremia*, *Botrytis, Mortierella*, and *Mucor*) were frequently isolated, despite their low proportions in the metabarcoding analysis. Metabarcoding is, thus, not sufficient in itself, and isolation definitely provides added value for the accurate characterization of the fungal community in residues. Moreover, three ASVs from three strains from the culture collection matched three unidentified ASVs with 100% identity. The strains were not identified due to a lack of identity with sequences in databases, but it would be possible to characterize them phenotypically, by enzyme profiling or pathotyping, to obtain additional information about the residue-associated fungal community. The isolation medium may affect the difference in growth rates between fungal species (selective effect), potentially accounted for differences between the two approaches. Whatever the medium used, the intrinsic growth rate *in vitro* may vary from a few millimeters (e.g. *Pyrenopeziza brassica* and *L. maculans*) to a few centimeters per week (*Fusarium* sp., which grow very rapidly in rich environments and are generally overrepresented), and rapidly growing strains may have been favored.

### Some methodological considerations

In general, amplicon-based metagenomic approaches are based on small fragments, whereas the molecular identification of single strains can be based on longer barcodes and Sanger sequencing (Hamad *et al*., 2017). This is one of the limitations of our comparison between culture-dependent and culture-independent approaches. Indeed, we found that an ASV obtained by metabarcoding could display 100% sequence identity to several cultured strains. This reflects the PCR fragment used for the culture-dependent approach: in metabarcoding, only the ITS1 part was used for identification, whereas, in single-strain barcoding, the entire ITS1-5.8S-ITS2 region was sequenced. Intraspecies polymorphism may be hidden because it leads to different ITS1 sequences (e.g. 100% identity of an ASV detected by metabarcoding to eight ASVs from the genus *Epicoccum*), so the culture-independent approach is likely to underestimate taxonomic – and therefore functional – fungal diversity. Some studies (e.g. Krehenwinkel *et al*., 2019; Heeger *et al*., 2018) have suggested using long-read sequencing techniques for metabarcoding to improve taxonomic resolution, thereby providing a more accurate perception of the functional diversity of the residue microbial community. Indeed, although the ITS region generally performs well as a fungal barcoding marker, it is less efficient for some genera and it may be necessary to sequence several genes to identify a strain correctly (Raja *et al*., 2017). Even in this case, taxonomic identification is dependent on the presence of the corresponding sequences in international databases. For bacteria, the v4 region of 16S rRNA gene has poor discriminatory power at species level (Callahan *et al*., 2019), but here we analyzed this same fragment by the same technique. Together with the high-throughput isolation, this may partly explain the greater similarity of the results obtained by the two approaches for bacterial profiles than for fungal profiles It appears to be essential to combine the two approaches if we wish to go beyond descriptive aspects and characterize the ecological function of a microbial community. The main advantage of microbial isolation is that the different strains can be preserved and then tested — individually or after the creation of synthetic communities (Großkopf and Soyer, 2014; Sergaki *et al*. 2018) — to decipher these ecological functions and identify positive specific interactions (Durán *et al*., 2018; Liu *et al*., 2019; Zhuang *et al*., 2020). Conversely, metabarcoding limits the biases linked to the cultivable/non-cultivable nature of microbes in the fine characterization of communities, and dedicated statistical analyses, such as linear discriminant analysis or ecological network analysis, are complementary approaches that could be used to identify putative microbial interactions (e.g. Kerdraon *et al*., 2019c; 2000).

### A still imprecise picture of residue microbial diversity that might be improved by culturomics: when one bias replaces another

The cultivable fraction of the fungal and bacterial communities of arable crop residues appeared to be neither as low as the ‘1%’ paradigm long associated with the rhizosphere microflora (Martiny, 2019), nor as high as the 50% sometimes reported for the phyllosphere microflora (Müller and Ruppel, 2014; Bai *et al*., 2015). In our study, this cultivable fraction was relatively high, at a value intermediate between those for soil and living plants. This finding is consistent with residues acting as an ecotone, at the interface between the soil and the phyllosphere (Kerdraon *et al*., 2019a). We used only one culture medium here, but different media and growth conditions could be used to highlight the power of culture-dependent approaches and their complementarity with metabarcoding (Lagier *et al*., 2012; Pfleiderer *et al*., 2013). The use of a single medium undoubtedly limited the scope of our results, particularly relative to those obtained by culturomics — high-throughput approaches based on multiple growth conditions (several media, selective or not, aerobic, anaerobic or micro-aerophilic, at different temperatures and over different time steps) — for other substrates. Culturomics can detect microbial species previously unknown or unisolated. It has emerged as a successful tool for isolating large numbers of bacteria, but has not yet been comprehensively applied to the description of fungal communities on substrates from agroecosystems. The application of culturomics to plant and soil microbiomes lags behind its use in human microbiome studies (Sarhan *et al*., 2019), which has led to a revival of culture-dependent approaches (Lagier *et al*., 2012; 2015, 2016) increasingly performed in parallel with culture-independent approaches (Ellis *et al*., 2003; Mijangos *et al*., 2009) combining cloning, sequencing (Kaplan *et al*., 2013), and metabarcoding (Lagier *et al*., 2012).

Several of the studies mentioned above highlight the difficulty describing bacterial communities exhaustively even if several sets of culture conditions are used. For fungi, one strong limiting factor is the differences in growth rate between taxa, which cannot necessarily be diminished through the use of different growth conditions. *L. maculans* and *Z. tritici* are two good examples: these two pathogenic species are widely found on oilseed rape and wheat residues, respectively, and are widely cultivable, but were never isolated here, probably due to competition with fast-growing species, such as *Fusarium* sp., many of which originate from the soil and colonize the surface of the residues. The use of selective media may be a partial solution, but this would introduce biases modifying the representation of microbiological diversity. These two approaches are complementary and could be combined, but it should be borne in mind that technical biases may remain, preventing the exhaustive characterization of microbial communities. For example, for fungal communities, some of the primers used for metabarcoding poorly amplify species from genus *Puccinia,* but *Puccinia* sp., which is a strict biotroph, cannot be isolated by culture.

## Conclusion

This is the first comparative analysis of this type to have been performed on arable crop residues, a matrix of major agroecological importance. By focusing on methodological aspects, we have extended a broader experimental work (Kerdraon *et al*., 2019b; 2019c; 2020), paving the way for further technical improvements. The isolation approach used here for bacteria was based on a single medium. However, it was sufficient to isolate not only most of the most abundant taxa identified by metabarcoding, but also some ASVs not retrieved by the culture-independent approach. For fungi, metabarcoding probably provided a more accurate picture than culture techniques, because of the aforementioned biases inherent to all culture-dependent approaches. However, several bacterial and fungal ASVs not retrieved by metabarcoding were, nevertheless, retrieved by culture. The power of the culture-independent strategy was confirmed, but the culture-dependent approach was found to be less weak than anticipated, highlighting the need to continue using ‘classic microbiology’ techniques. The two approaches were found to be complementary, and their combination provided a more complete and accurate description of the microbiological diversity of residues, without the need for culturomics. This is an important point, and may even be considered a methodological advantage, because any deviation from the initial environmental parameters in axenic culture is known to modify microbial community structure.

## Acknowledgments

We thank Julie Sappa for her help correcting our English.

## Availability of data and materials

The raw metabarcode sequencing data is available from the European Nucleotide Archive (ENA) under the study accession PRJEB27255 (Sample SAMEA4723701 to SAMEA4724326), already presented in Kerdraon *et al*. (2019). The molecular typing data of the strains isolated is presented in Table S1 (bacteria) and Table S2 (fungi).

## Supplementary material

**Table S1. Bacterial strains isolated from three sets of residues.** Strains were isolated after the maceration of two pieces of wheat residue (W) and one piece of oilseed rape (O) residue collected from the wheat monoculture plot (W_M_) and the two wheat-oilseed rape rotation plots (W_R_ and O_R_) in October 2015. Molecular typing of the strains was performed with the v4 region of the 16S rRNA gene. In total, 158 amplicon sequence variants (ASVs) were obtained with this molecular marker. The number of strains for each ASV is displayed. The number of SNPs between ASVs recovered by the culture-dependent and metabarcoding approaches is indicated.

**Table S2. Fungal strains isolated from pieces of residue.** Strains were isolated on PDA and identified from 1080 pieces of wheat (W) and oilseed rape (O) residue collected from the wheat monoculture plot (W_M_) and the two wheat-oilseed rape rotation plots (W_R_ and O_R_) in 2015-2016 and 2016-2017. Molecular typing of the strains was performed with the ITS1-5.8S-ITS2 region. In total, 424 strains were obtained with this molecular marker. The corresponding morphotype photographs are provided in **Fig. S2**, when available.

**Table S3. Genera of fungal strains isolated and identified (*n*=424) from 1080 pieces of wheat (W) and oilseed rape (O) residue collected from the wheat monoculture plot (W_M_) and the two wheat-oilseed rape rotation plots (W_R_ and O_R_) in 2015-2016 and 2016-2017.** Molecular typing of the strains was performed with the ITS1-5.8S-ITS2 region. Data for the three sampling periods (October, December and February) were pooled for each cropping season. The genera not detected by metabarcoding in the study by Kerdraon *et al*. (2019b) are indicated by *.

**Figure S1. Photographs of 171 fungal morphotypes representative of the 424 fungal taxa.** Strains were isolated on PDA medium (from 4 to 10 days of culture at 18°C in the dark) from 1080 pieces of wheat and oilseed rape residue (W_M_, W_R_ and O_R_) in 2015-2016 and 2016-2017. The genus, and species when known, to which each morphotype was affiliated are indicated in **Table S2**.

