## Supplementary material for "Synergy of culture-dependent molecular identification and whole-community metabarcode sequencing for characterizing the microbiota of arable crop residues": Table S3

**Table S3. Genera of fungal strains isolated and identified (*n*=424) from 1080 pieces of wheat (W) and oilseed rape (O) residue collected from the wheat monoculture plot ($W_M$) and the two wheat-oilseed rape rotation plots ($W_R$ and $O_R$) in 2015-2016 and 2016-2017.** Molecular typing of the strains was performed with the ITS1-5.8S-ITS2 region. Data for the three sampling periods (October, December and February) were pooled for each cropping season. The genera not detected by metabarcoding in the study by Kerdraon *et al.* (2019b) are indicated by *.

| | 2015-2016 | | | 2016-2017 | | |
|---|---|---|---|---|---|---|
| | $W_M$ | $W_R$ | $O_R$ | $W_M$ | $W_R$ | $O_R$ |
| *Actinomucor* * | 0 | 0 | 0 | 0 | 0 | 1 |
| *Alternaria* | 1 | 13 | 1 | 0 | 0 | 8 |
| *Boeremia* * | 0 | 0 | 0 | 0 | 0 | 3 |
| *Botrytis* * | 0 | 0 | 0 | 0 | 0 | 1 |
| *Chaetomium* | 3 | 0 | 0 | 0 | 0 | 4 |
| *Cladorrhinum* | 0 | 2 | 5 | 1 | 0 | 0 |
| *Cladosporium* | 0 | 0 | 1 | 0 | 1 | 7 |
| *Epicoccum* | 3 | 6 | 7 | 9 | 10 | 13 |
| *Fusarium* | 18 | 15 | 13 | 58 | 68 | 38 |
| *Laetisaria* | 0 | 0 | 0 | 0 | 0 | 0 |
| *Lophodermium* | 0 | 0 | 0 | 0 | 0 | 0 |
| *Leptosphaeria* | 2 | 0 | 0 | 1 | 0 | 0 |
| *Microdochium* | 0 | 2 | 0 | 1 | 3 | 0 |
| *Mortierella* * | 6 | 5 | 20 | 5 | 4 | 6 |
| *Mucor* | 8 | 2 | 6 | 5 | 1 | 12 |
| *Neurospora* * | 0 | 0 | 0 | 0 | 1 | 0 |
| *Nigrospora* | 0 | 9 | 0 | 0 | 0 | 0 |
| *Phaeosphaeria* | 1 | 0 | 0 | 1 | 0 | 0 |
| *Septoriella* | 0 | 0 | 0 | 0 | 0 | 0 |
| *Stagonosporopsis* * | 0 | 0 | 0 | 0 | 0 | 1 |
| *Trichoderma* | 0 | 5 | 0 | 1 | 0 | 2 |
| *Waitea* | 0 | 1 | 0 | 0 | 0 | 0 |
| NI | 1 | 2 | 0 | 0 | 0 | 0 |
| Total number of strains | 43 | 62 | 53 | 82 | 88 | 96 |
| Total number of genera identified | 8 | 10 | 7 | 9 | 7 | 12 |
