## Supplementary material for "Synergy of culture-dependent molecular identification and whole-community metabarcode sequencing for characterizing the microbiota of arable crop residues": Figure S1

**Figure S1. Photographs of 171 fungal morphotypes representative of the 424 fungal taxa.** Strains were isolated on PDA medium (from 4 to 10 days of culture at 18°C in the dark) from 1080 pieces of wheat and oilseed rape residue ($W_M$, $W_R$ and $O_R$) in 2015-2016 and 2016-2017. The genus, and species when known, to which each morphotype was affiliated are indicated in **Table S2**.

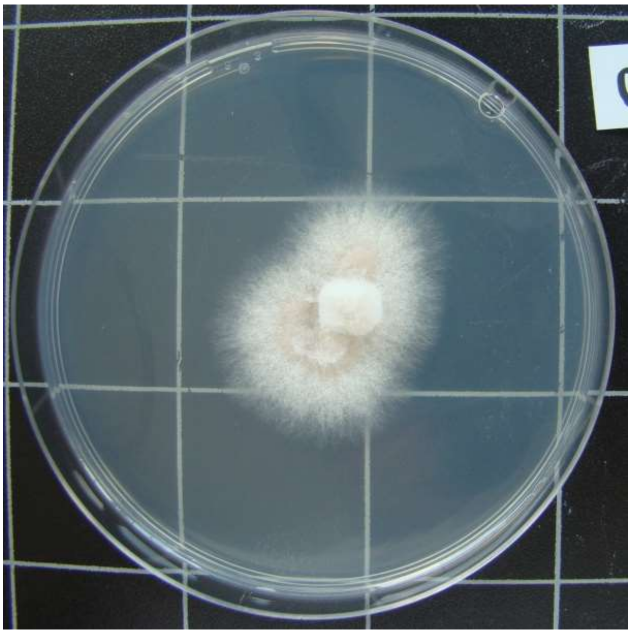

0216-HDG2.A-6

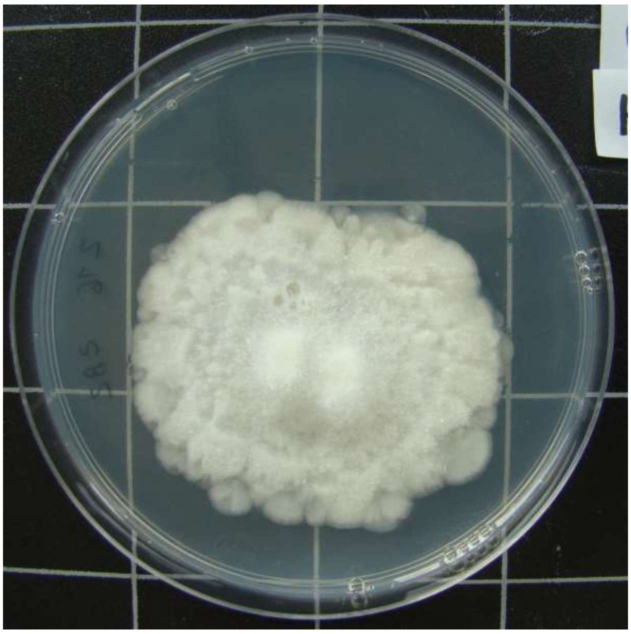

0216-HDG2.B-2

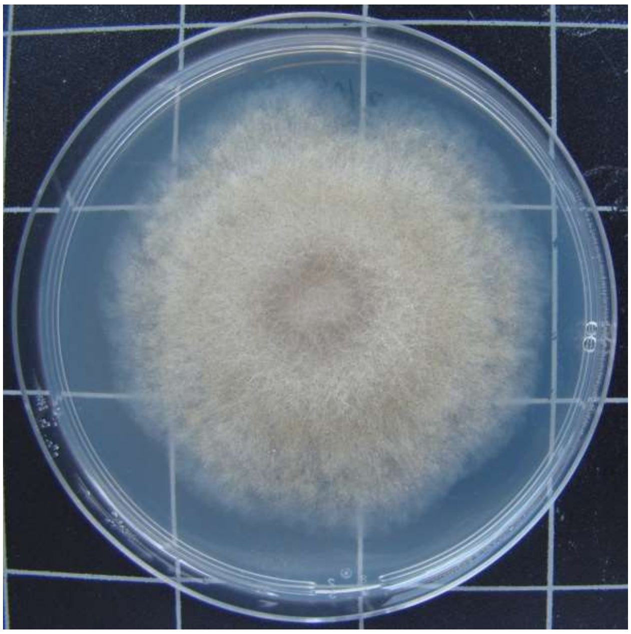

0216-HDG2.C-3

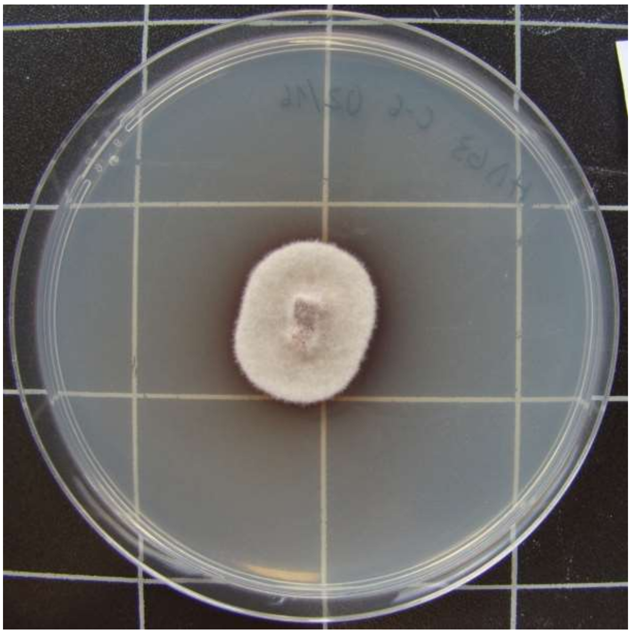

0216-HDG3.C-6

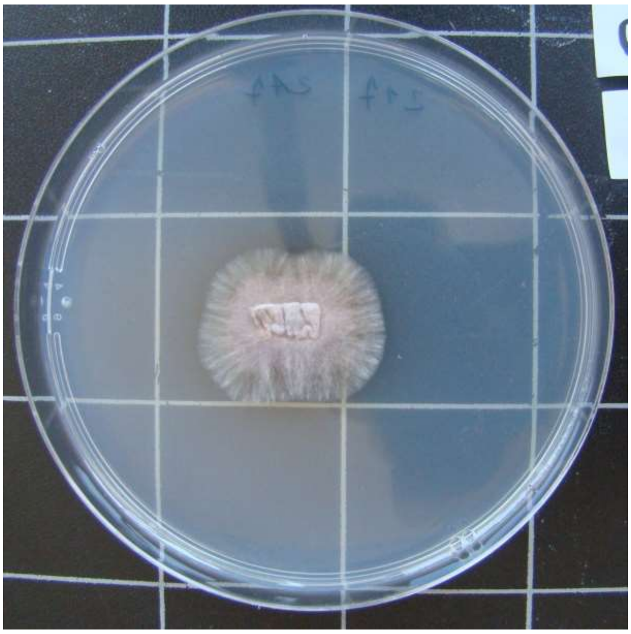

0217-HDG2.A-7

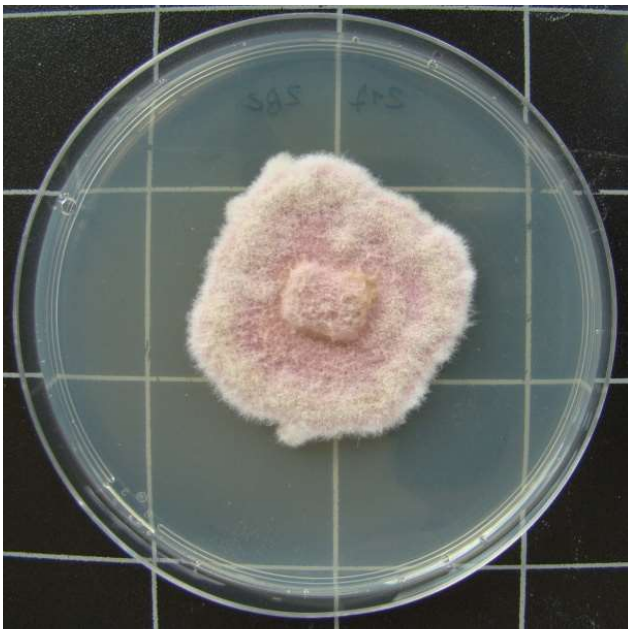

0217-HDG2.B-2

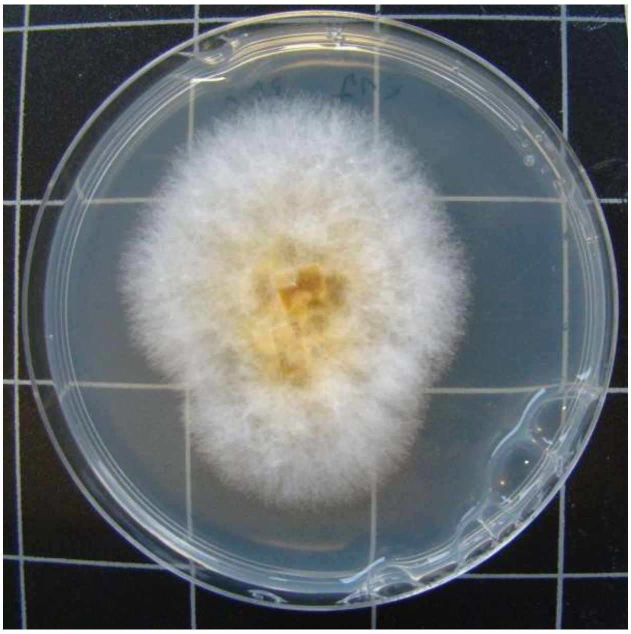

0217-HDG3.E-6

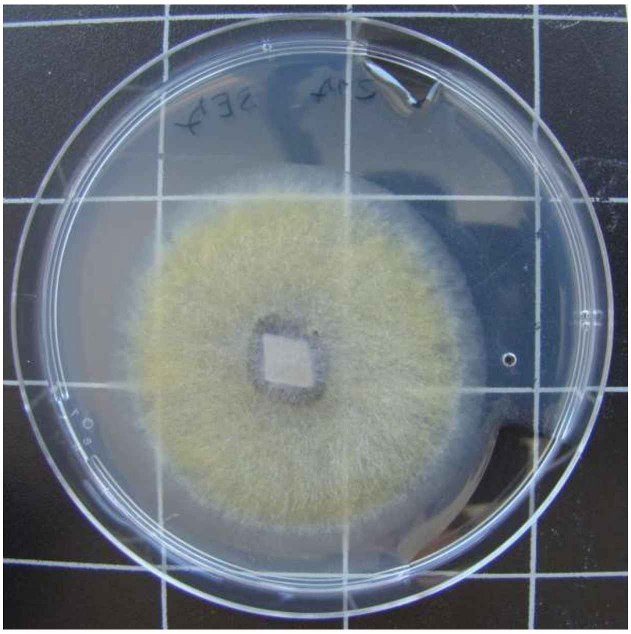

0217-HDG3.E-7

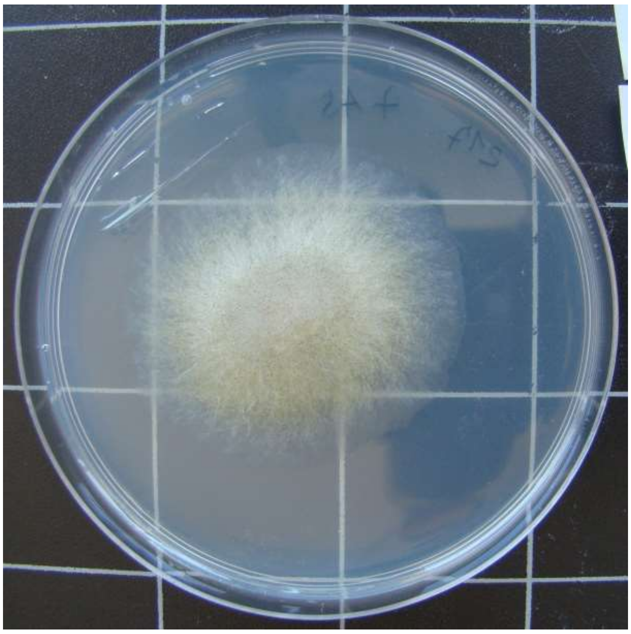

0217-TTF7.A-5

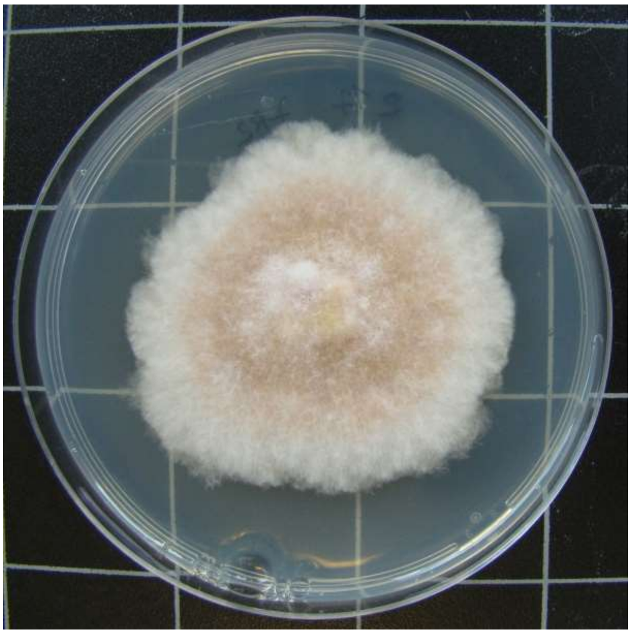

0217-TTF7.B-2

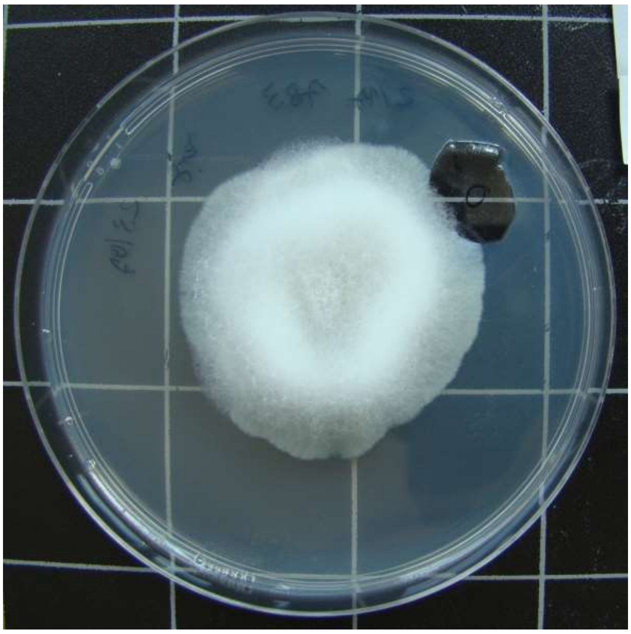

0217-TTF7.B-3

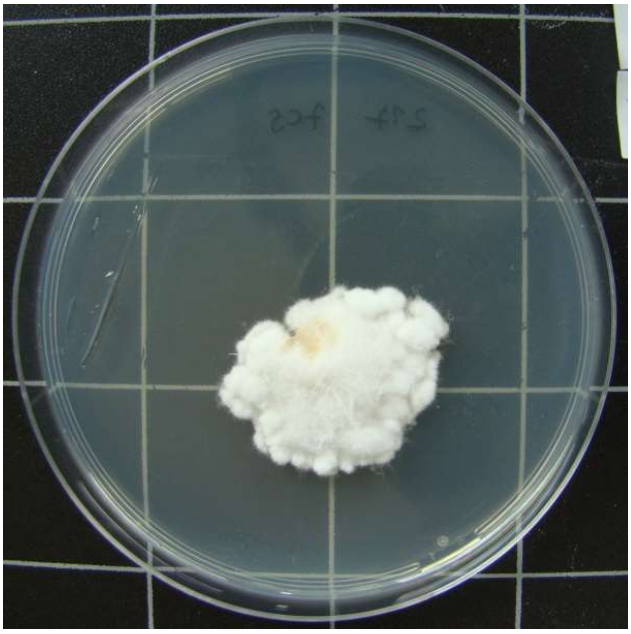

0217-TTF7.C-5

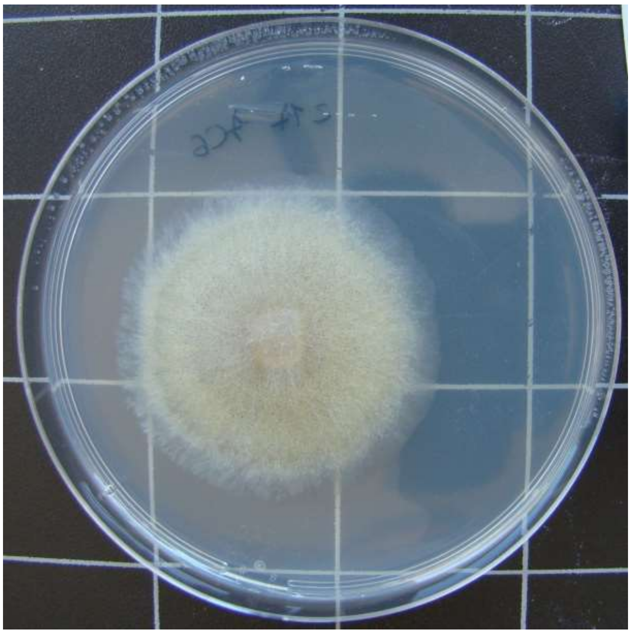

0217-TTF7.C-6

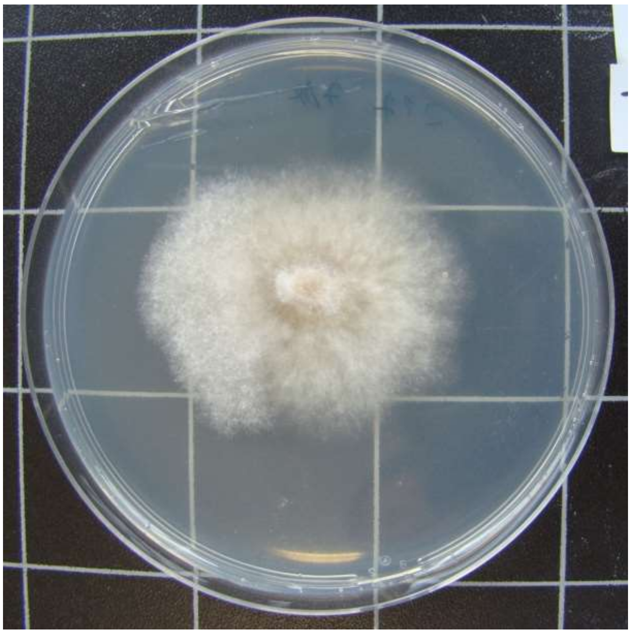

0217-TTF7.D-4

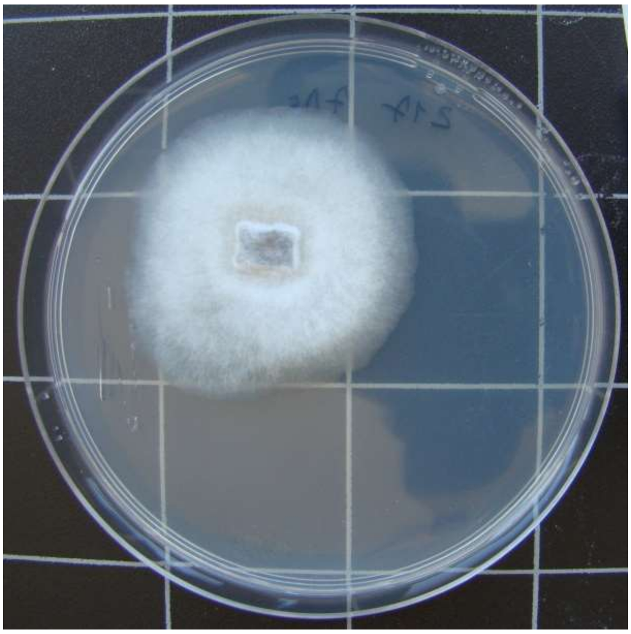

0217-TTF7.D-5

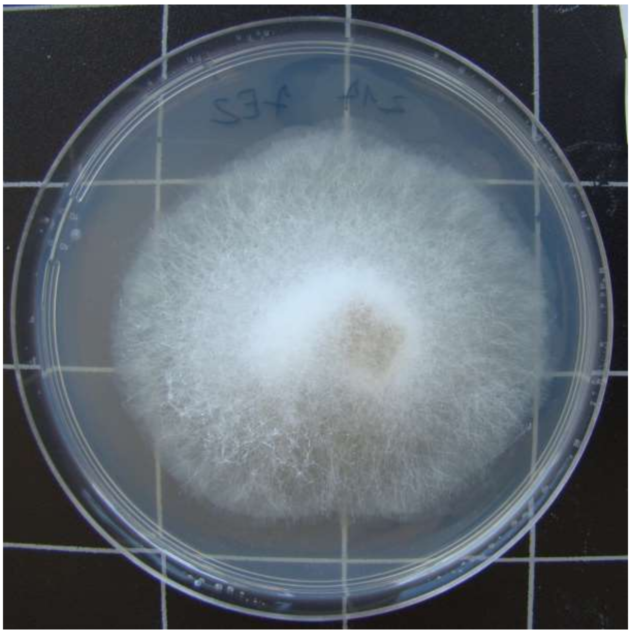

0217-TTF7.E-2

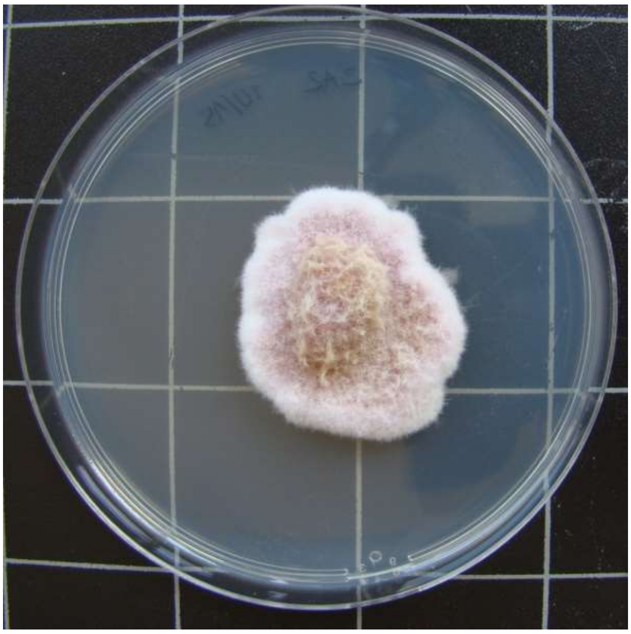

1015-HDG2.A-2

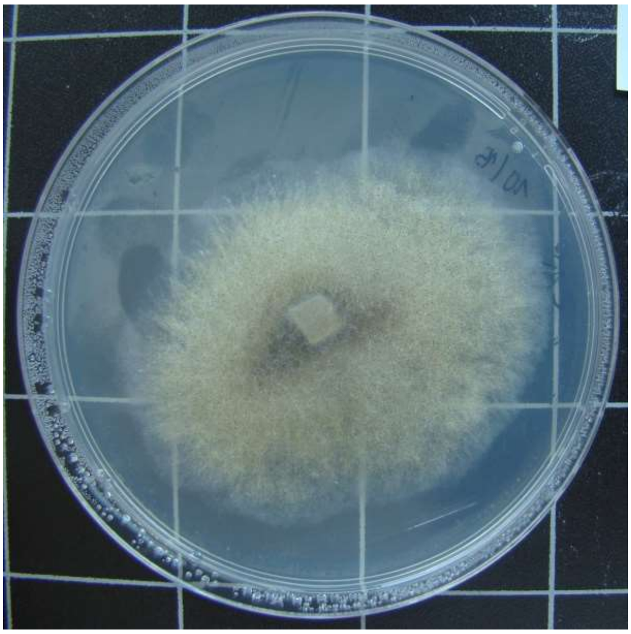

1015-HDG2.A-3

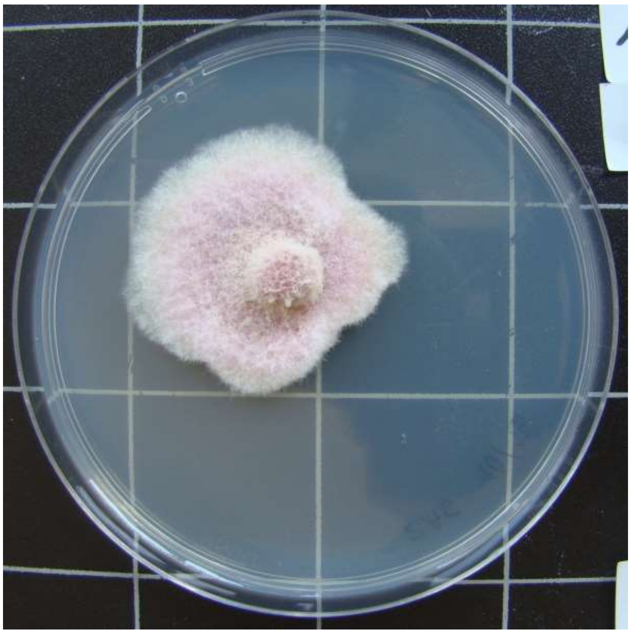

1015-HDG2.A-6

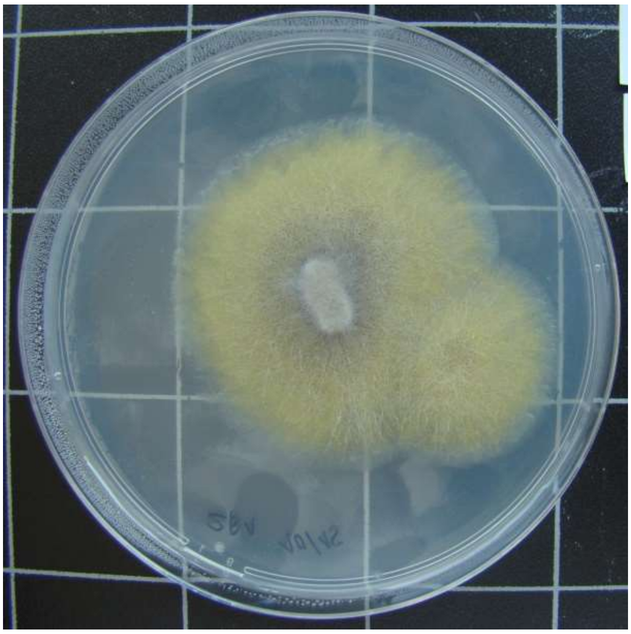

1015-HDG2.B-1

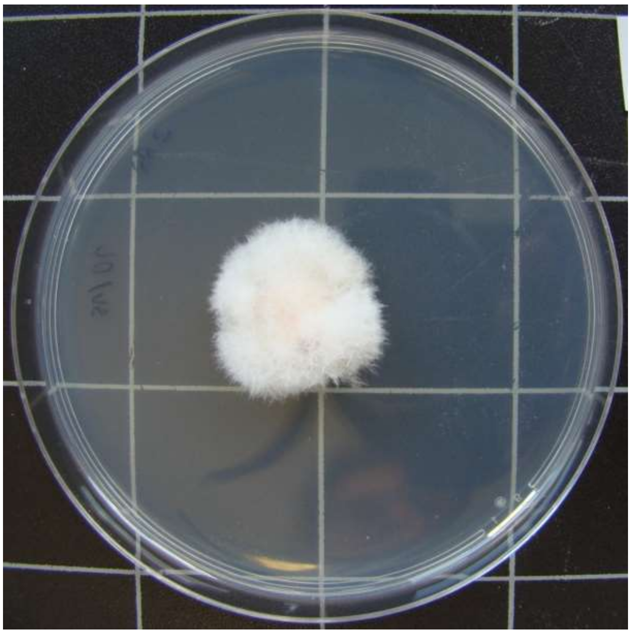

1015-HDG2.B-3

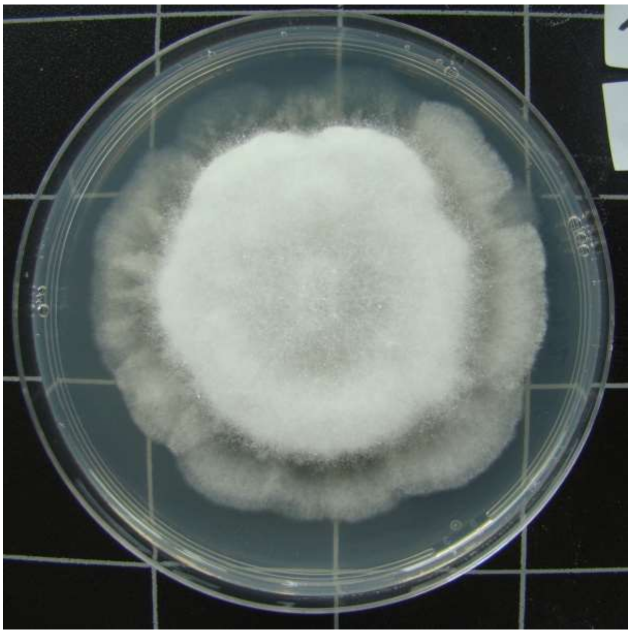

1015-HDG2.B-4

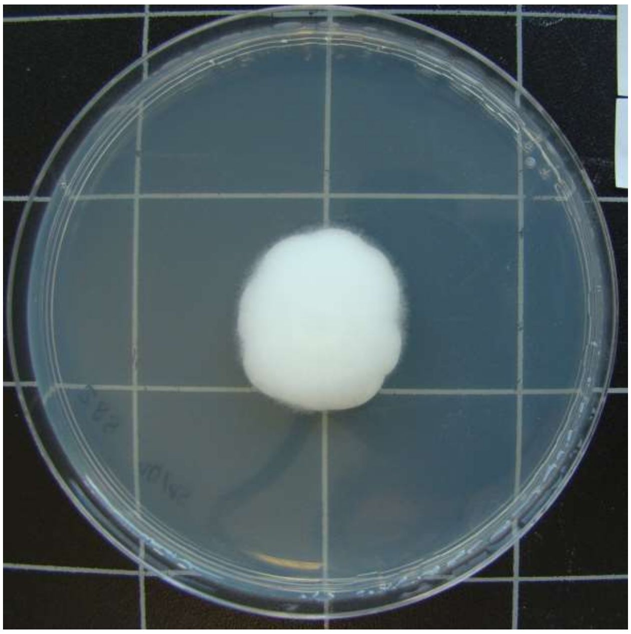

1015-HDG2.B-5

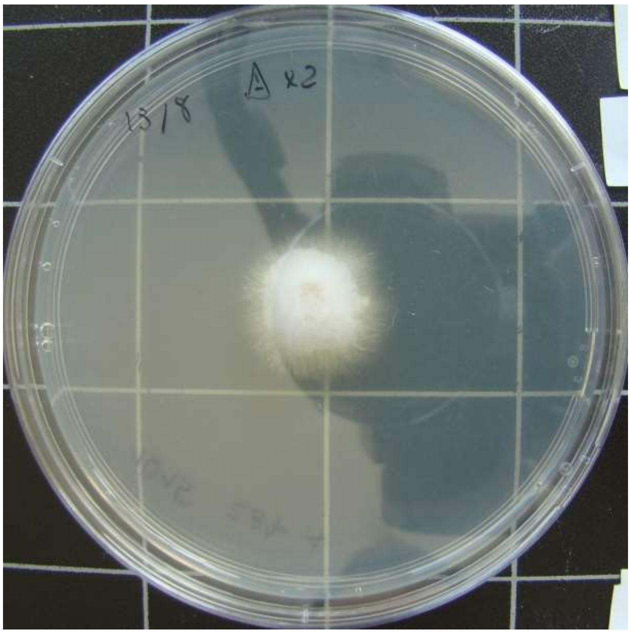

1015-HDG2.B-7

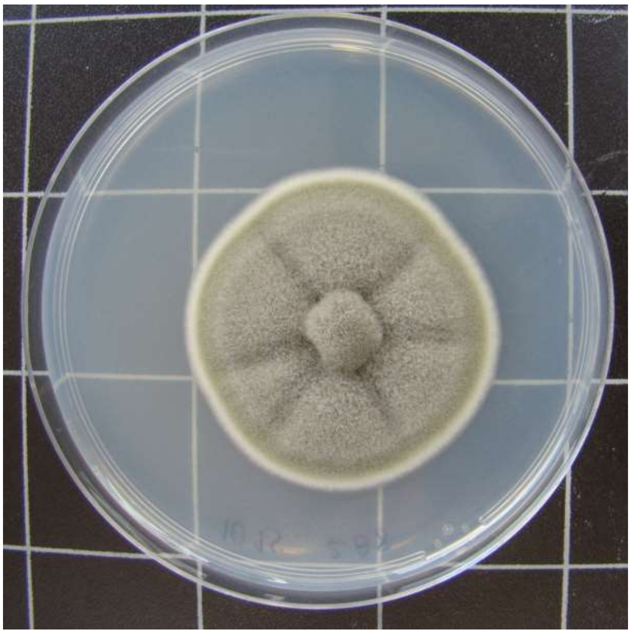

1015-HDG2.B-8

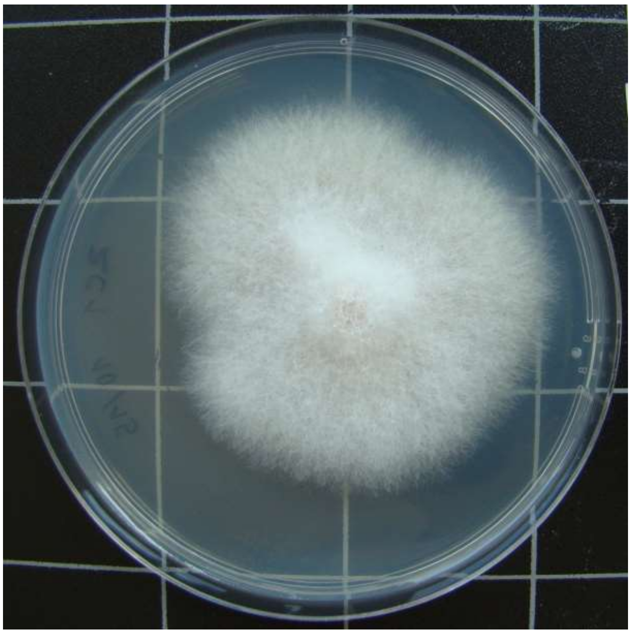

1015-HDG2.C-1

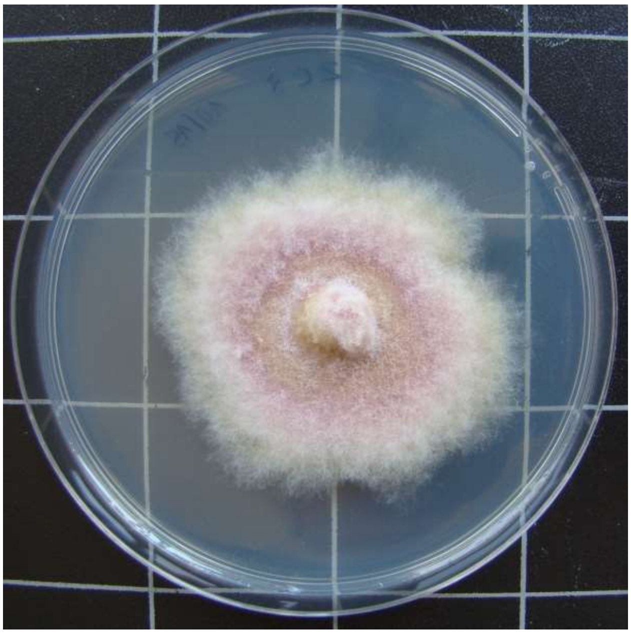

1015-HDG2.C-3

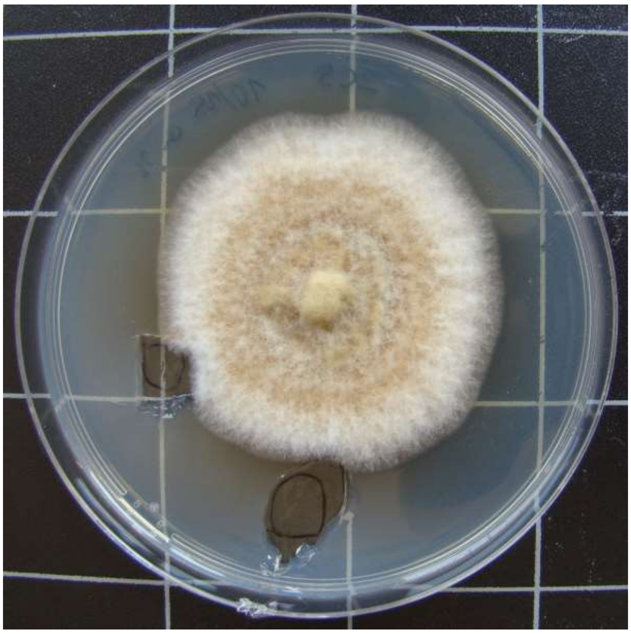

1015-HDG2.C-5

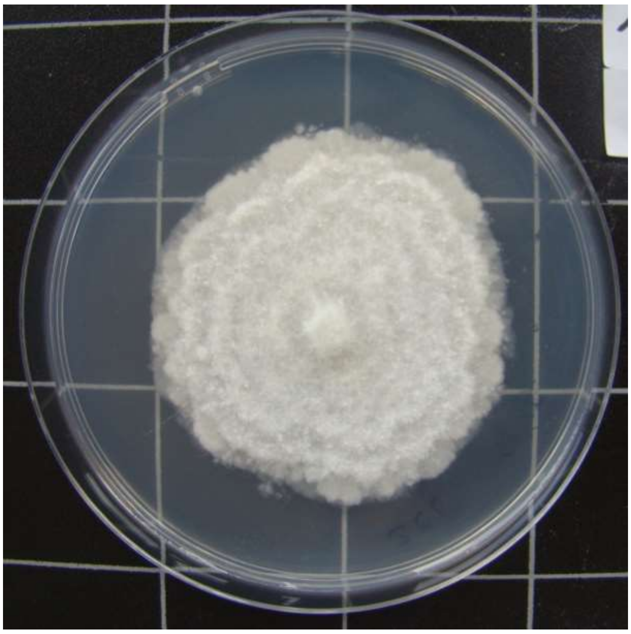

1015-HDG2.C-6

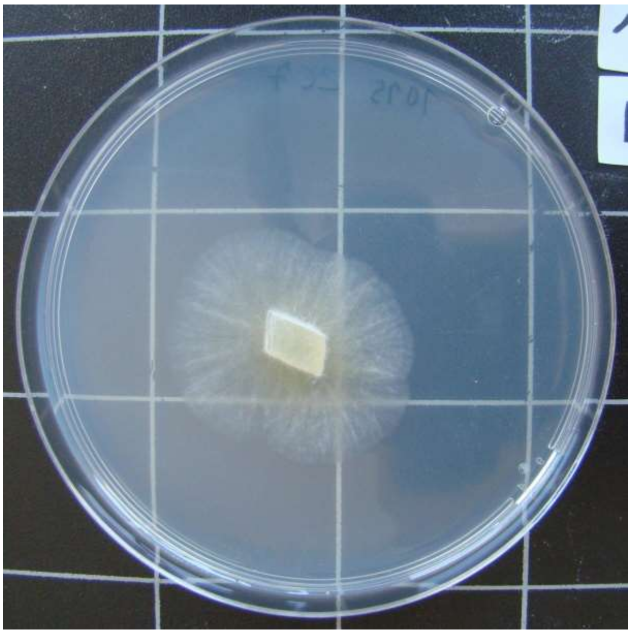

1015-HDG2.C-7

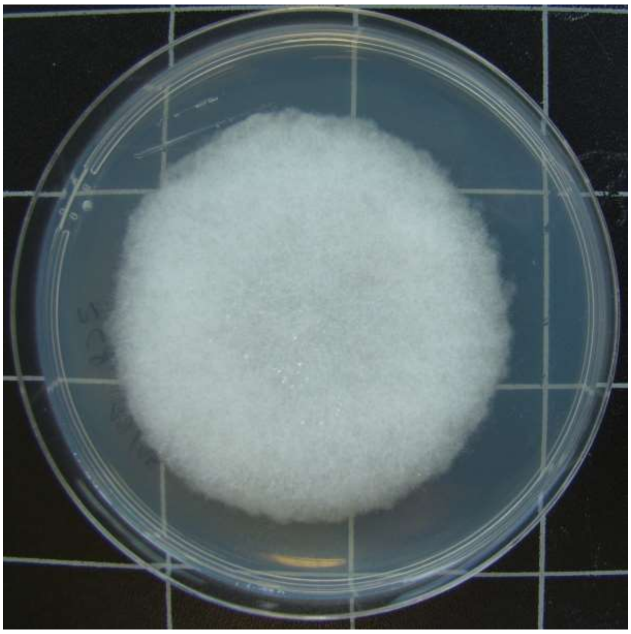

1015-HDG2.C-8

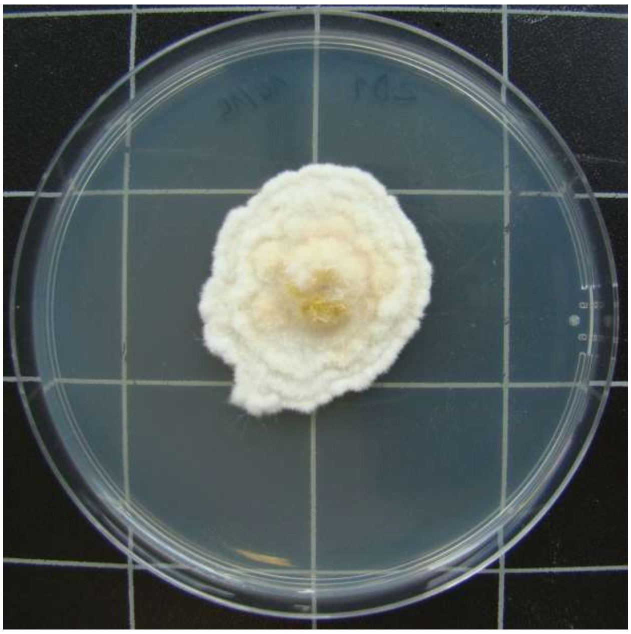

1015-HDG2.D-1

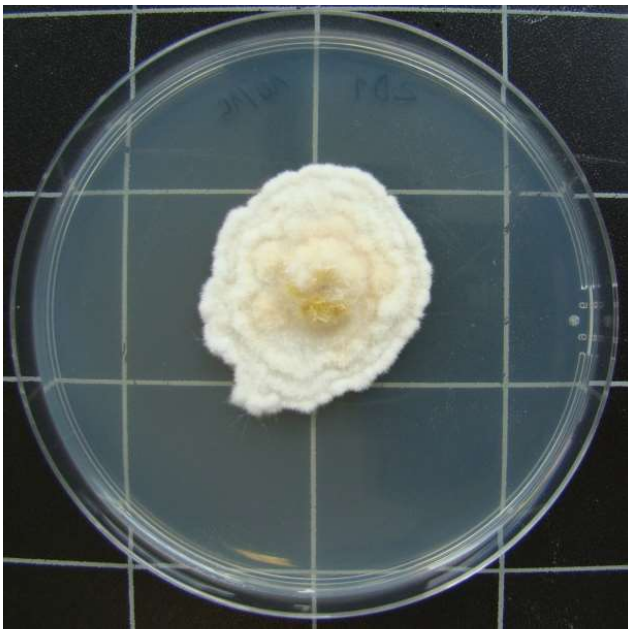

1015-HDG2.D-1

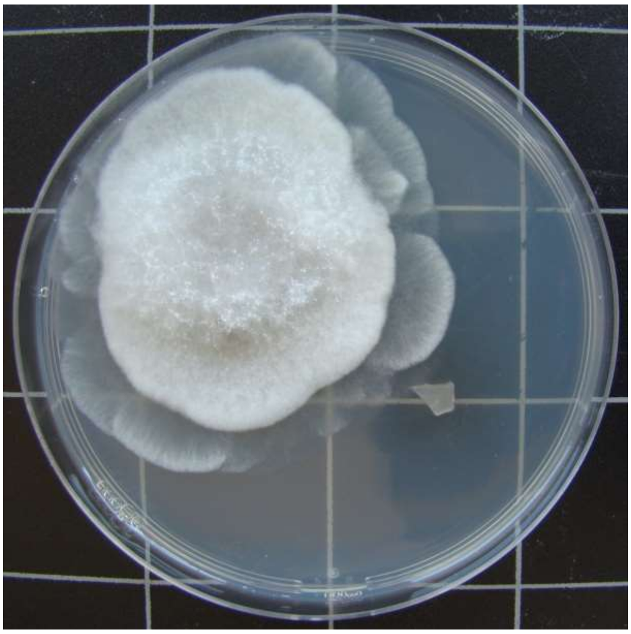

1015-HDG2.D-2

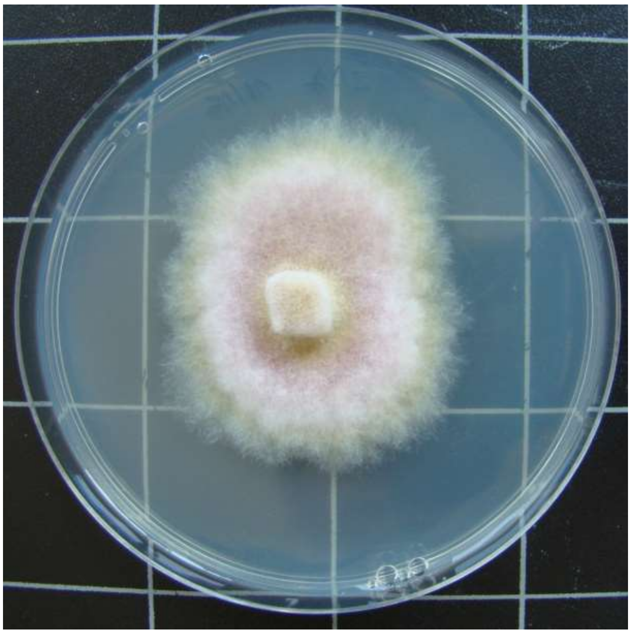

1015-HDG2.D-7

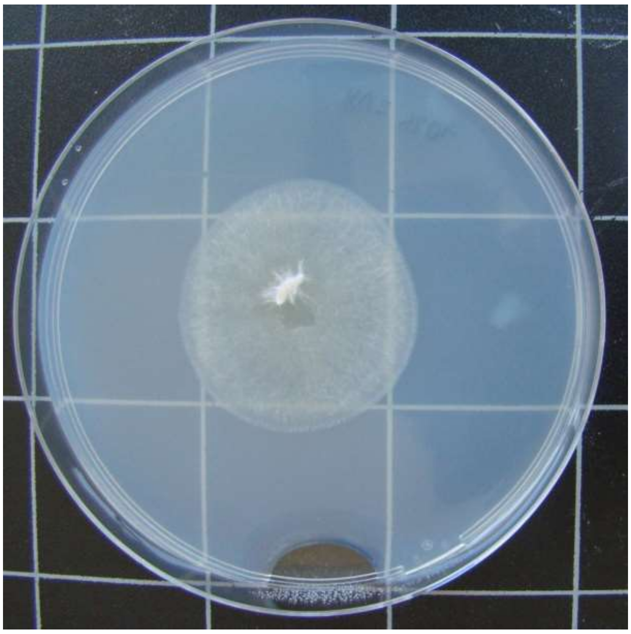

1015-HDG2.D-8

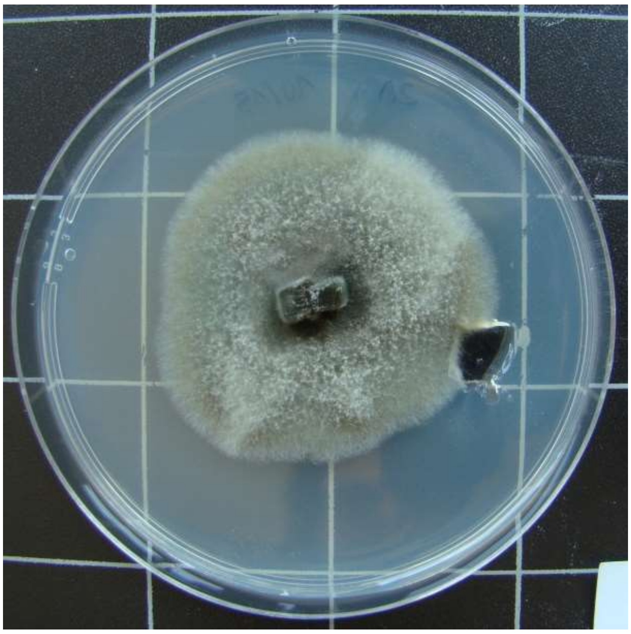

1015-HDG2.D-9

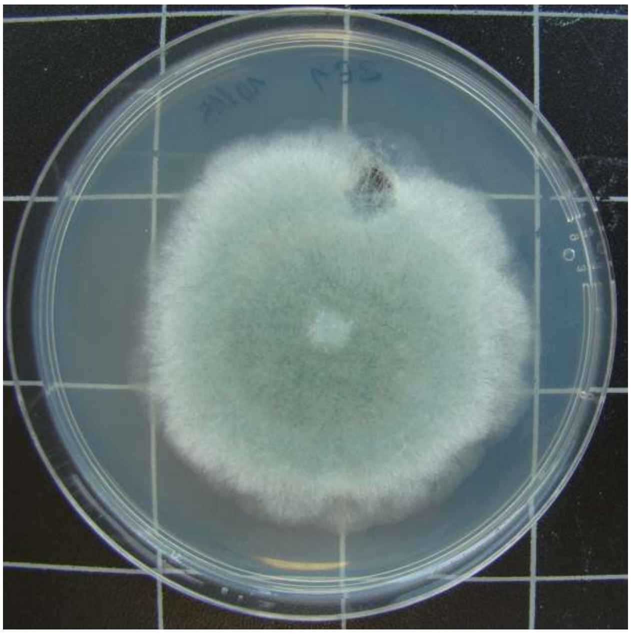

1015-HDG2.E-1

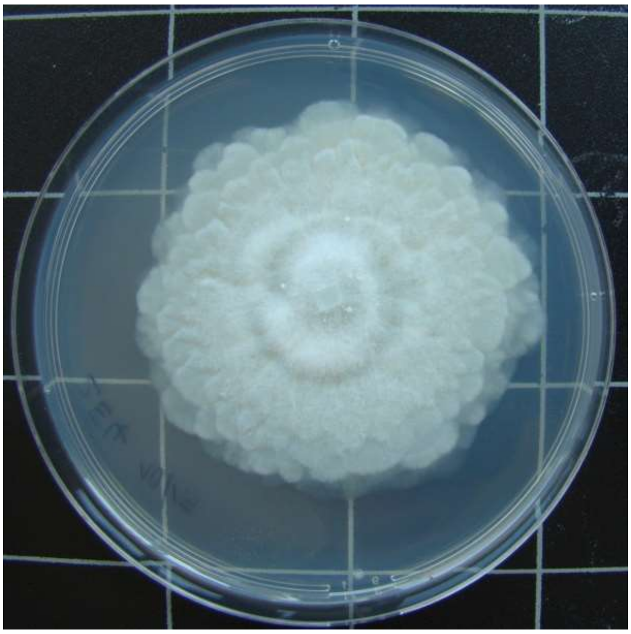

1015-HDG2.E-4

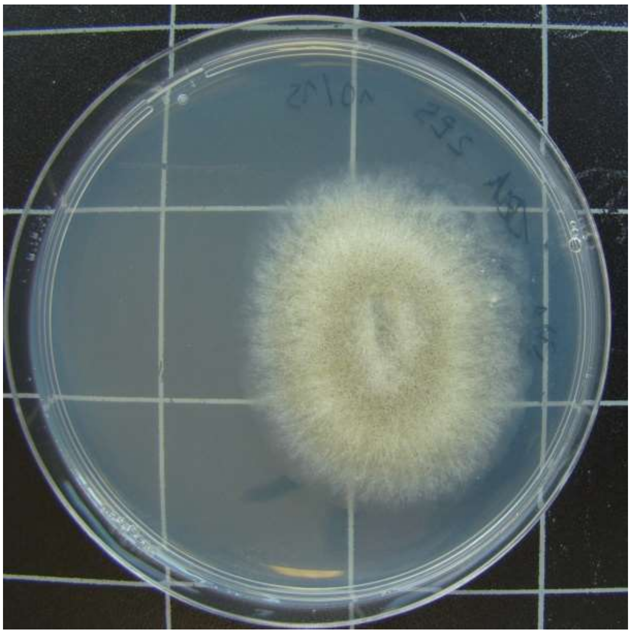

1015-HDG2.E-5

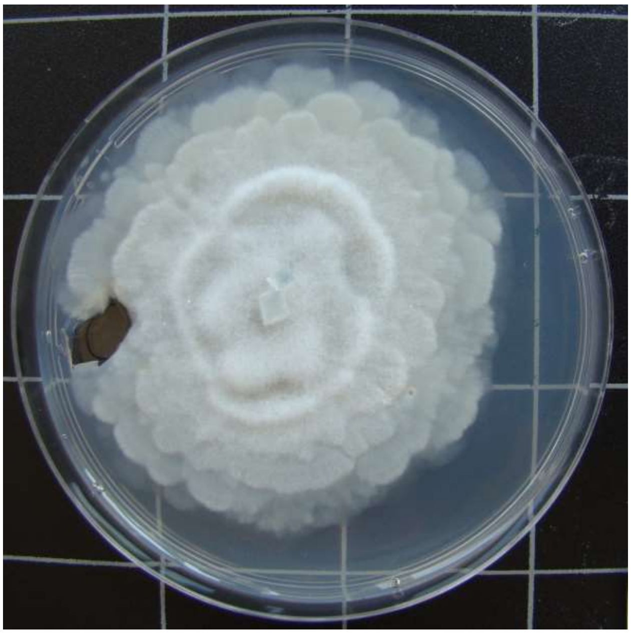

1015-HDG2.E-6

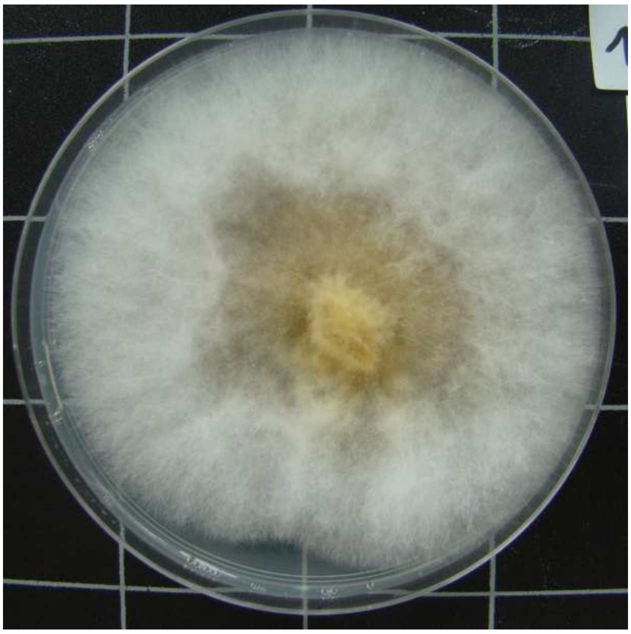

1015-HDG3.A-1

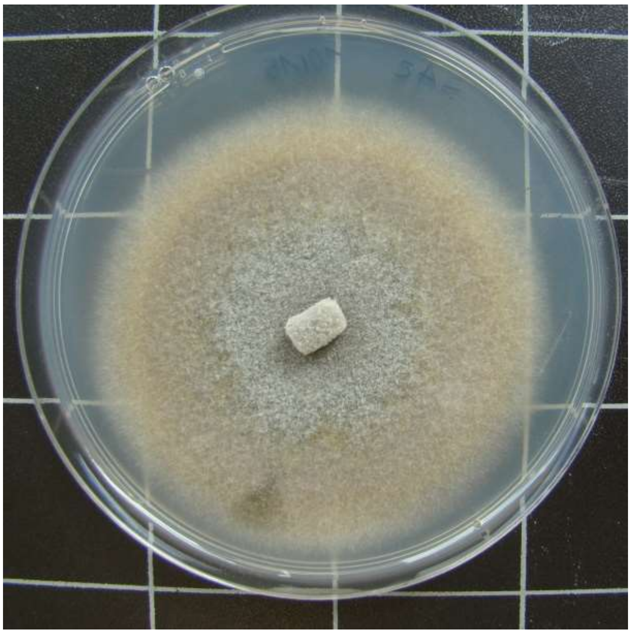

1015-HDG3.A-2

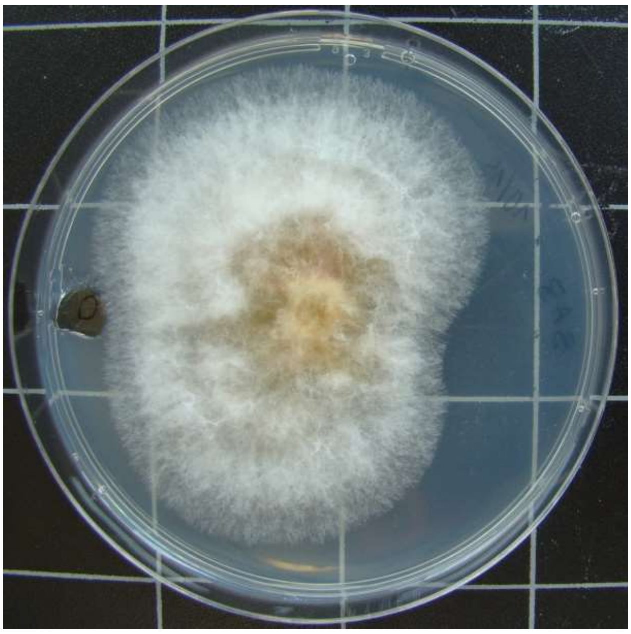

1015-HDG3.A-3

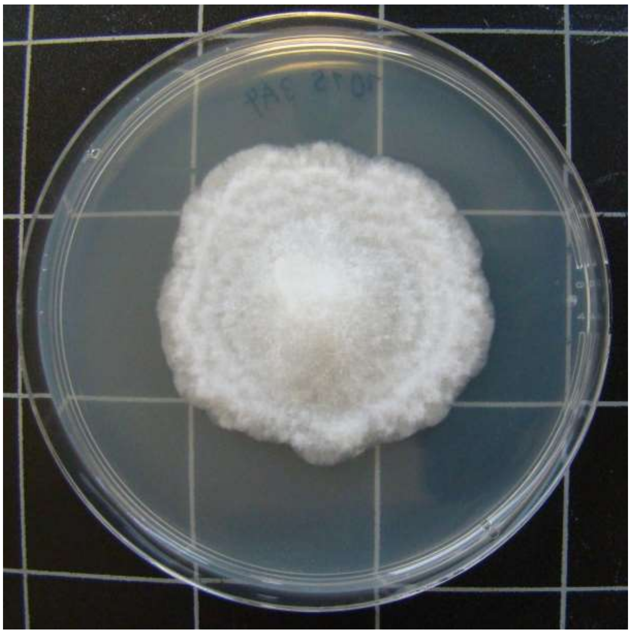

1015-HDG3.A-4

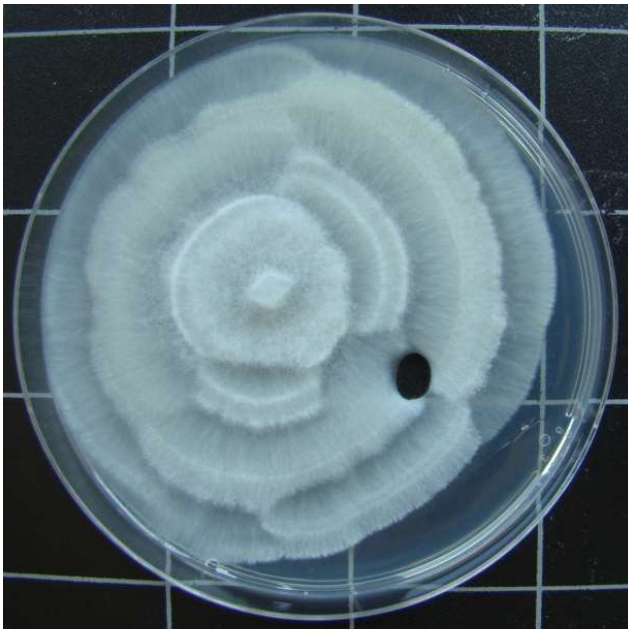

1015-HDG3.A-5

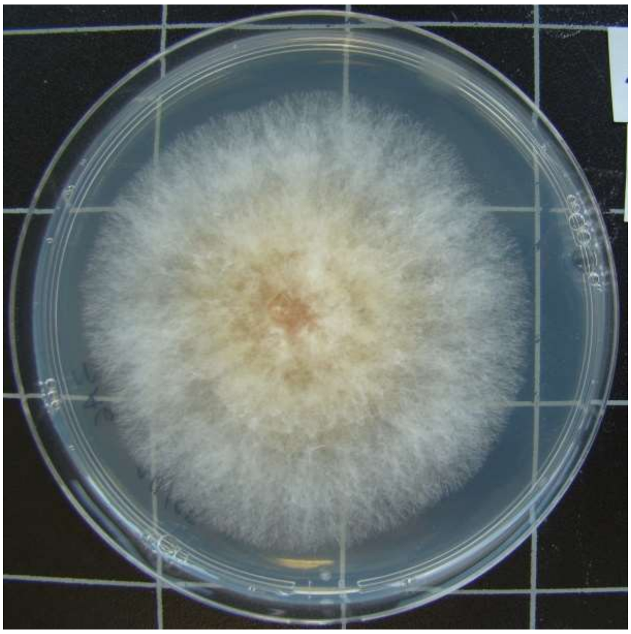

1015-HDG3.A-6

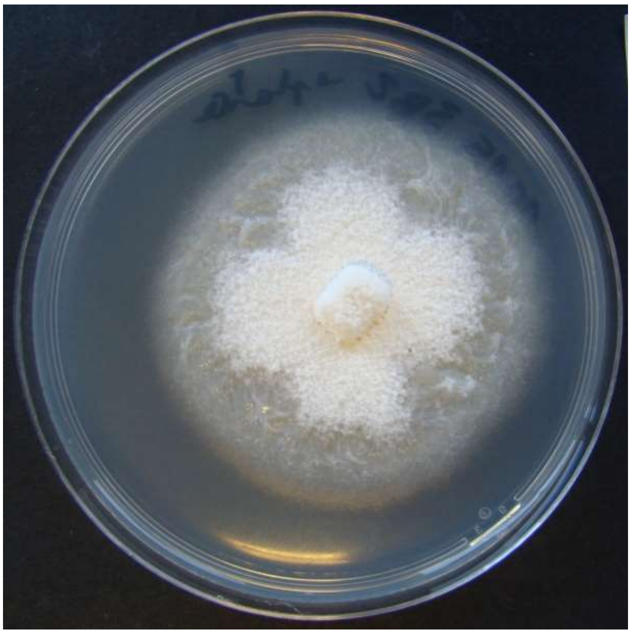

1015-HDG3.B-2

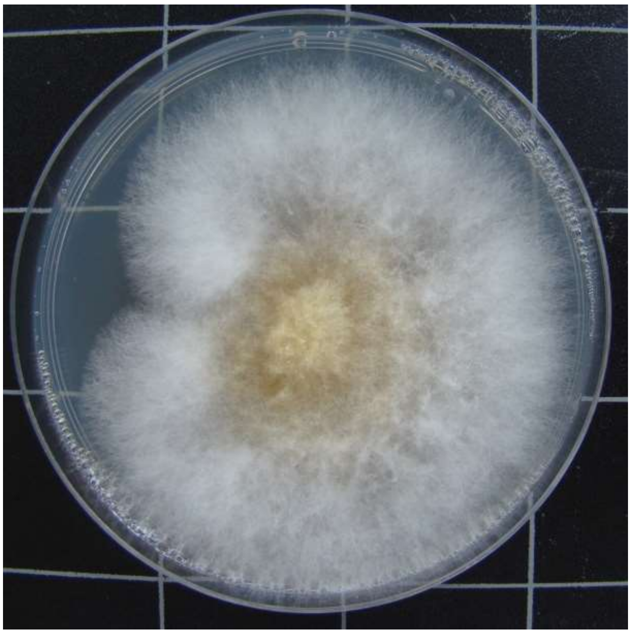

1015-HDG3.B-3

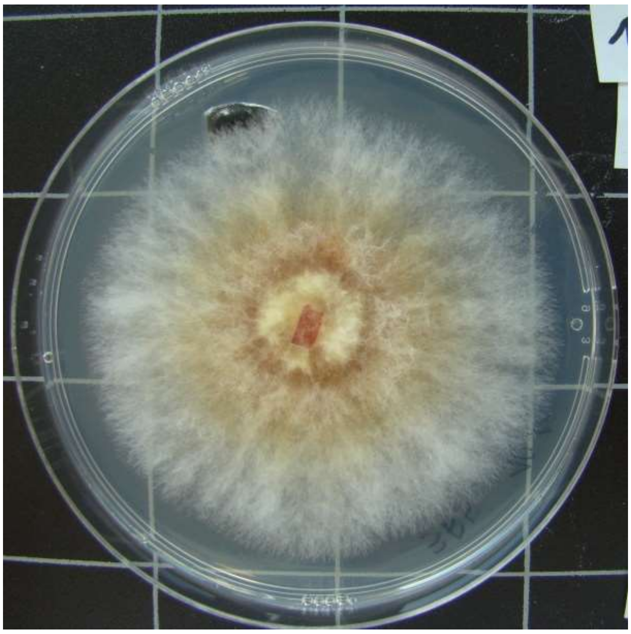

1015-HDG3.B-5

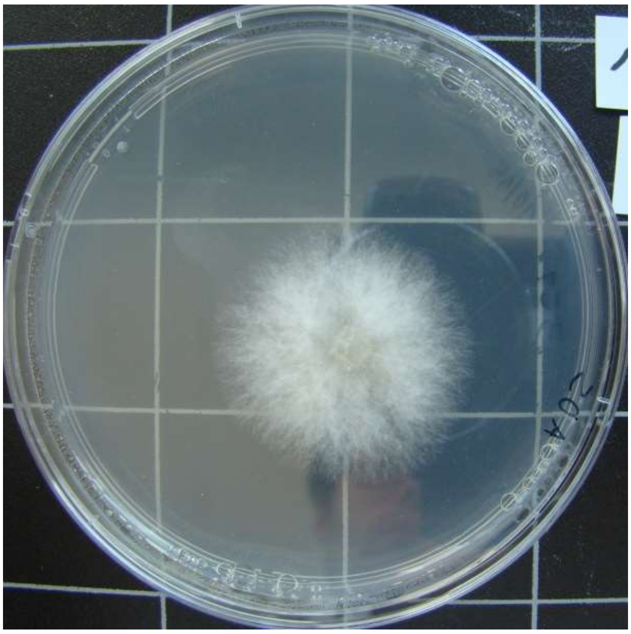

1015-HDG3.C-1

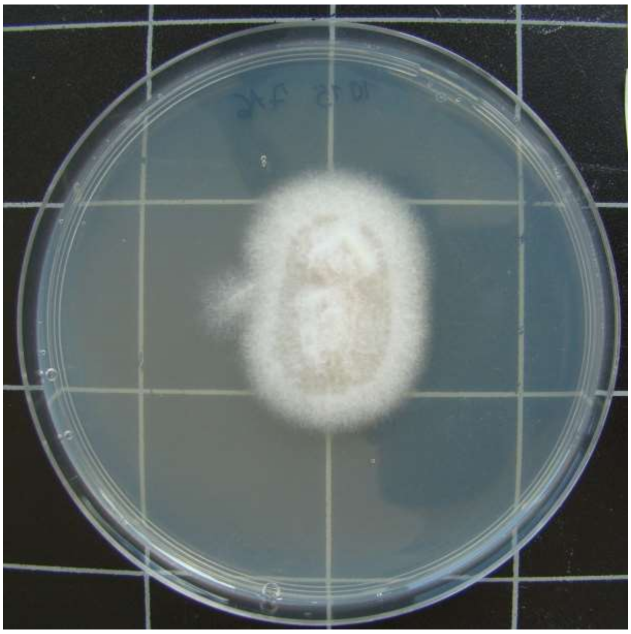

1015-TTF7.A-6

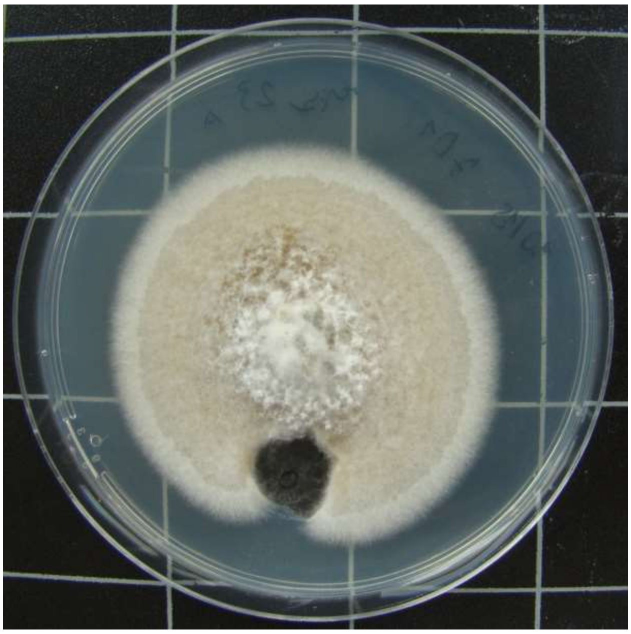

1015-TTF7.D-1

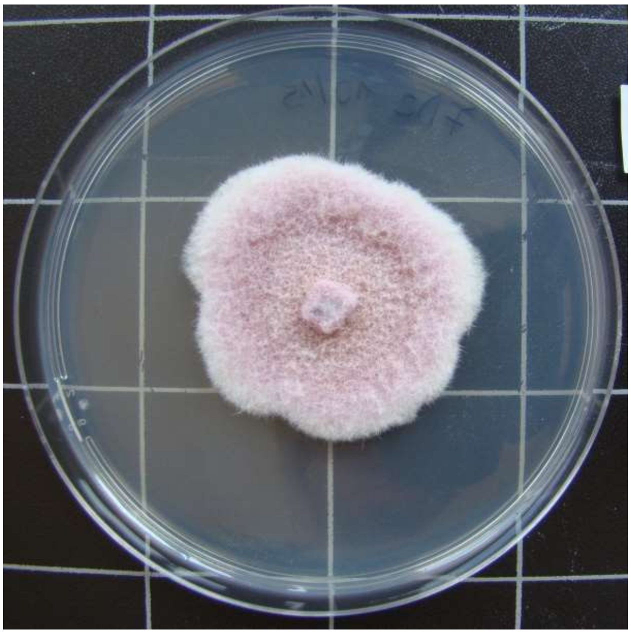

1015-TTF7.D-2

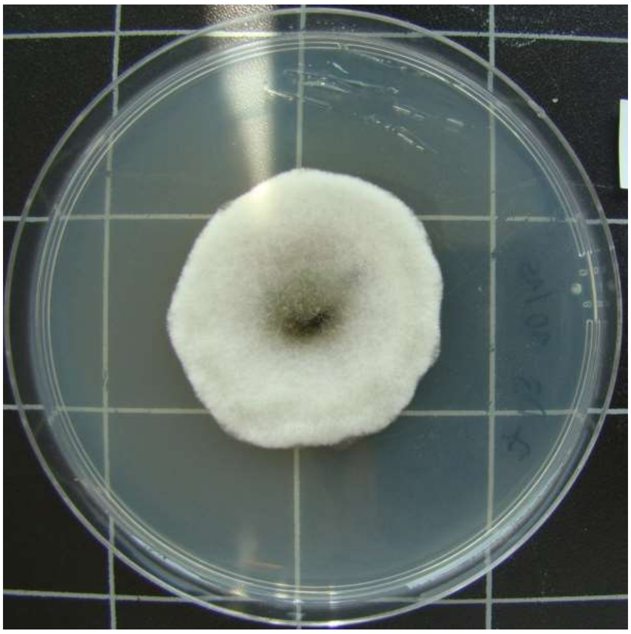

1015-TTF7.D-3

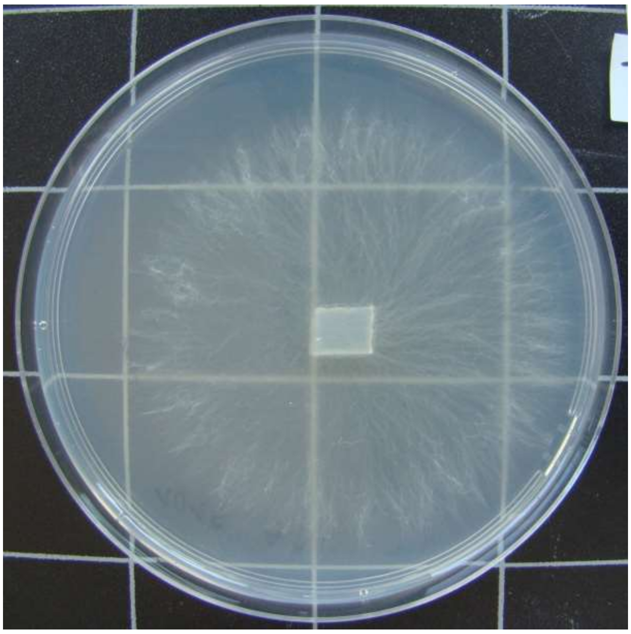

1015-TTF7.D-5

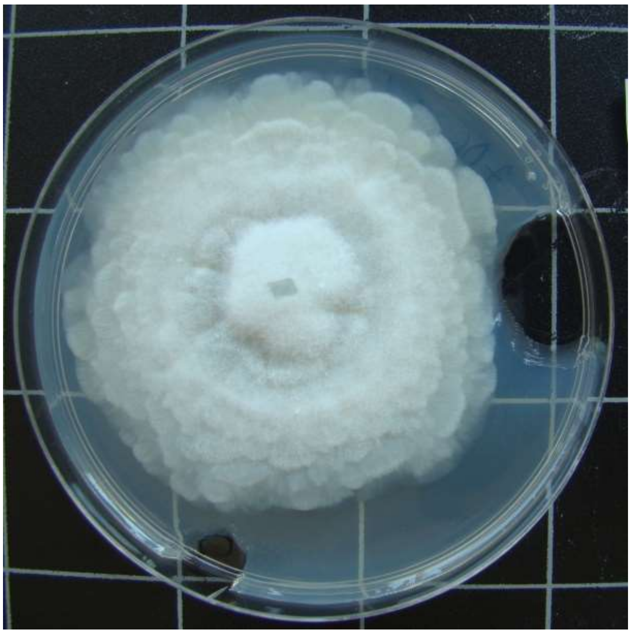

1015-TTF7.D-6

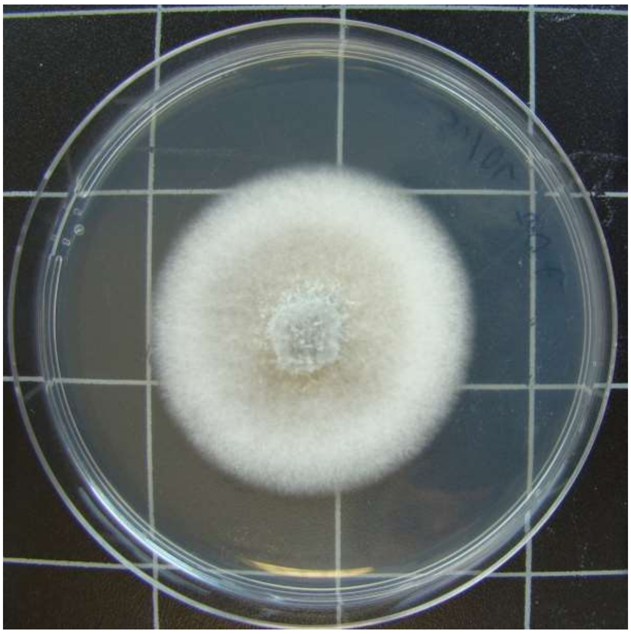

1015-TTF7.D-7

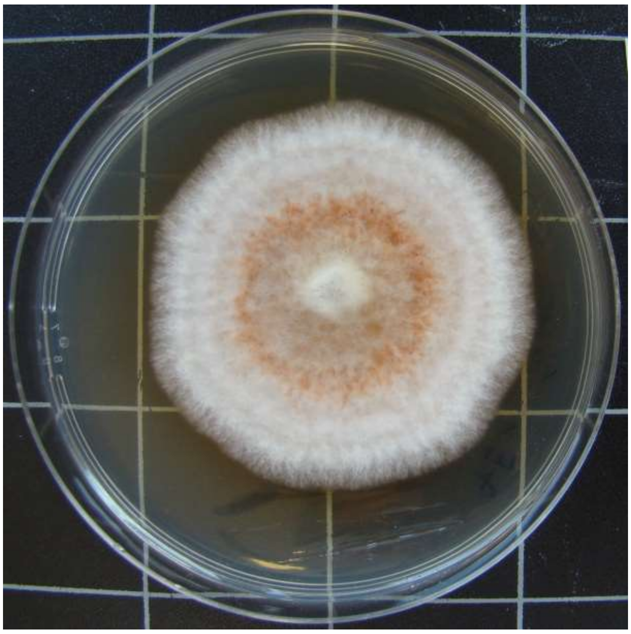

1015-TTF7.E-1

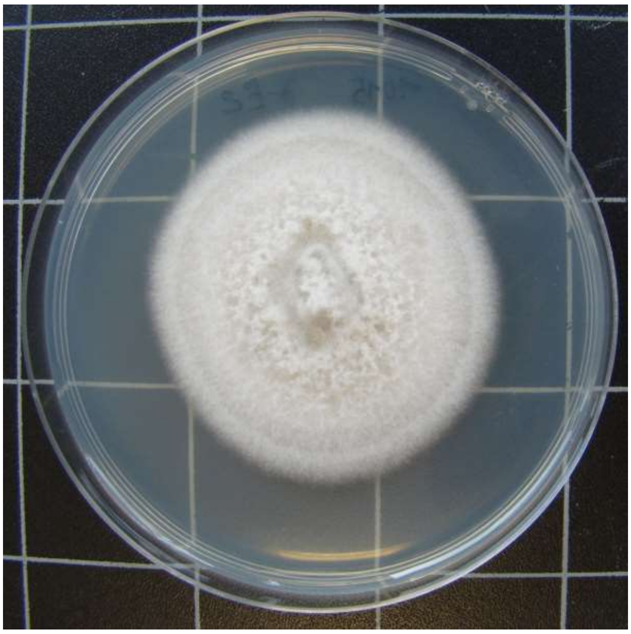

1015-TTF7.E-2

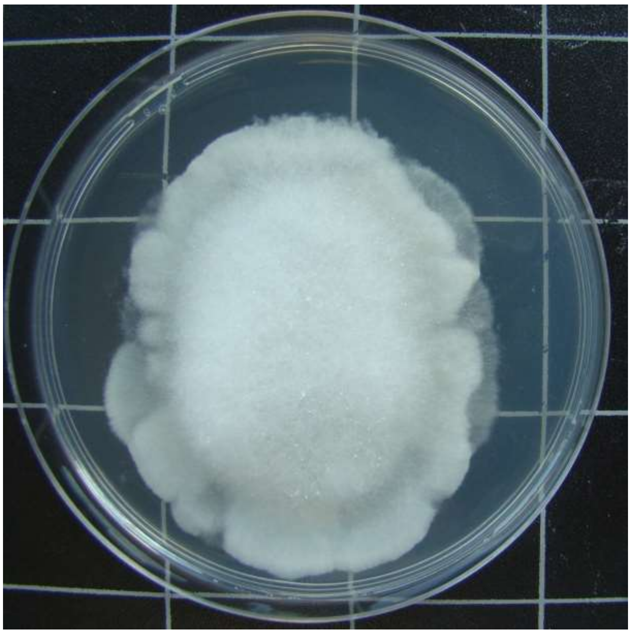

1015-TTF7.E-3

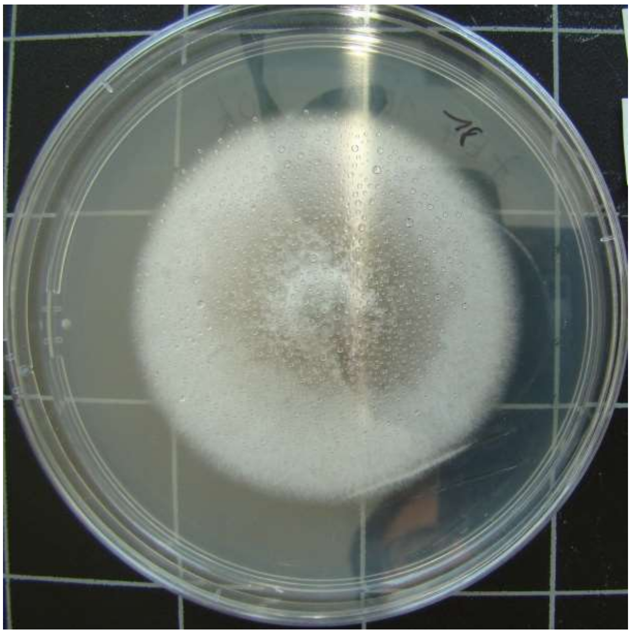

1015-TTF7.E-4

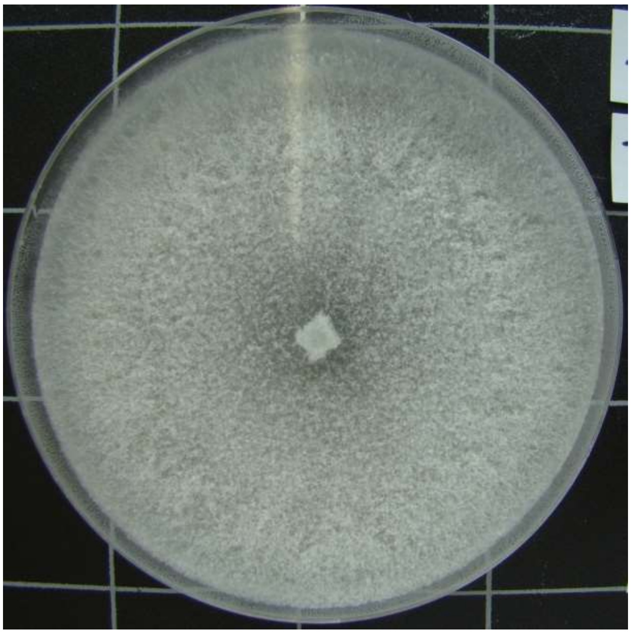

1015-TTF7.E-5

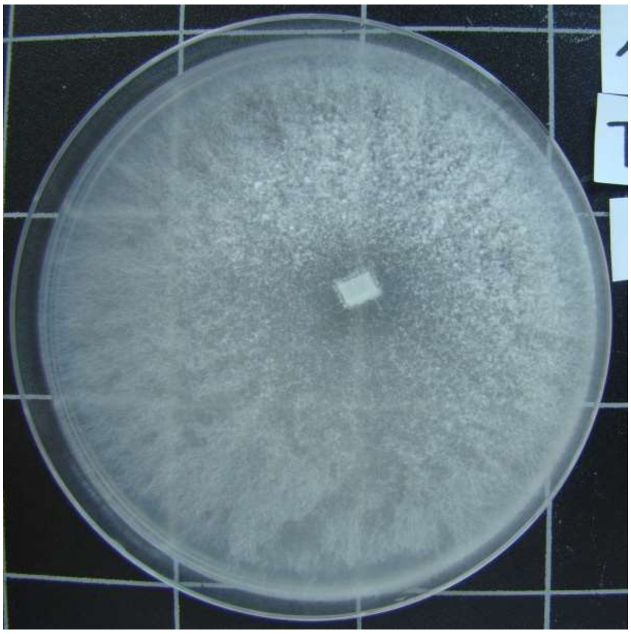

1015-TTF7.E-6

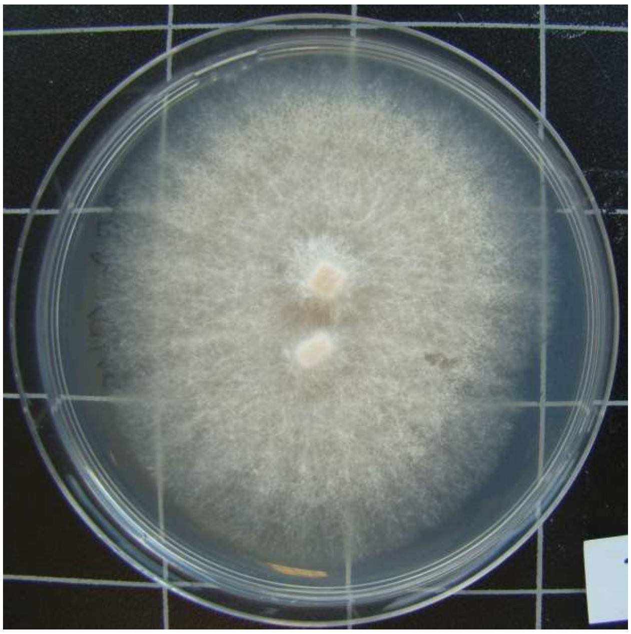

1015-TTF7.E-7

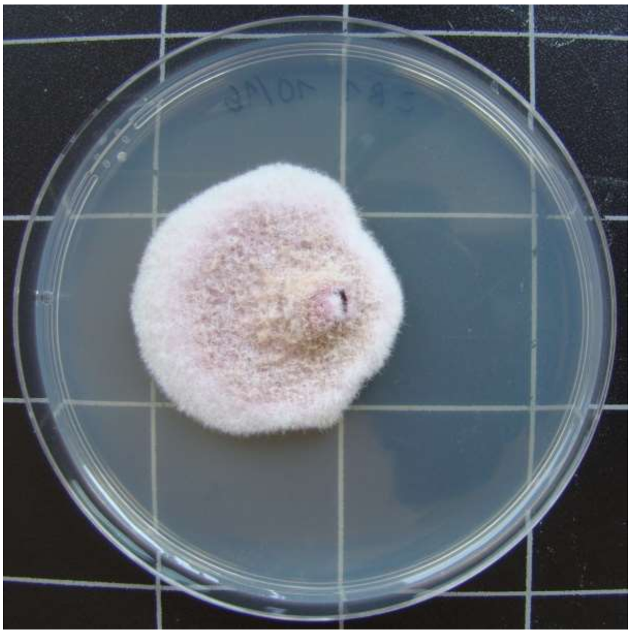

1016-HDG2.B-1

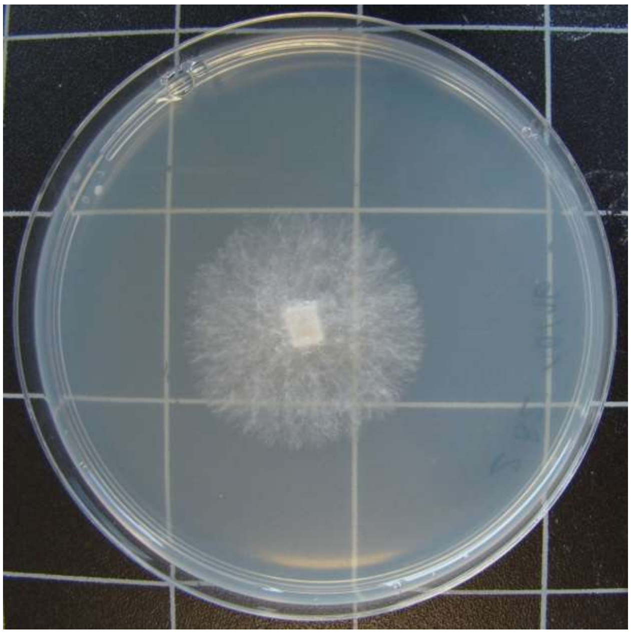

1016-HDG2.B-2

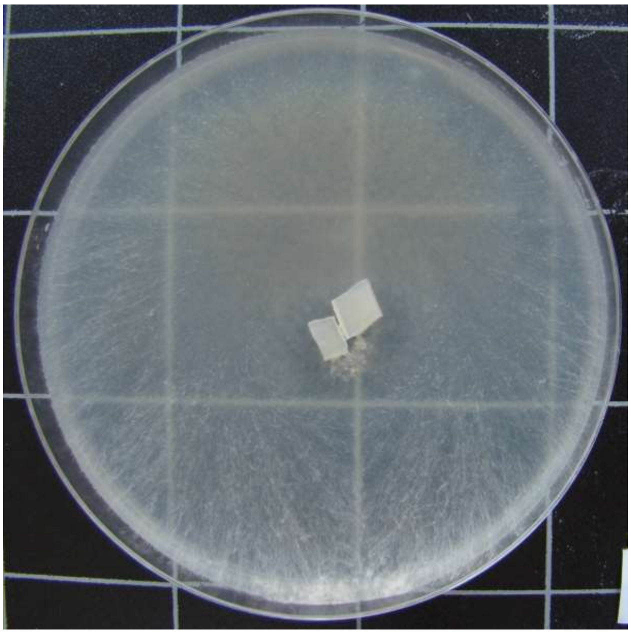

1016-HDG2.B-4

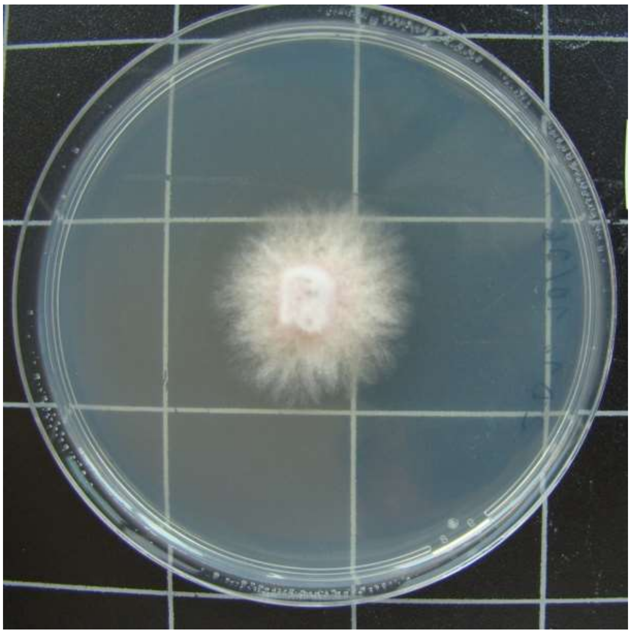

1016-HDG2.D-1

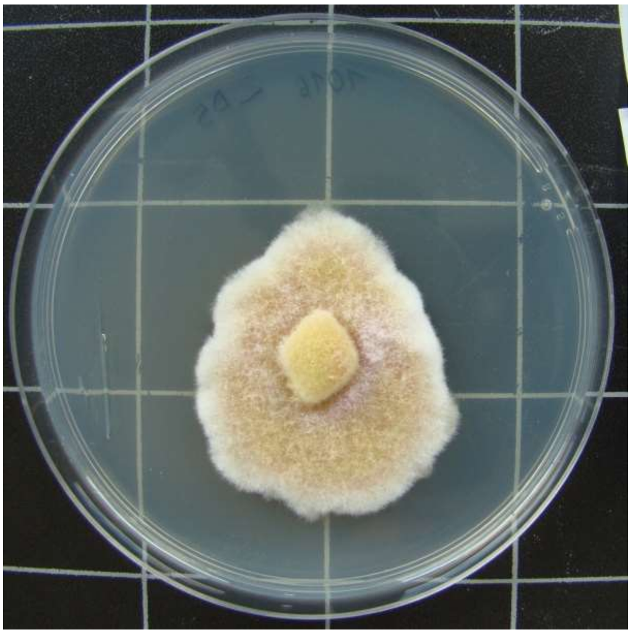

1016-HDG2.D-5

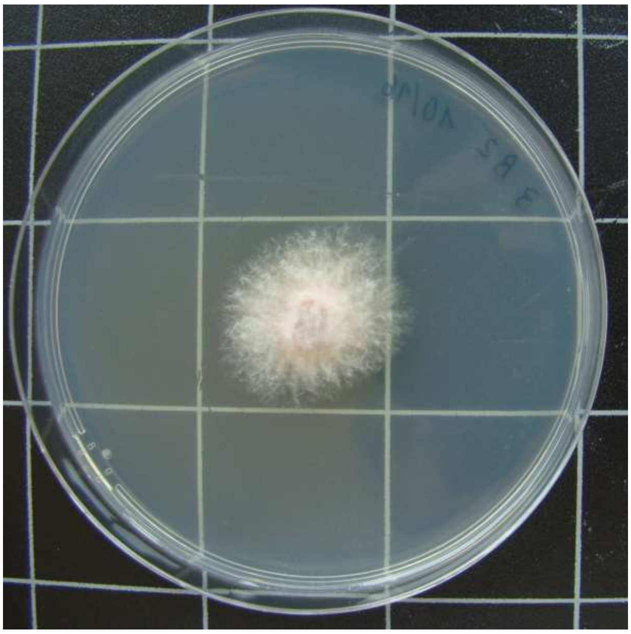

1016-HDG3.B-2

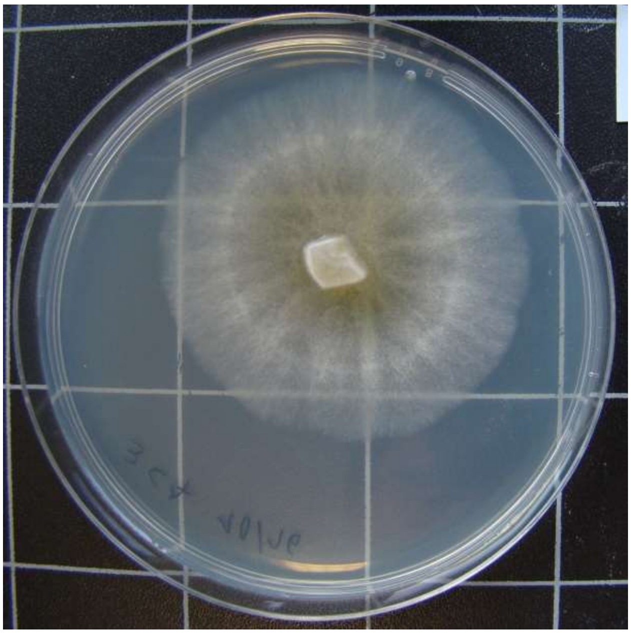

1016-HDG3.C-7

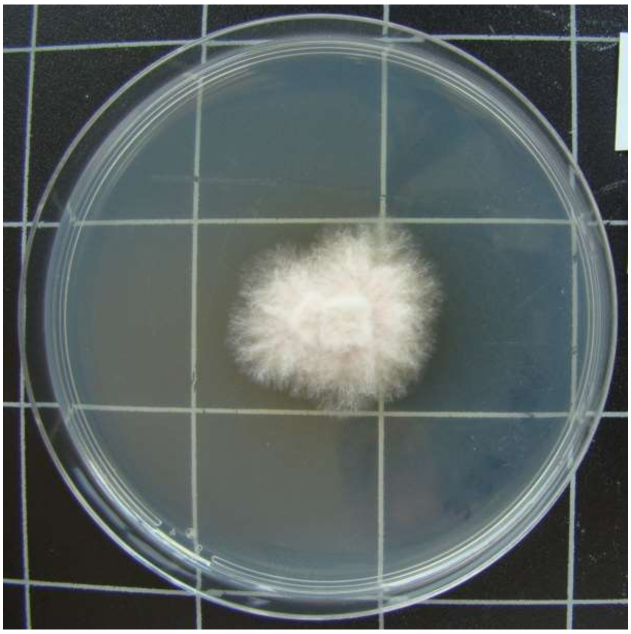

1016-HDG3.E-1

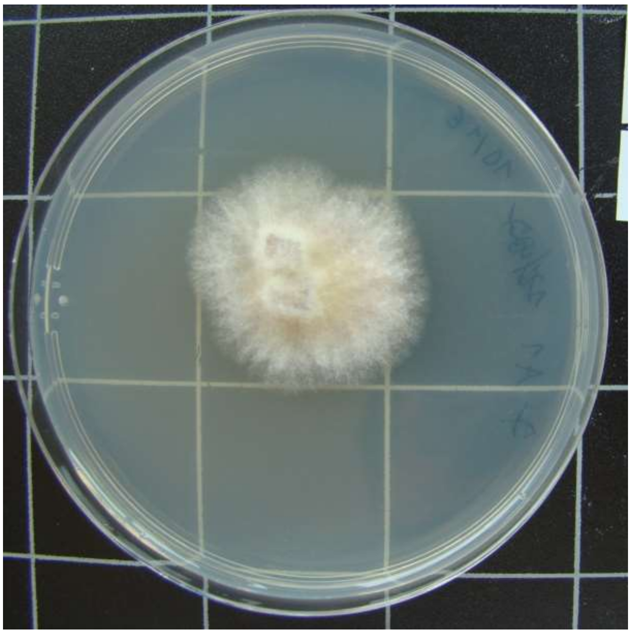

1016-TTF7.A-1

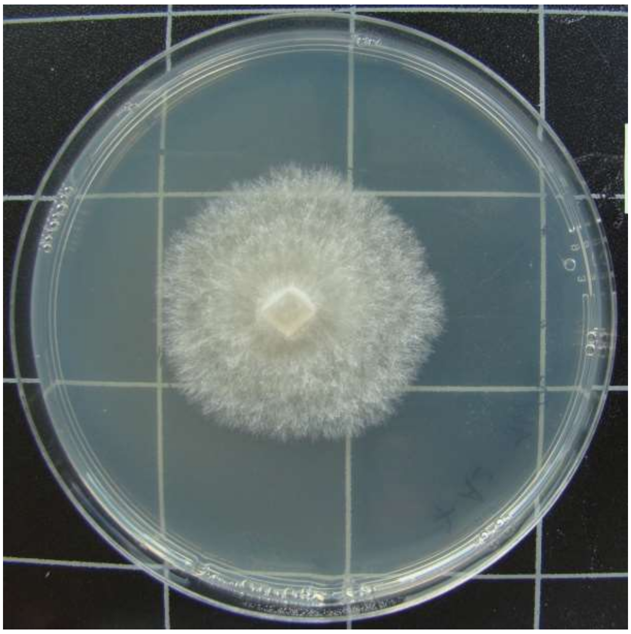

1016-TTF7.A-2

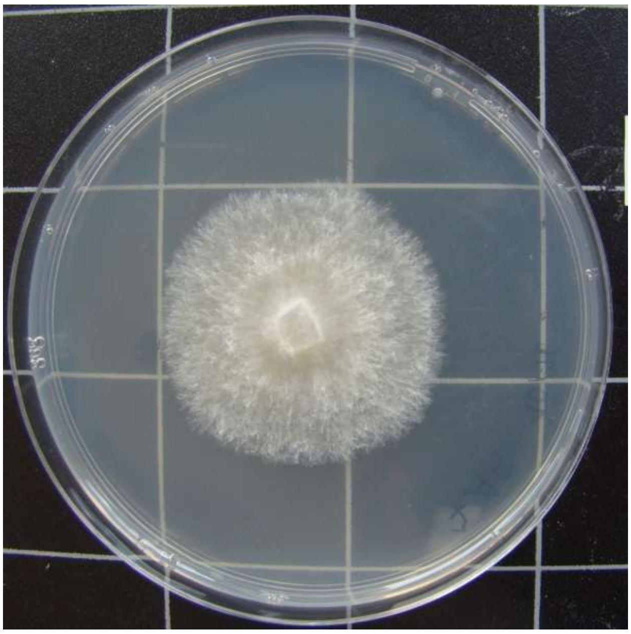

1016-TTF7.A-5

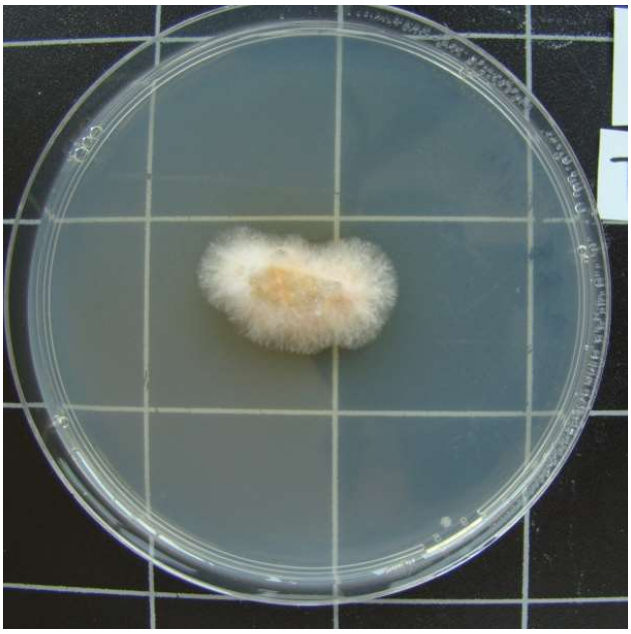

1016-TTF7.B-8

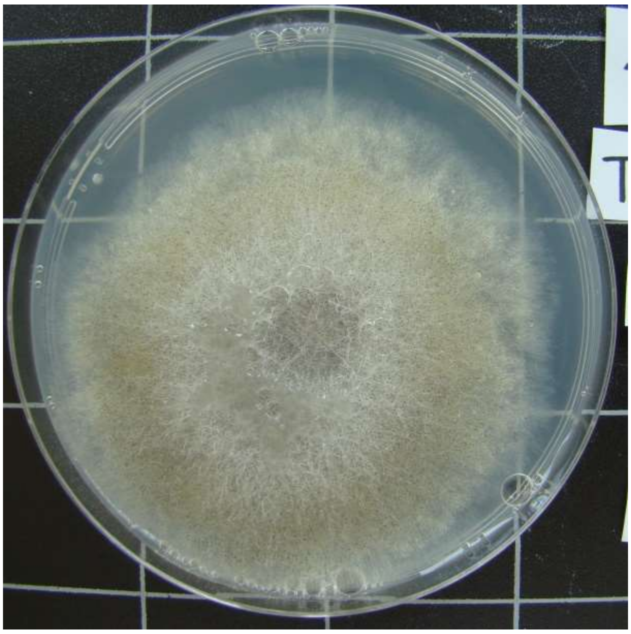

1016-TTF7.C-4

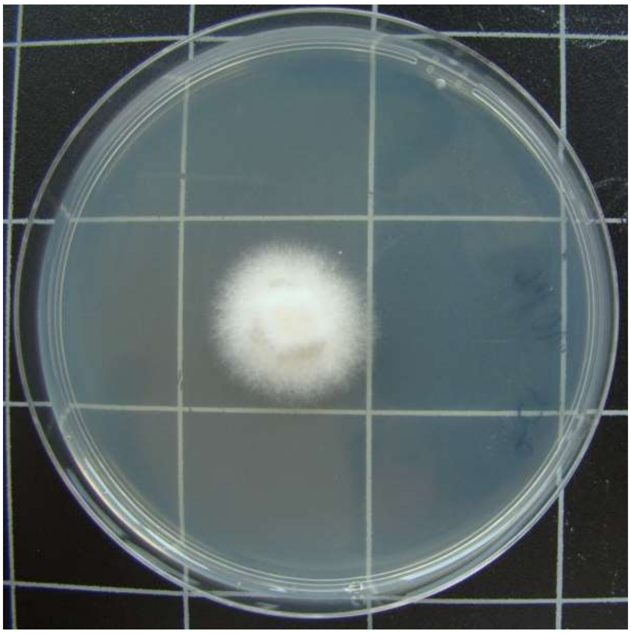

1016-TTF7.C-8

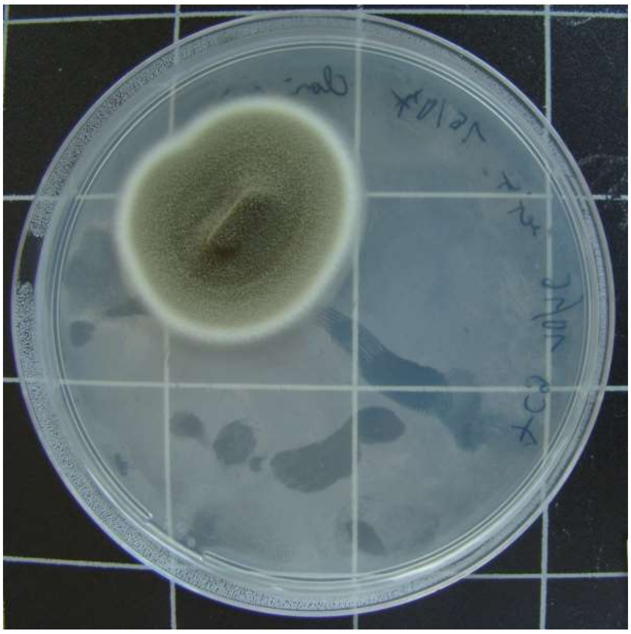

1016-TTF7.C-9

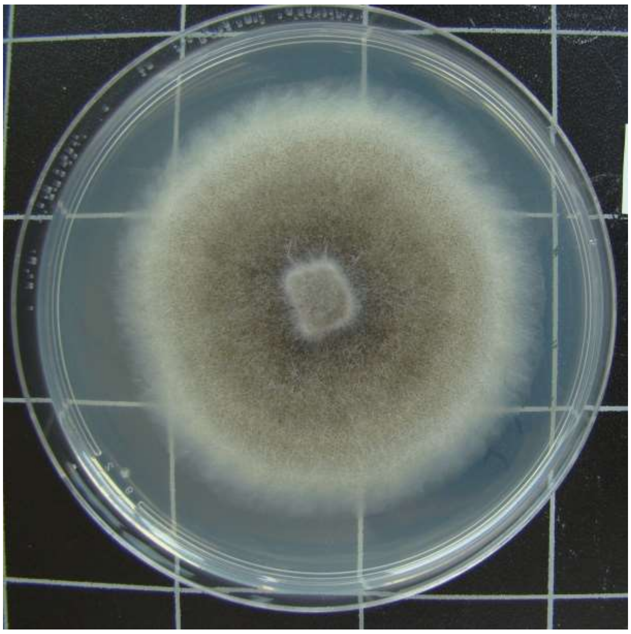

1016-TTF7.D-3

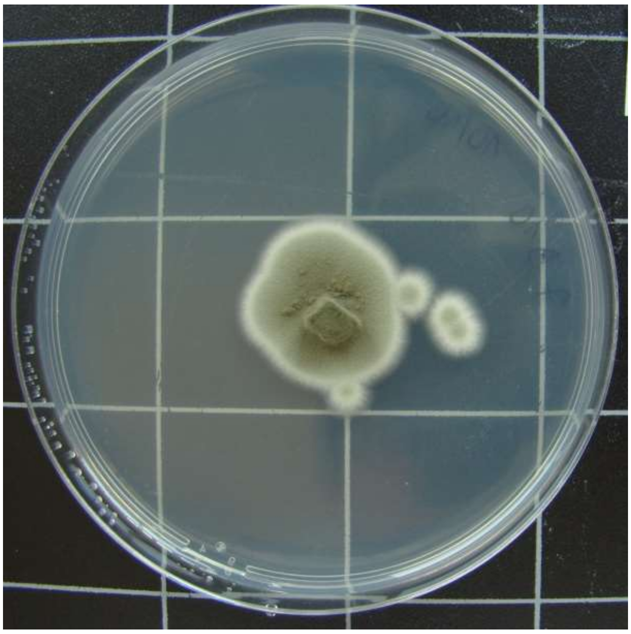

1016-TTF7.D-10

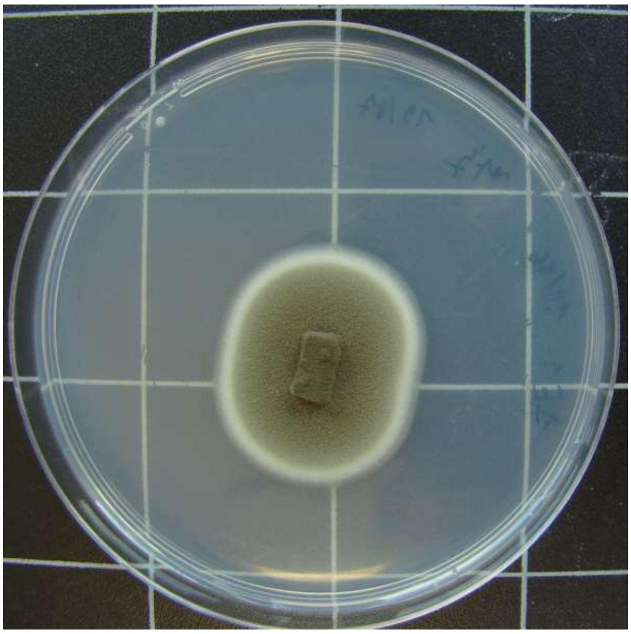

1016-TTF7.E-1

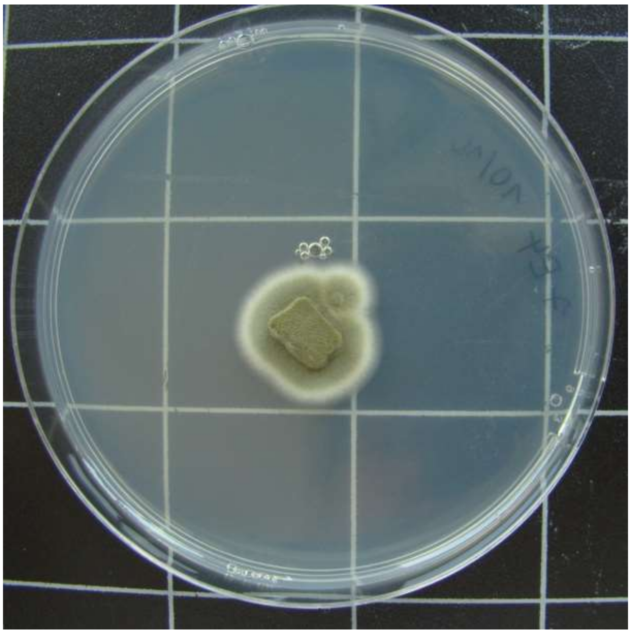

1016-TTF7.E-4

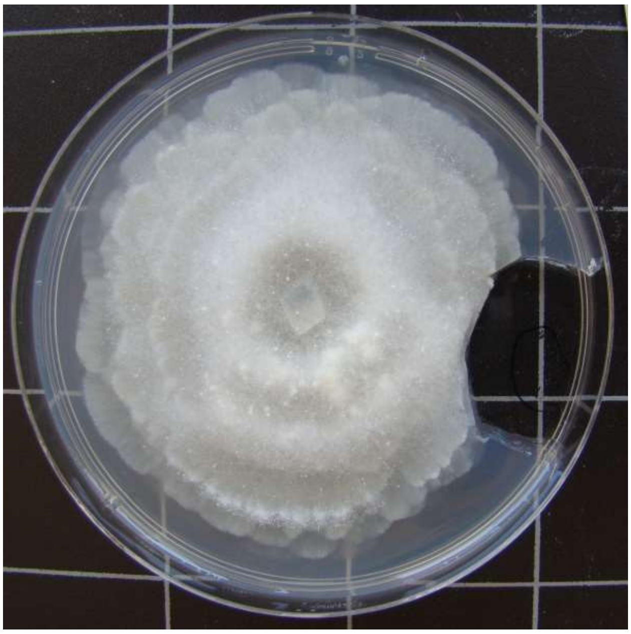

1215-HDG2.A-2

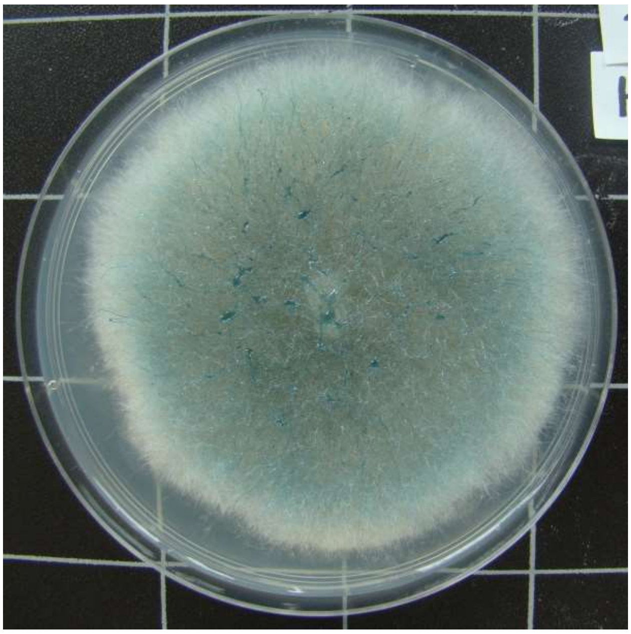

1215-HDG2.A-3

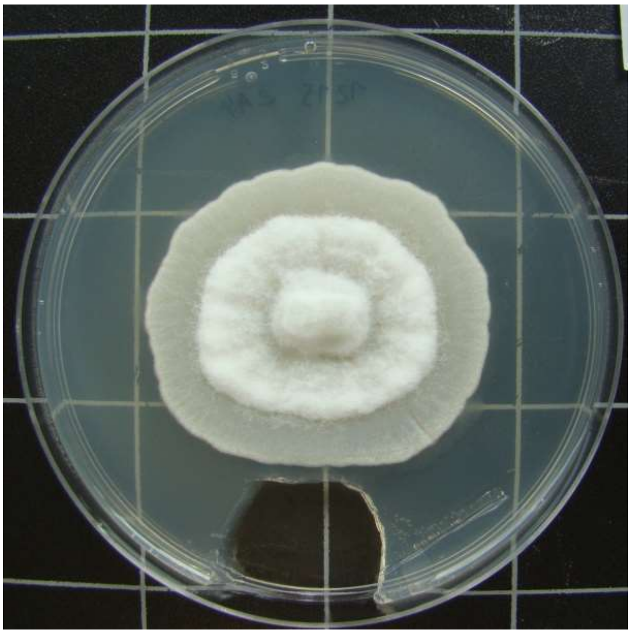

1215-HDG2.A-4

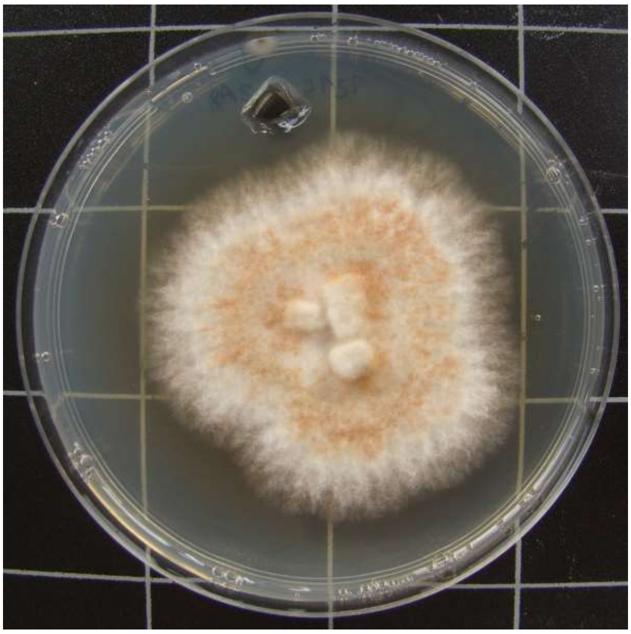

1215-HDG2.A-8

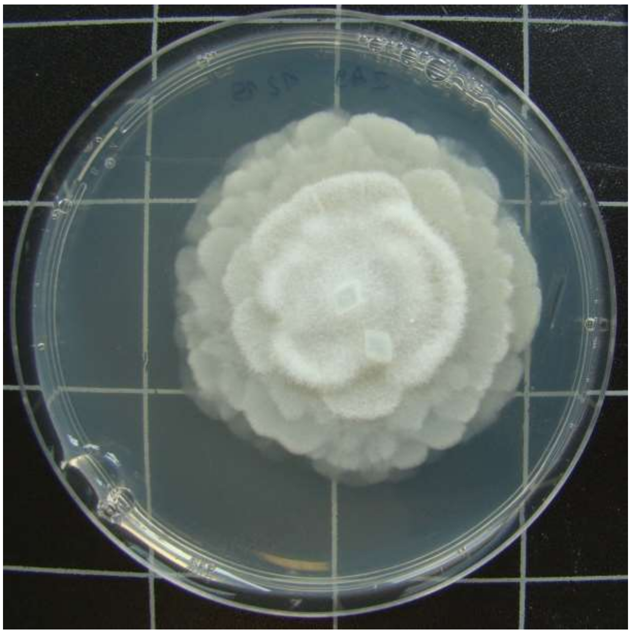

1215-HDG2.A-9

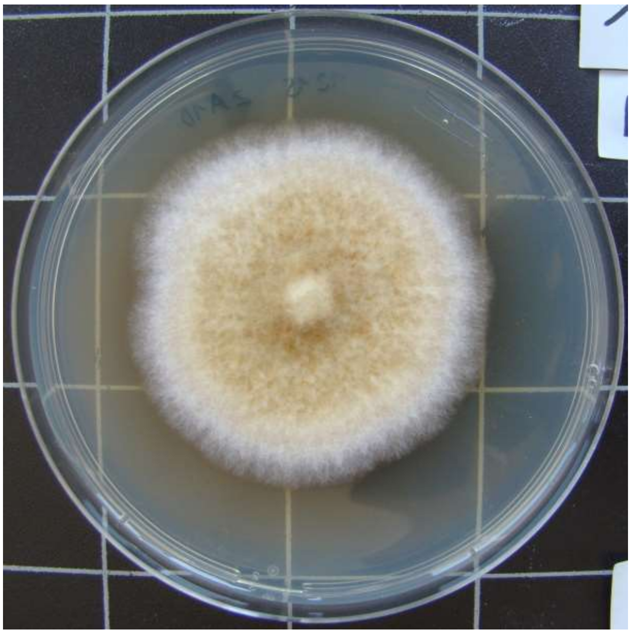

1215-HDG2.A-10

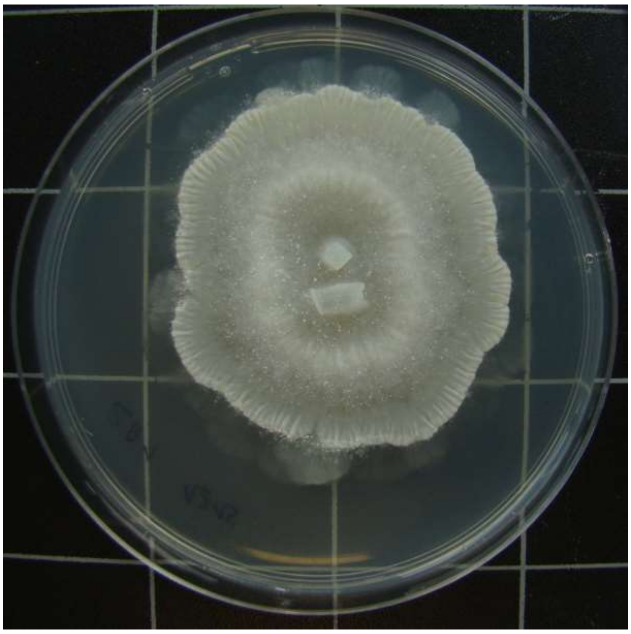

1215-HDG2.B-1

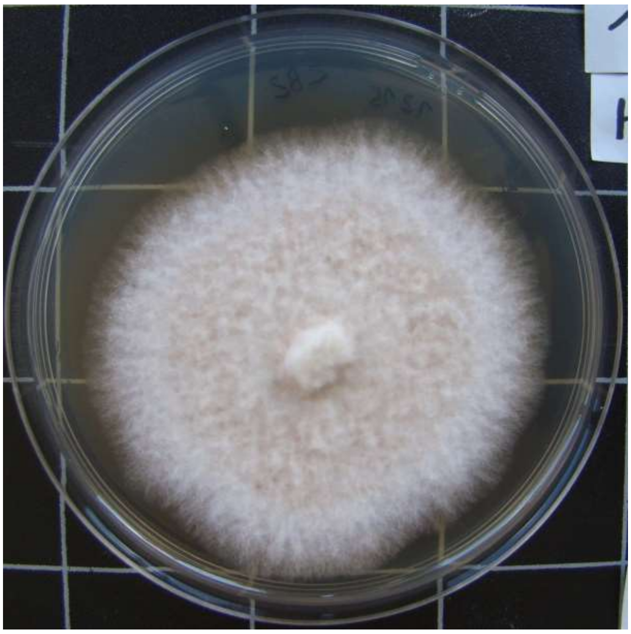

1215-HDG2.B-2

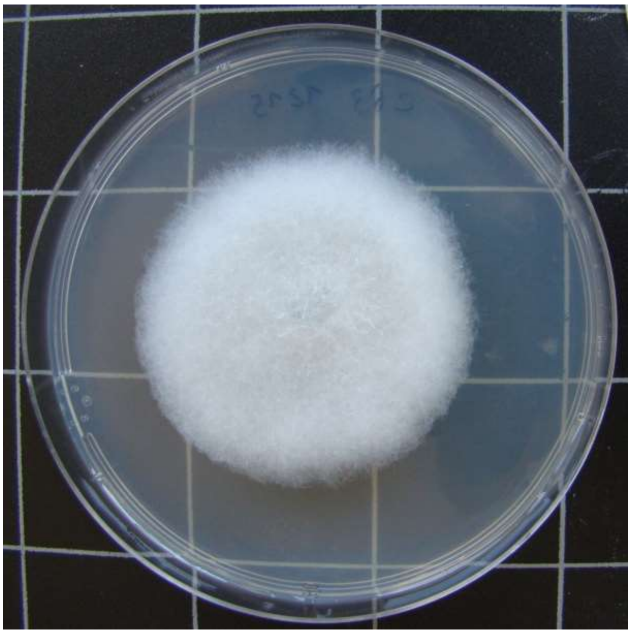

1215-HDG2.B-3

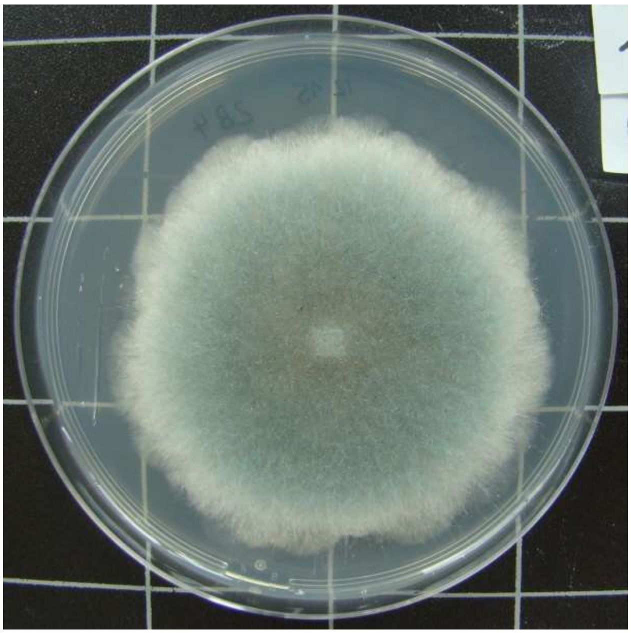

1215-HDG2.B-4

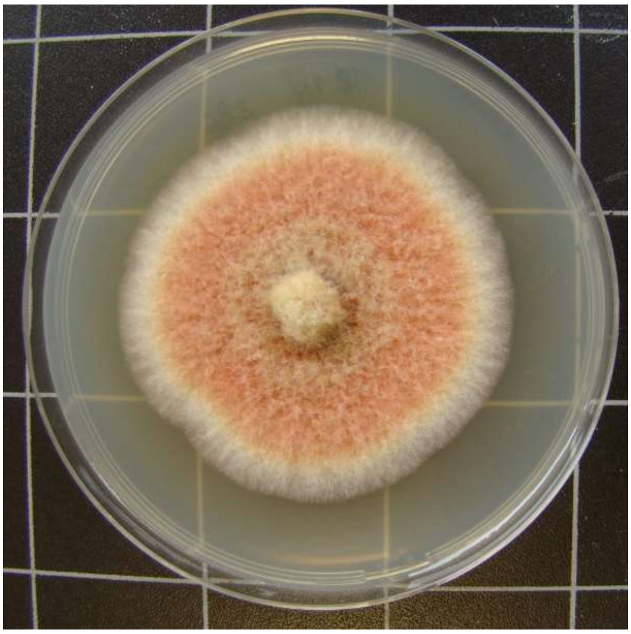

1215-HDG2.B-6

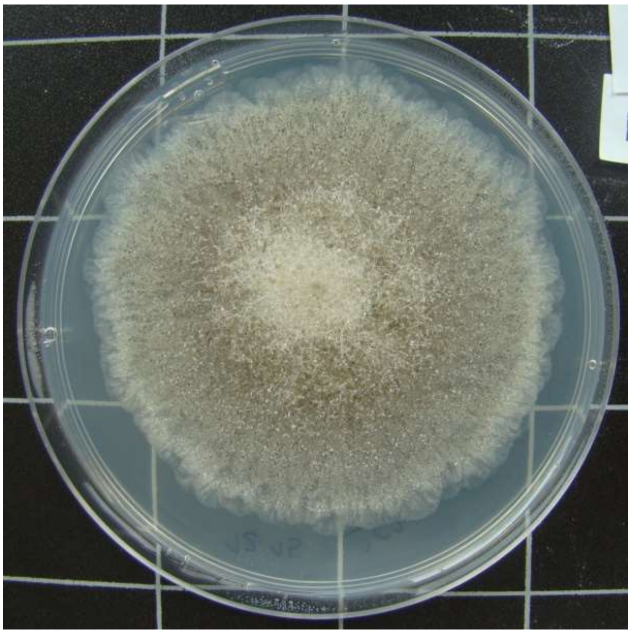

1215-HDG2.C-1

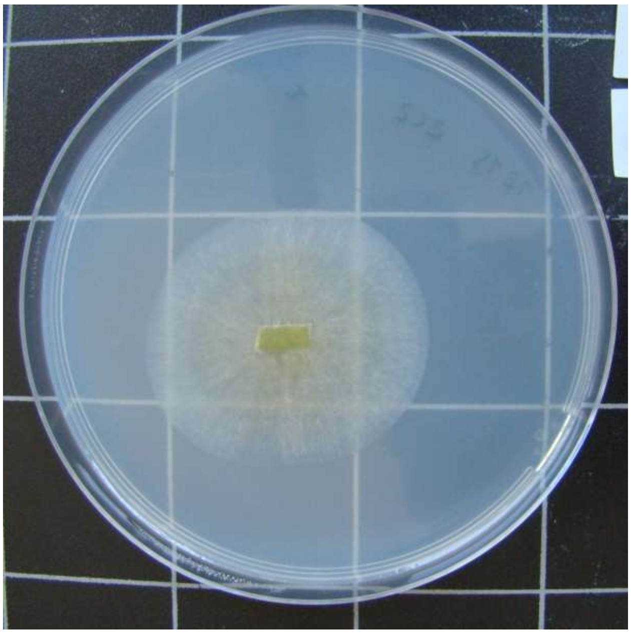

1215-HDG2.C-2

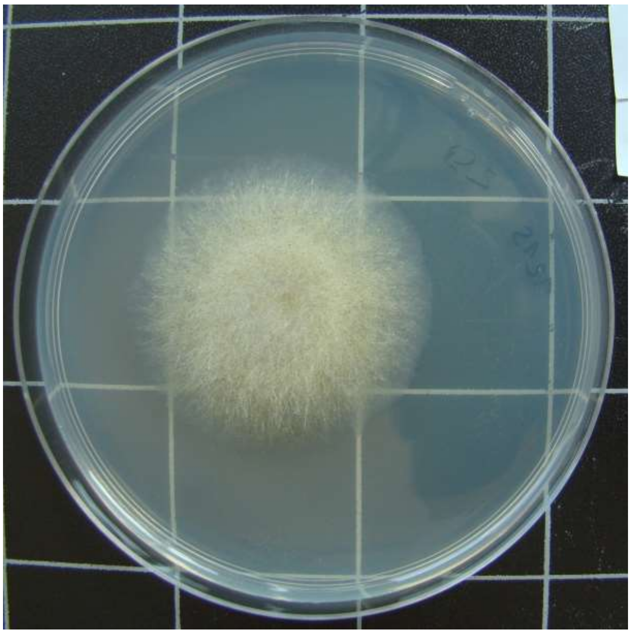

1215-HDG2.C-3

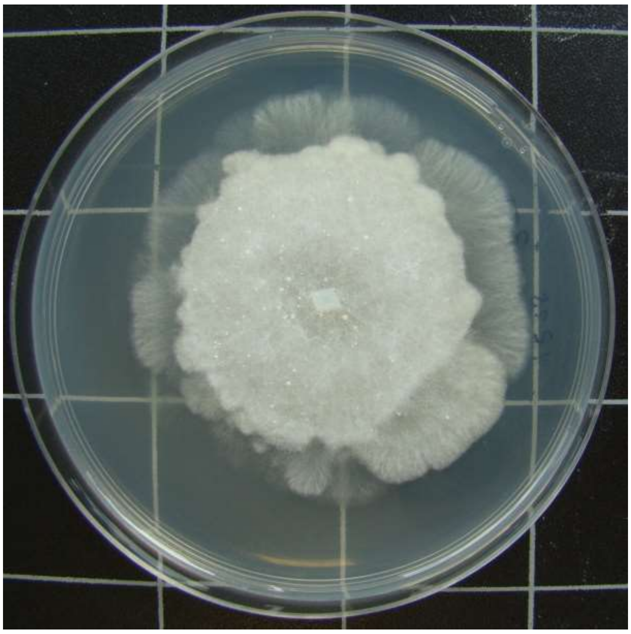

1215-HDG2.C-4

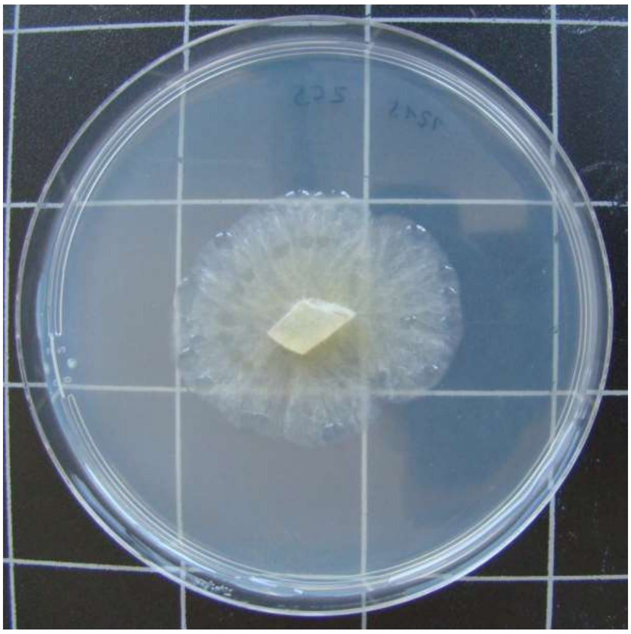

1215-HDG2.C-5

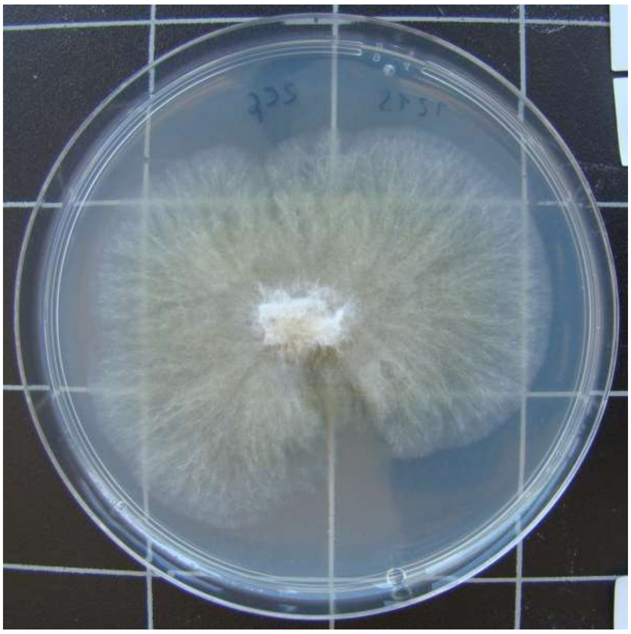

1215-HDG2.C-6

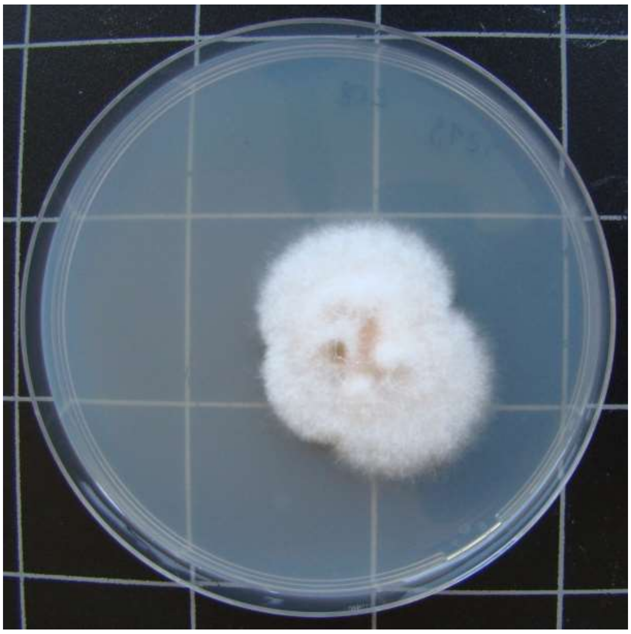

1215-HDG2.C-8

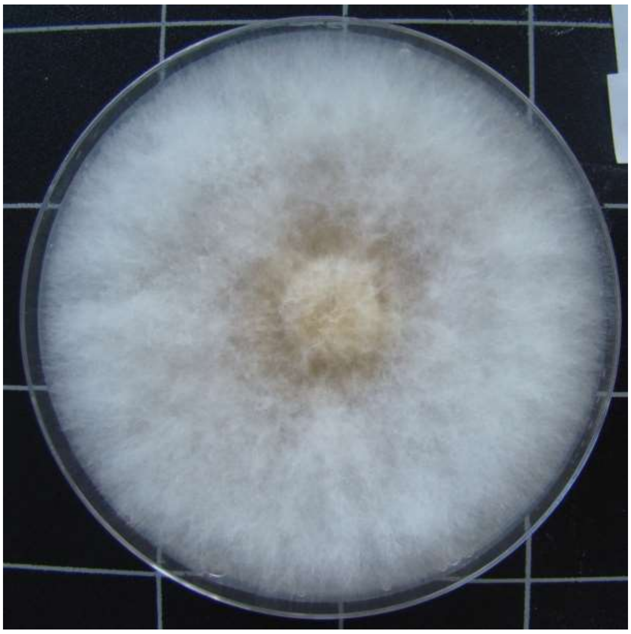

1215-HDG2.D-2

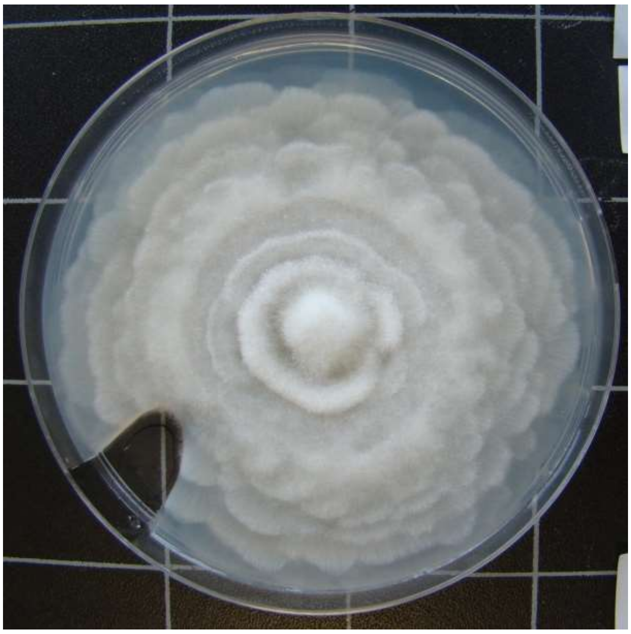

1215-HDG2.D-3

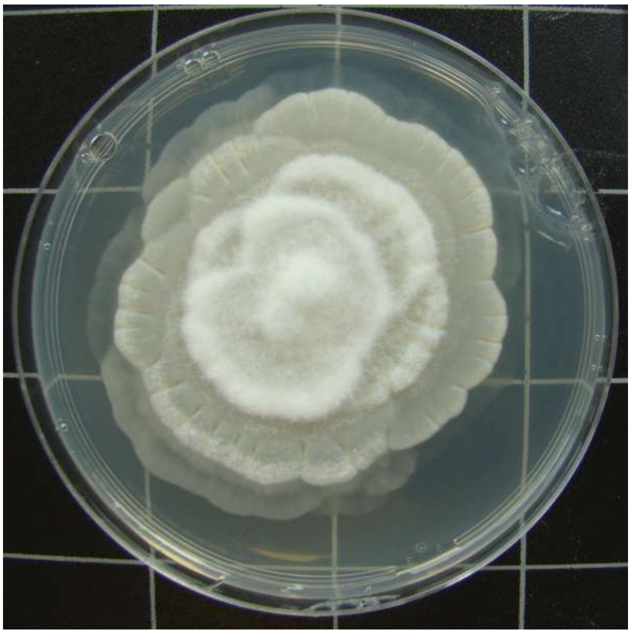

1215-HDG2.D-6

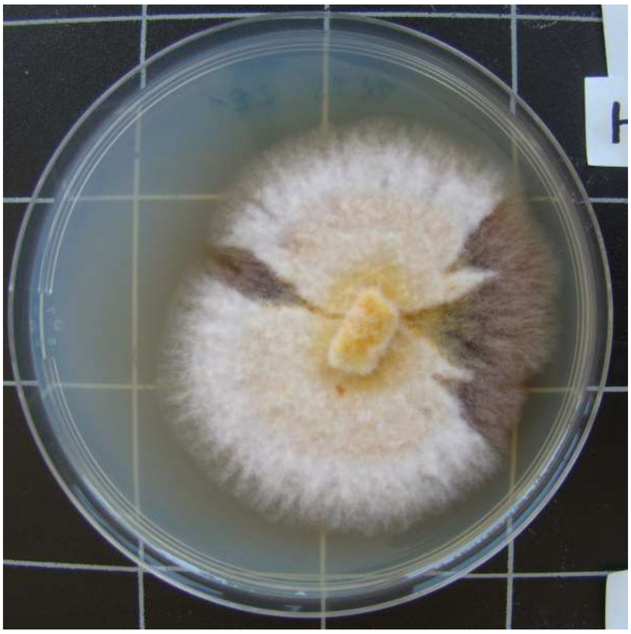

1215-HDG2.E-1

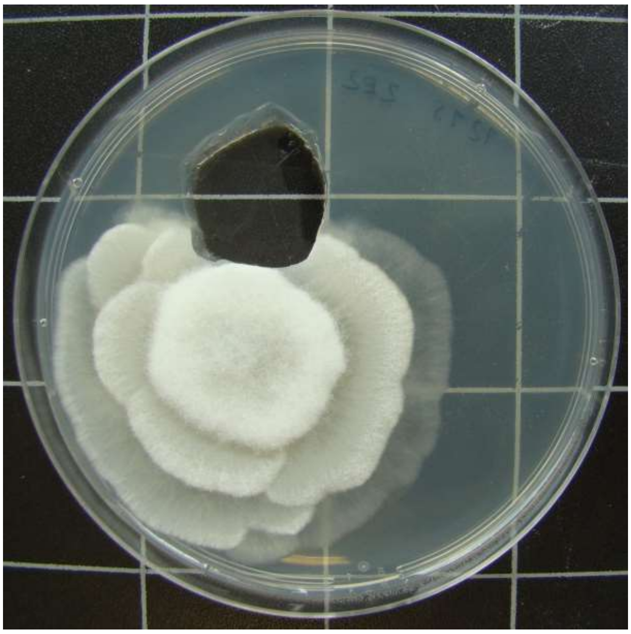

1215-HDG2.E-2

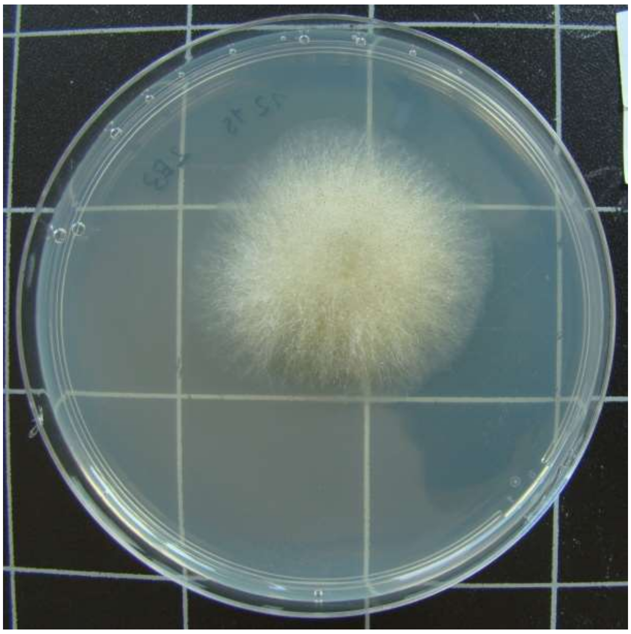

1215-HDG2.E-3

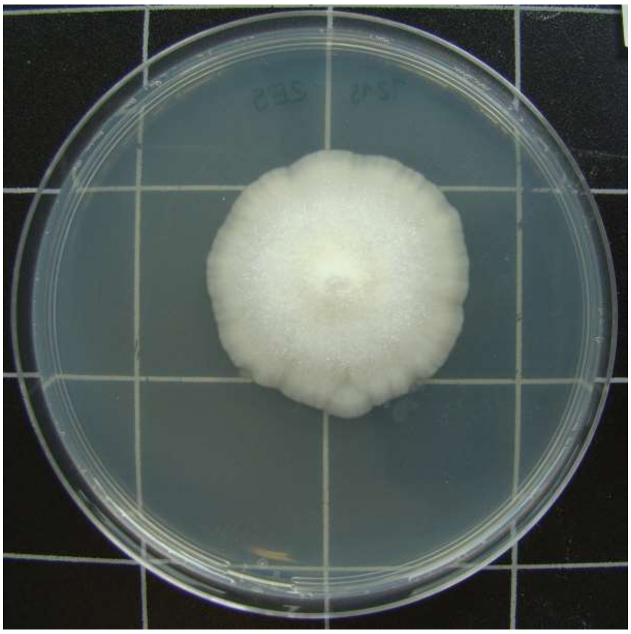

1215-HDG2.E-5

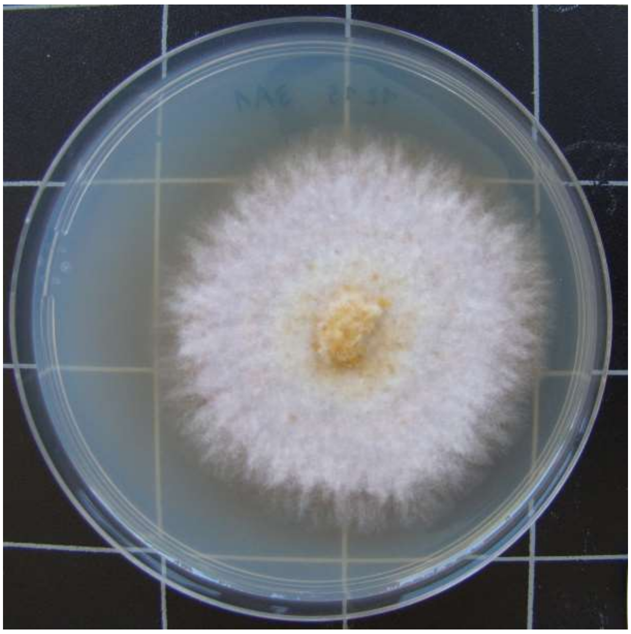

1215-HDG3.A-1

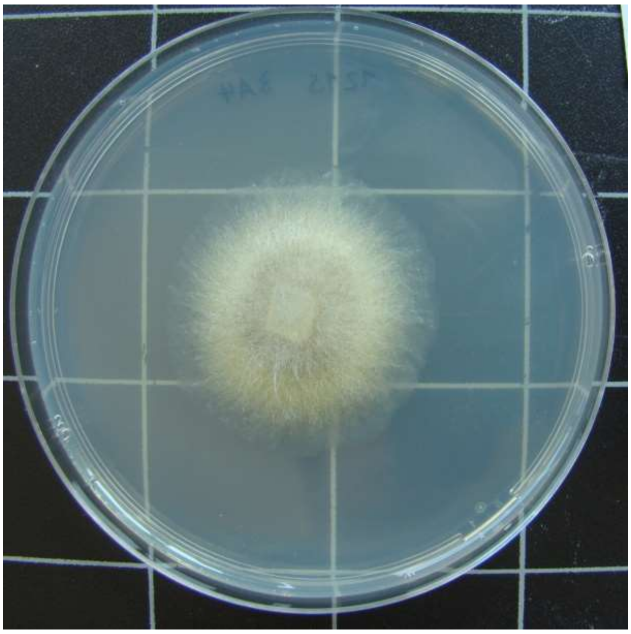

1215-HDG3.A-4

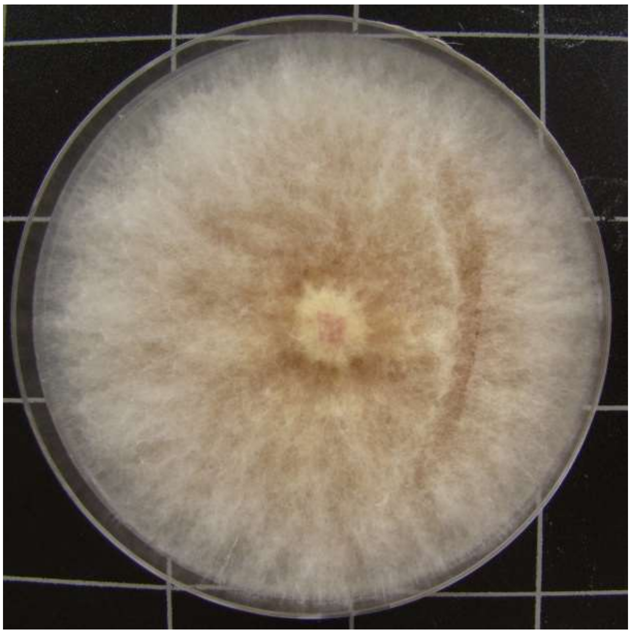

1215-HDG3.A-5

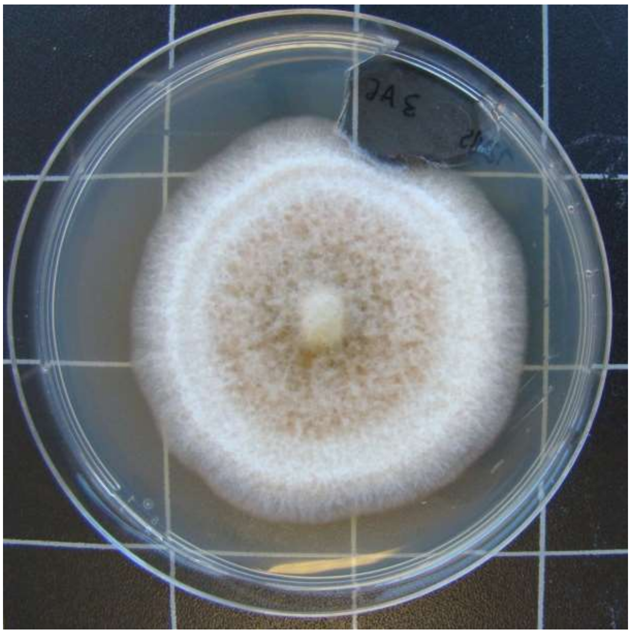

1215-HDG3.A-6

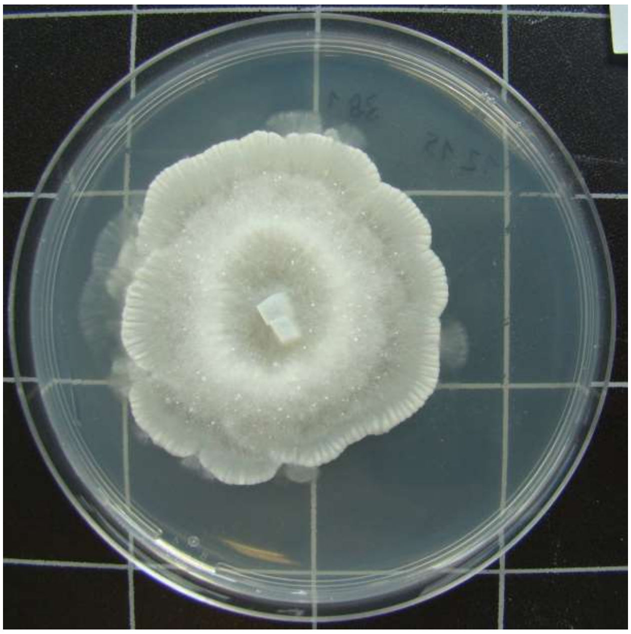

1215-HDG3.B-1

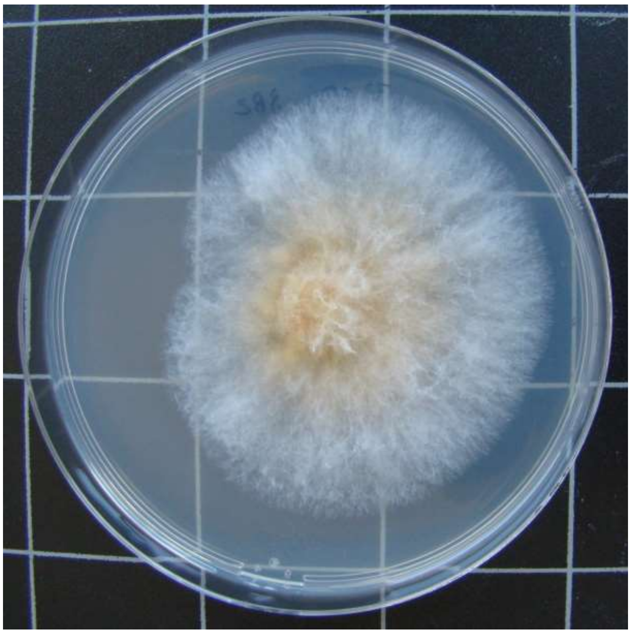

1215-HDG3.B-2

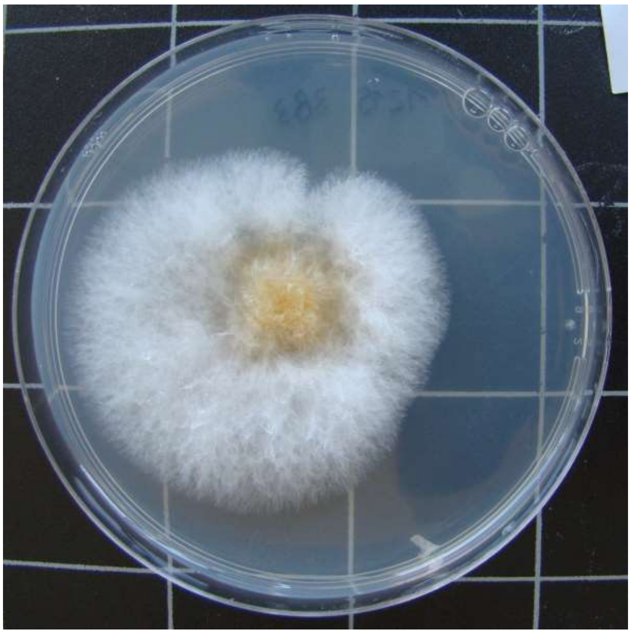

1215-HDG3.B-3

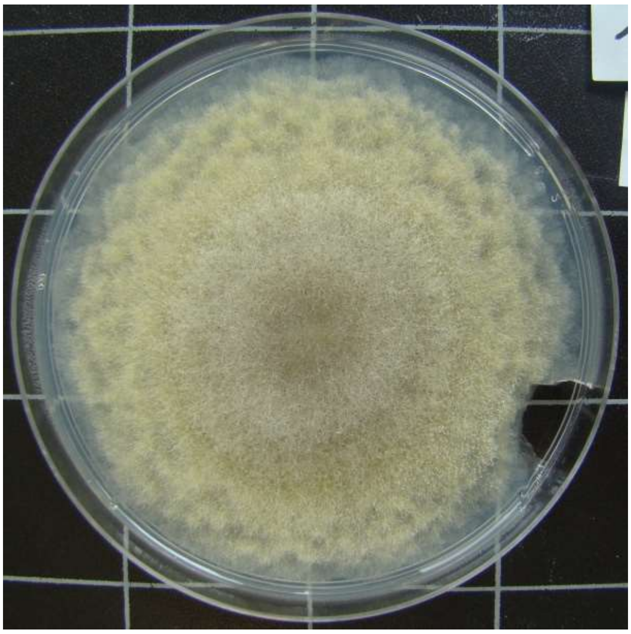

1215-HDG3.B-4

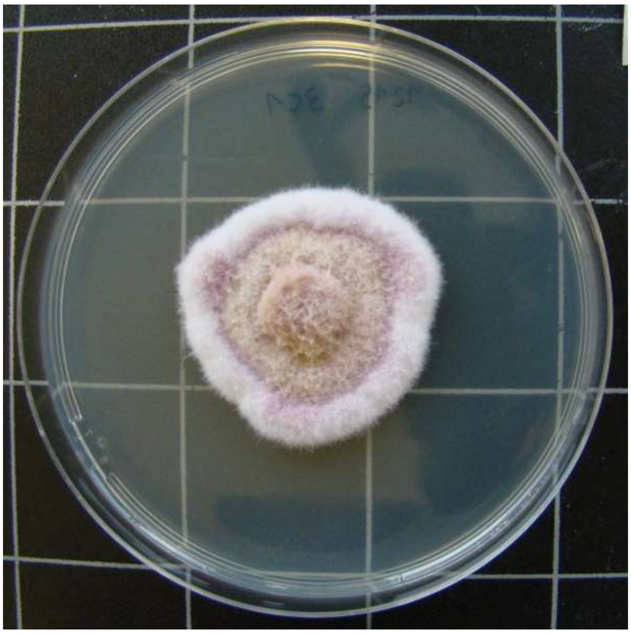

1215-HDG3.C-1

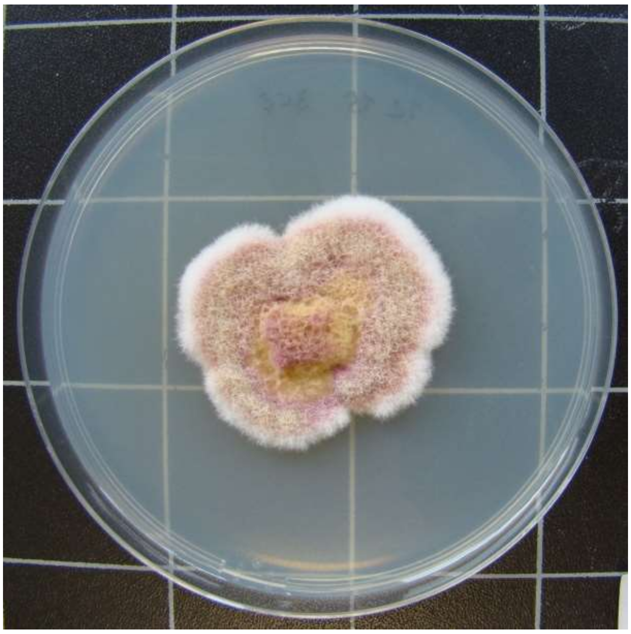

1215-HDG3.C-2

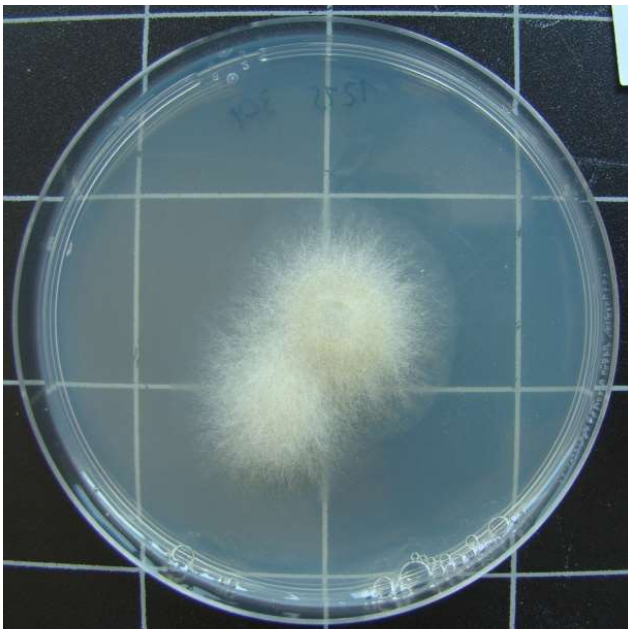

1215-HDG3.C-4

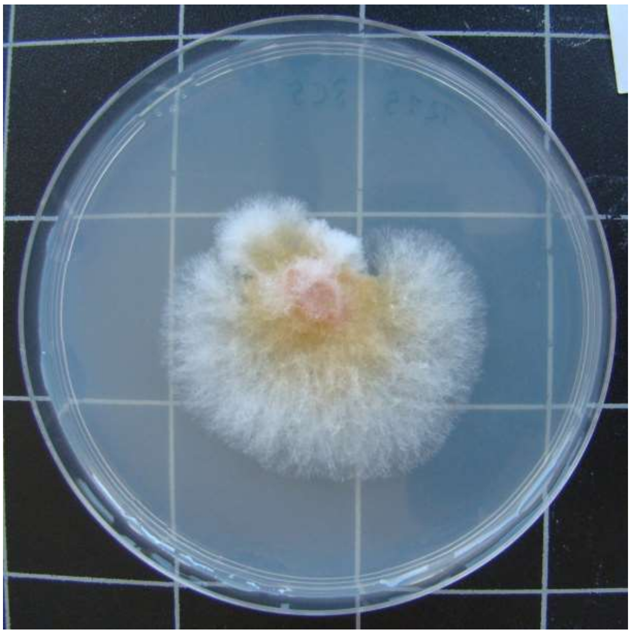

1215-HDG3.C-5

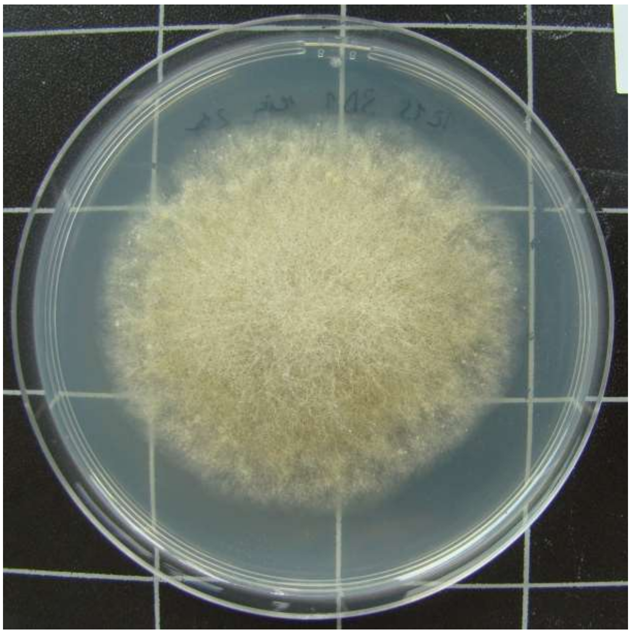

1215-HDG3.D-1

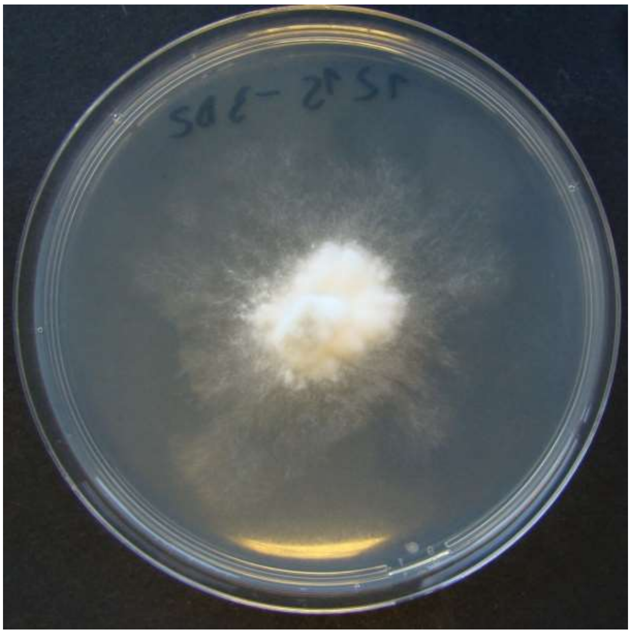

1215-HDG3.D-2

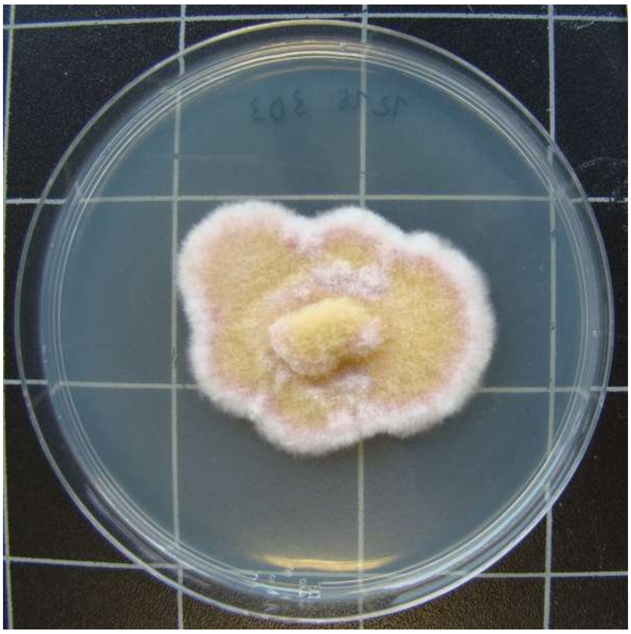

1215-HDG3.D-3

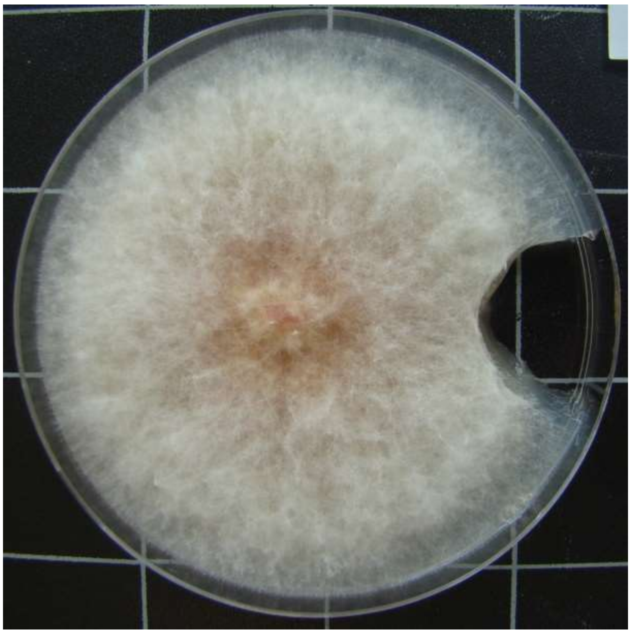

1215-HDG3.D-4

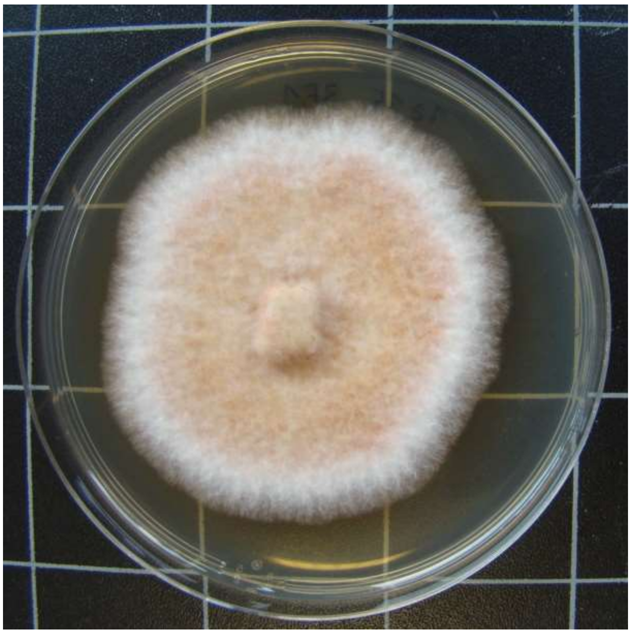

1215-HDG3.E-1

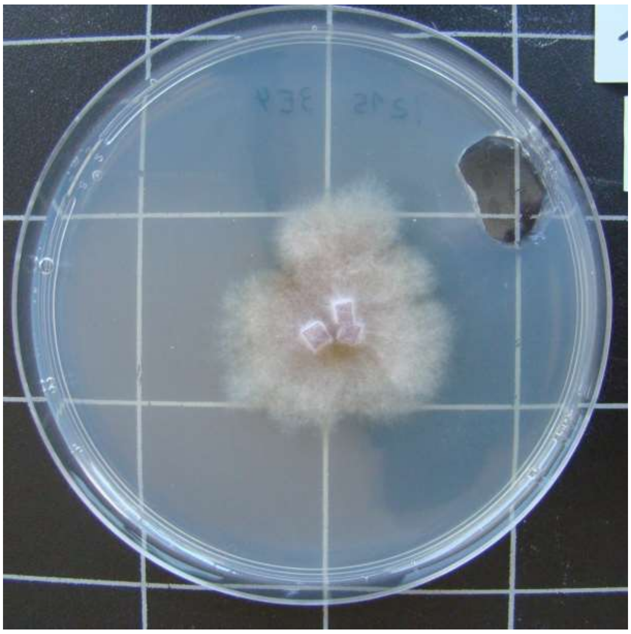

1215-HDG3.E-4

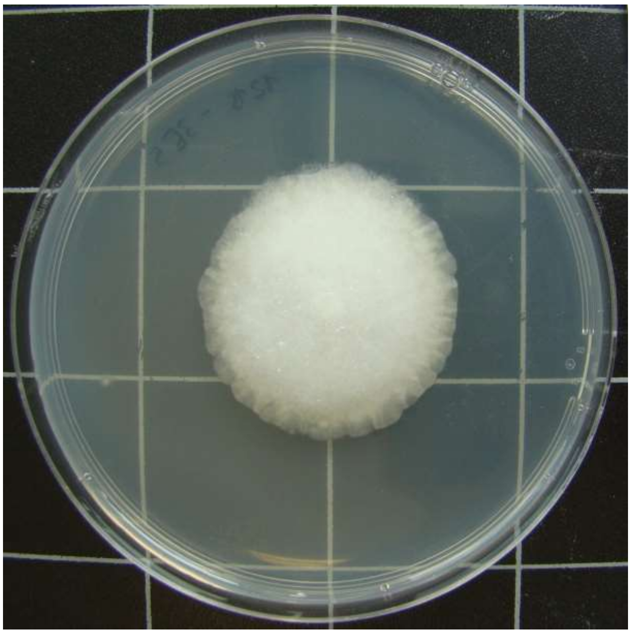

1215-HDG3.E-5

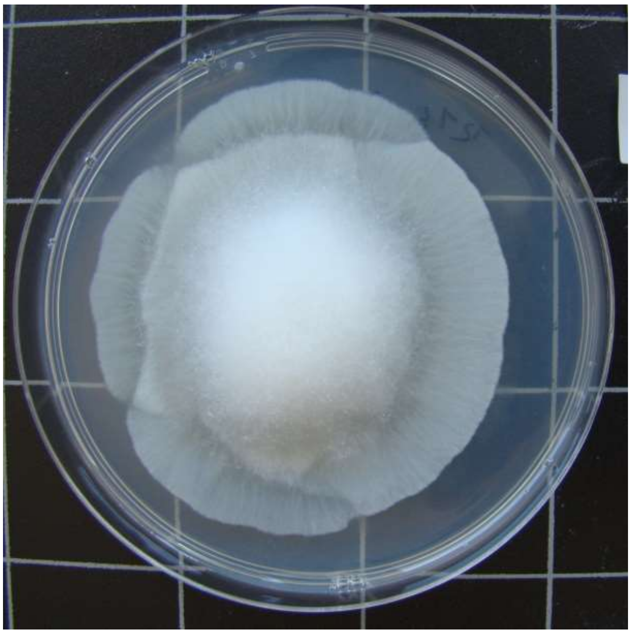

1215-TTF7.A-2

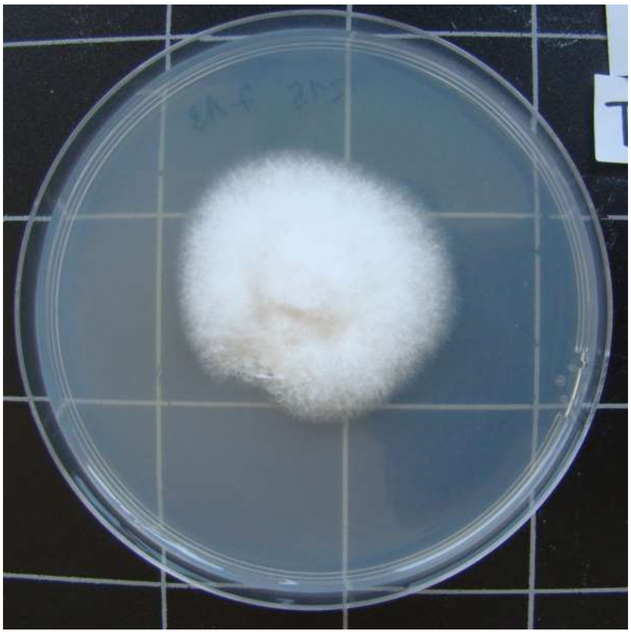

1215-TTF7.A-3

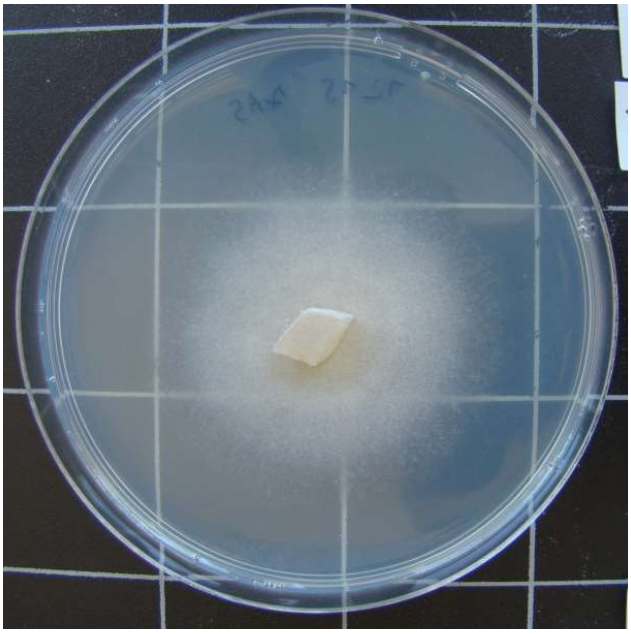

1215-TTF7.A-5

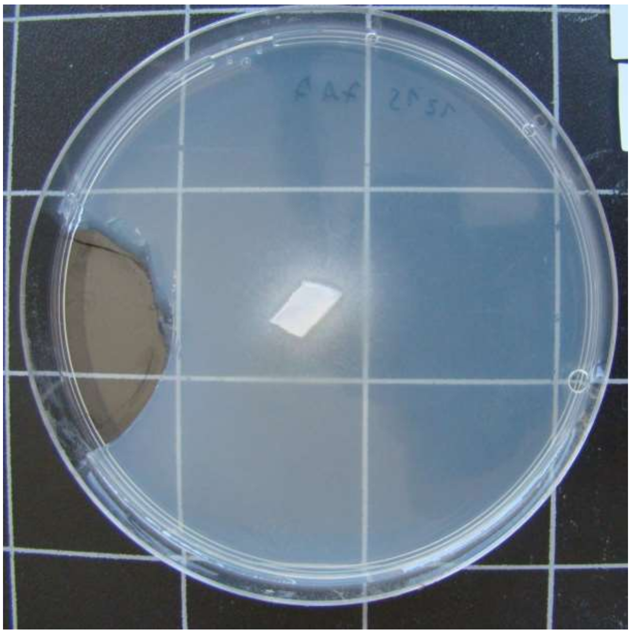

1215-TTF7.A-7

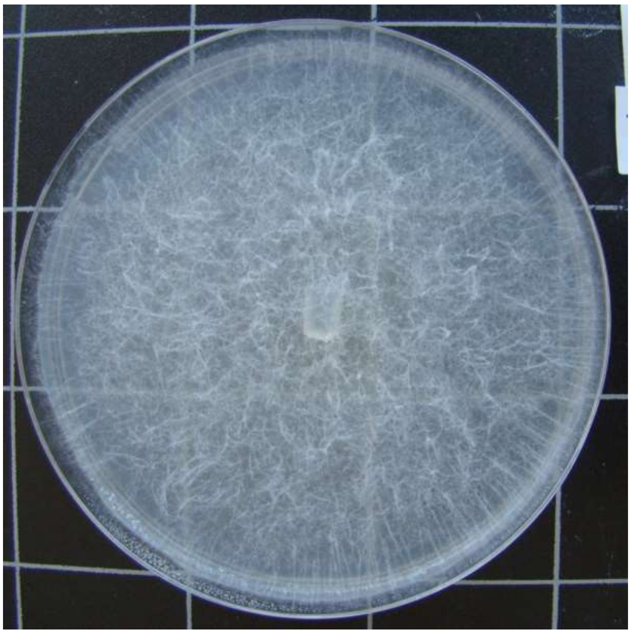

1215-TTF7.A-8

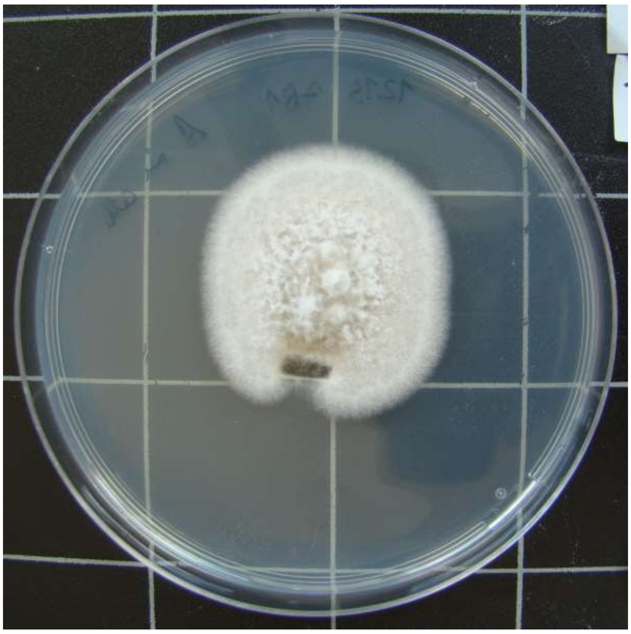

1215-TTF7.B-1

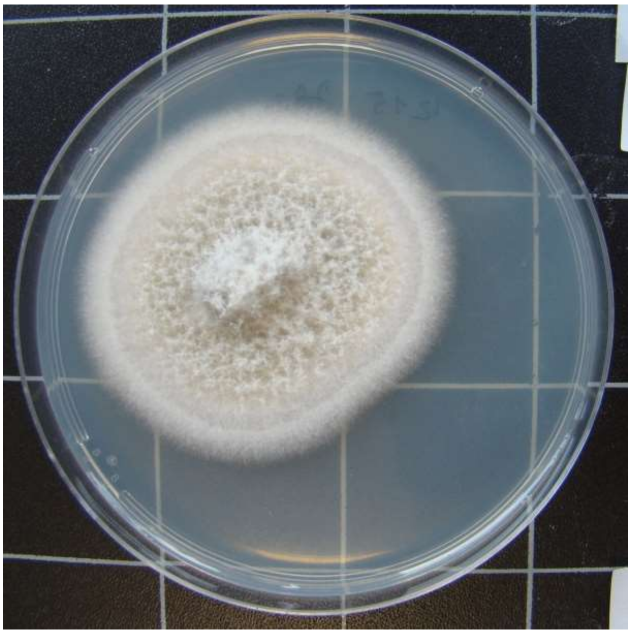

1215-TTF7.B-2

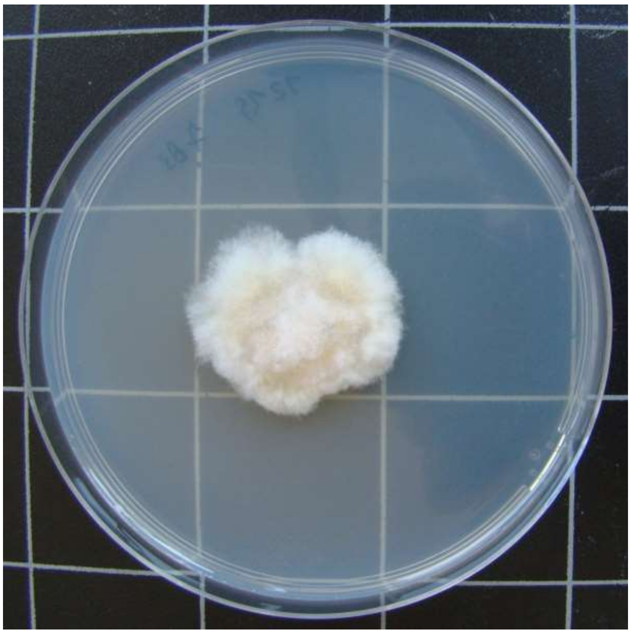

1215-TTF7.B-3

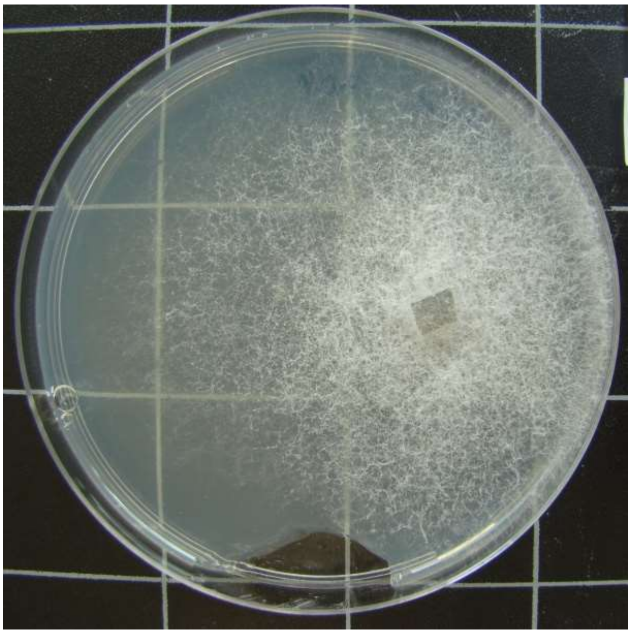

1215-TTF7.B-4

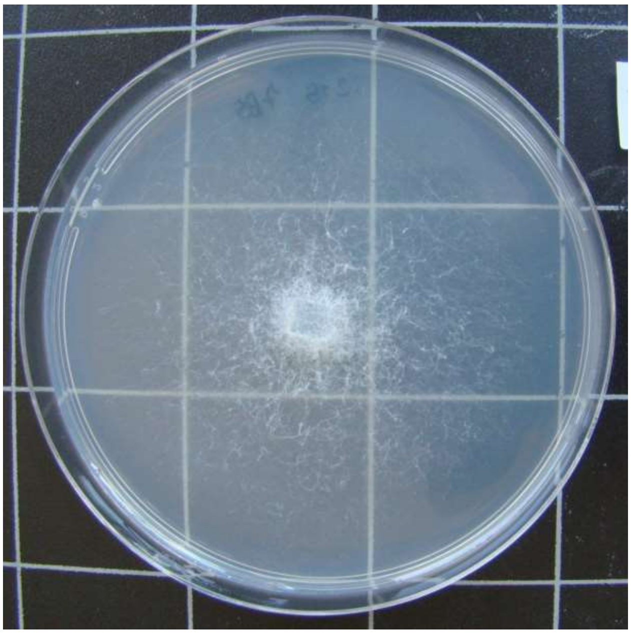

1215-TTF7.B-5

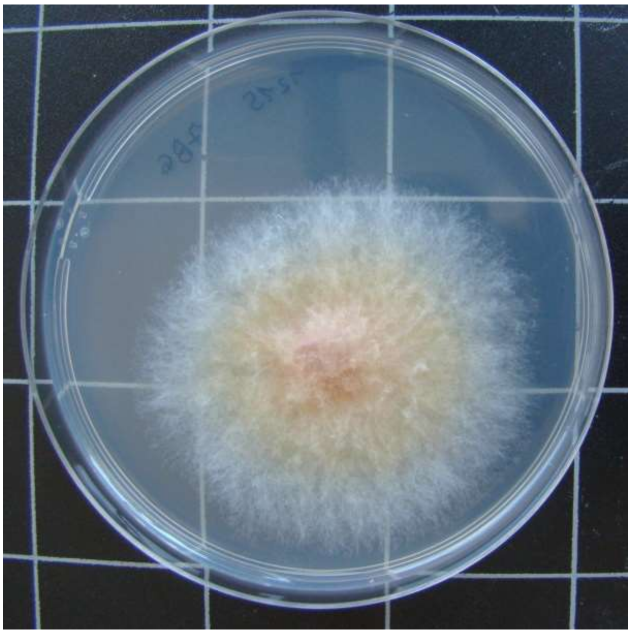

1215-TTF7.B-6

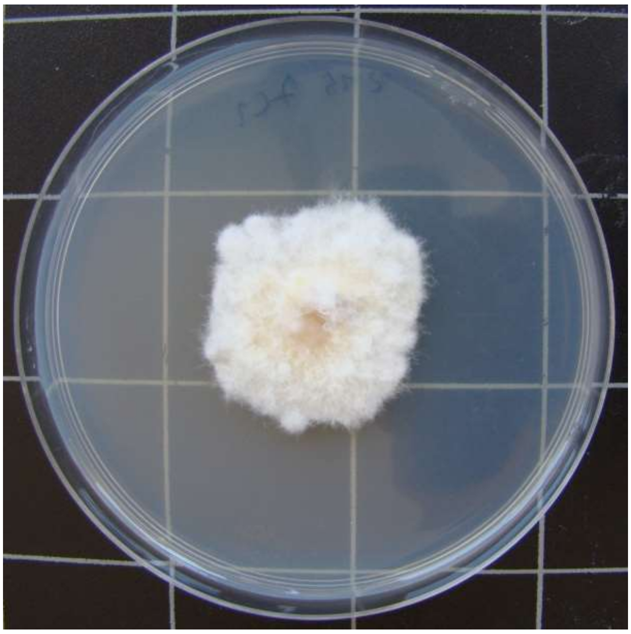

1215-TTF7.C-1

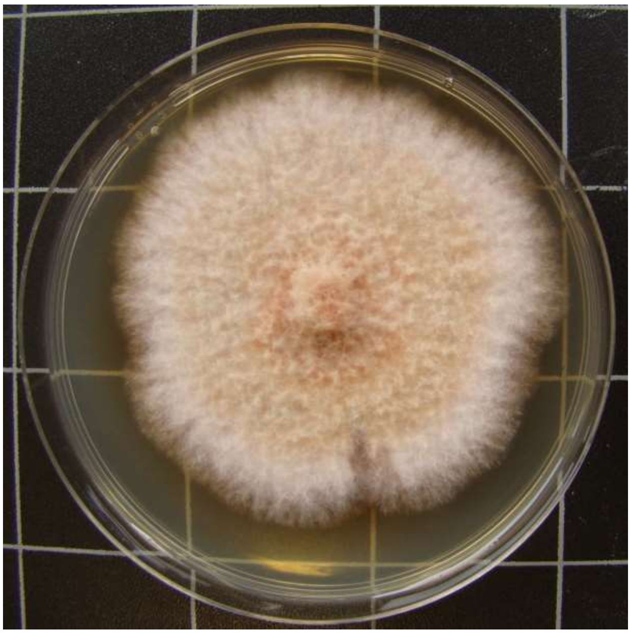

1215-TTF7.C-3

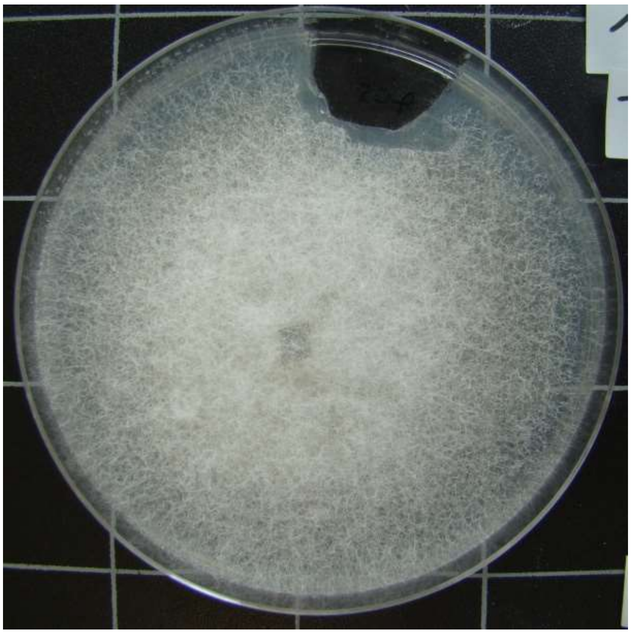

1215-TTF7.C-5

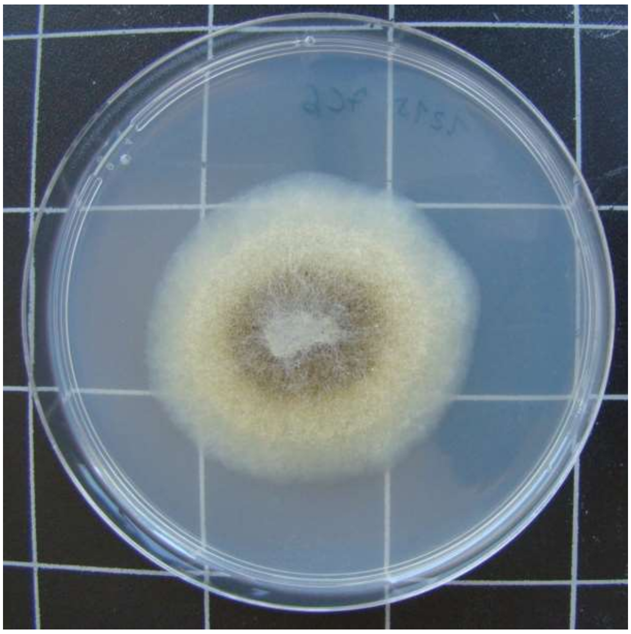

1215-TTF7.C-6

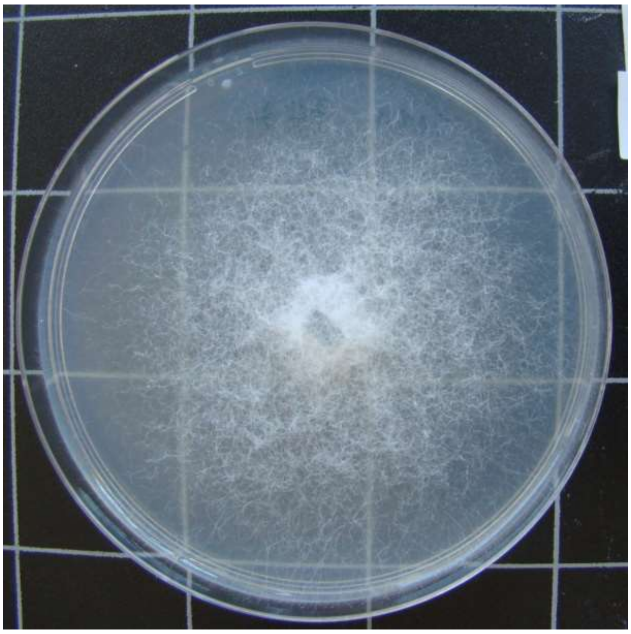

1215-TTF7.C-7

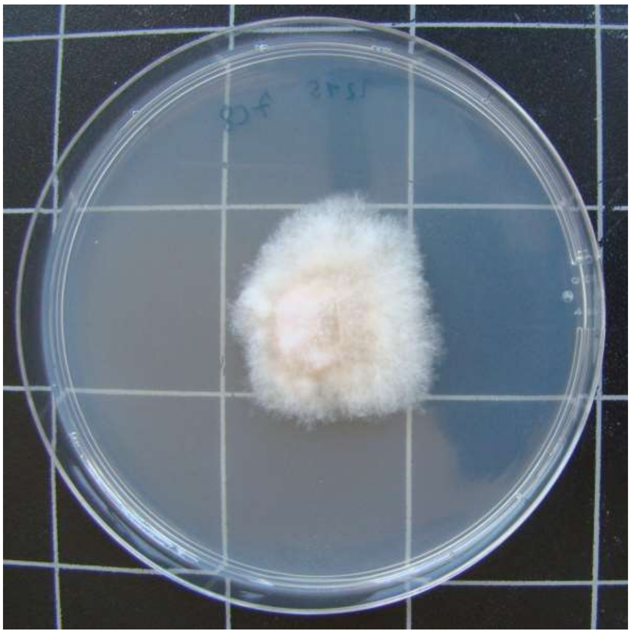

1215-TTF7.C-8

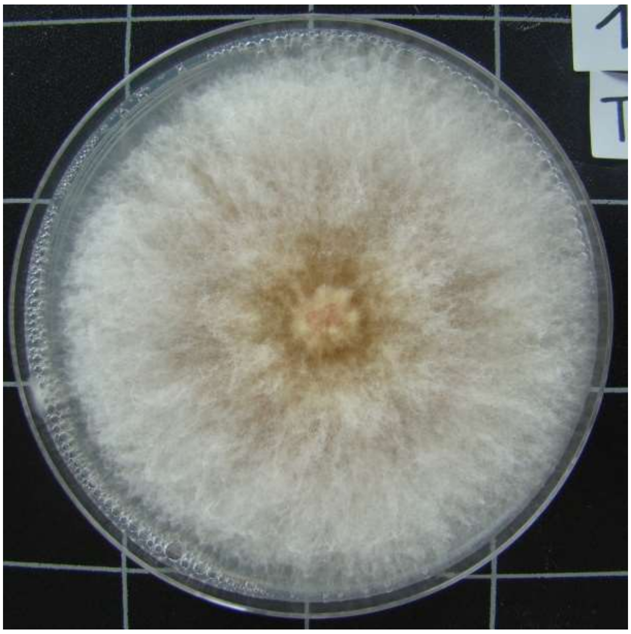

1215-TTF7.C-9

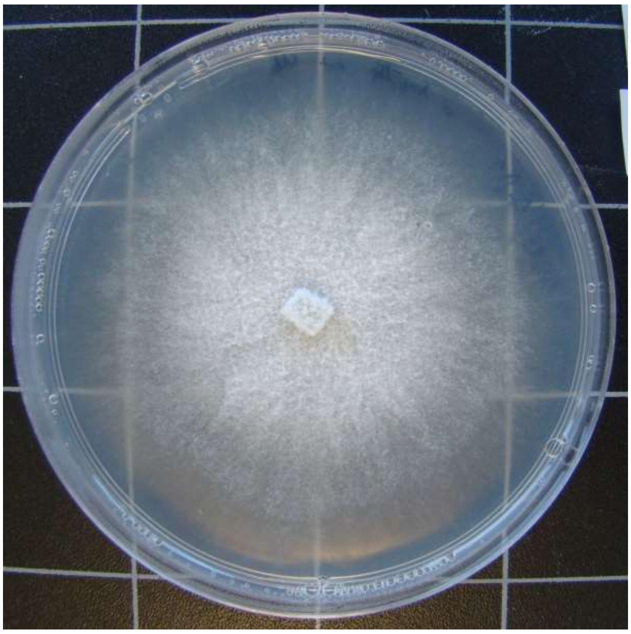

1215-TTF7.D-1

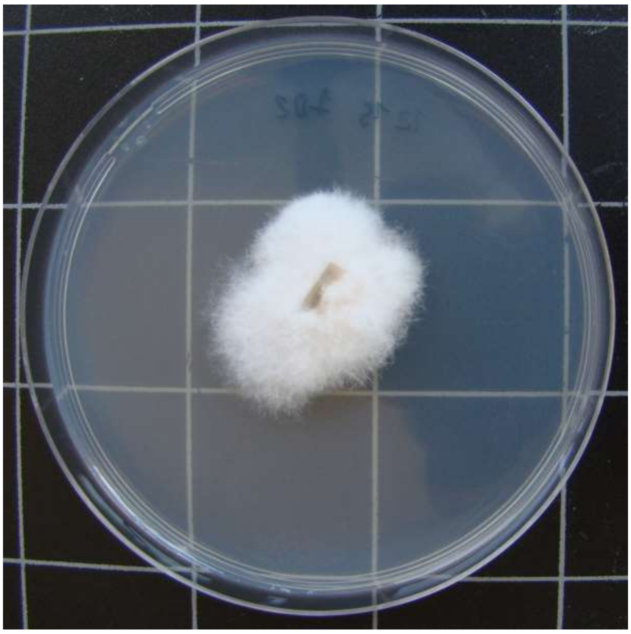

1215-TTF7.D-2

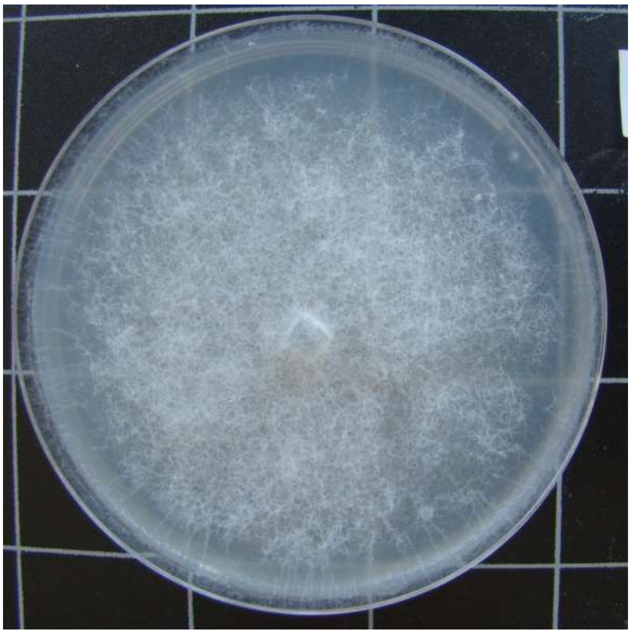

1215-TTF7.D-3

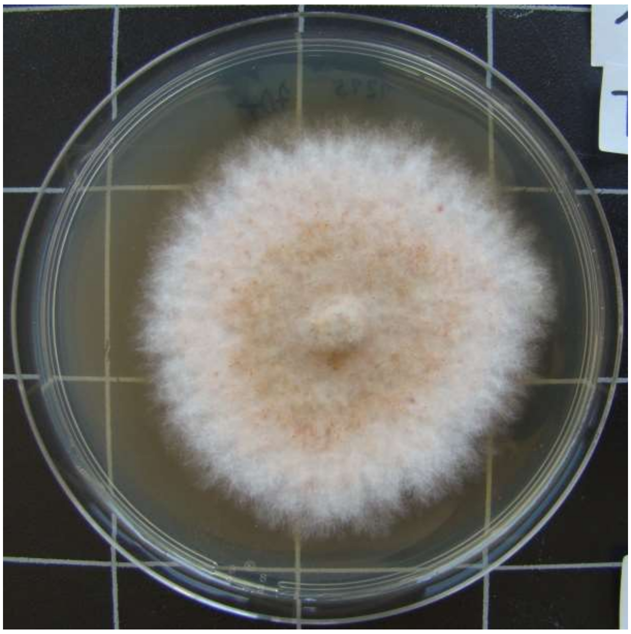

1215-TTF7.D-4

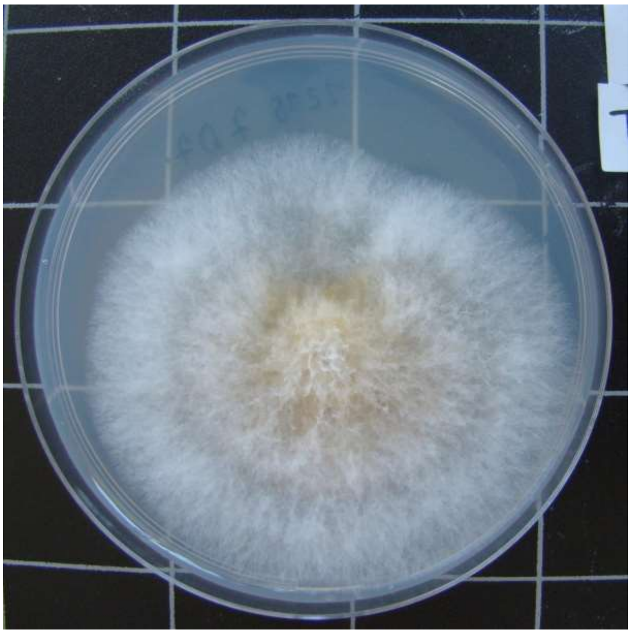

1215-TTF7.D-7

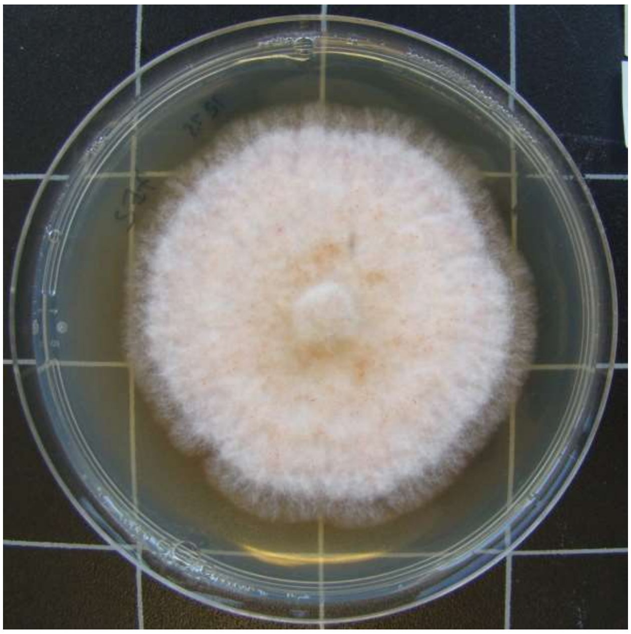

1215-TTF7.E-2

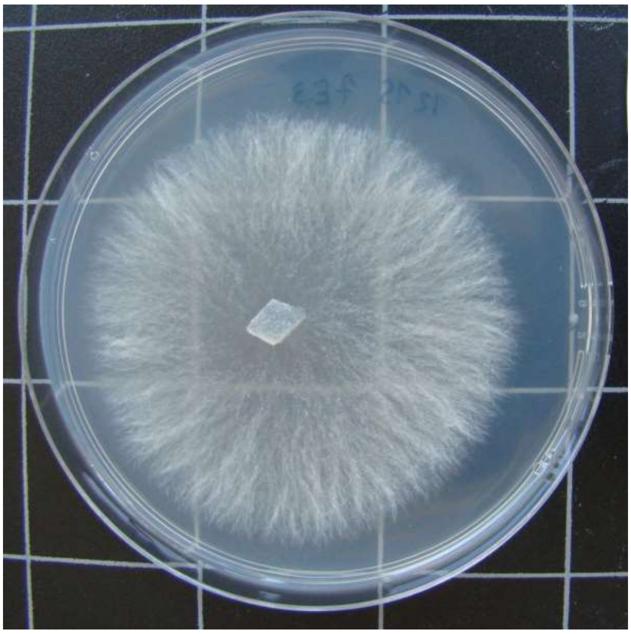

1215-TTF7.E-3

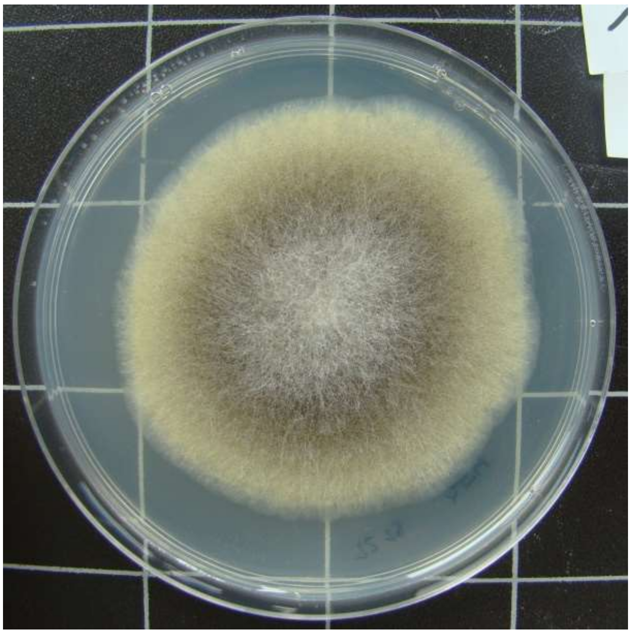

1215-TTF7.E-4

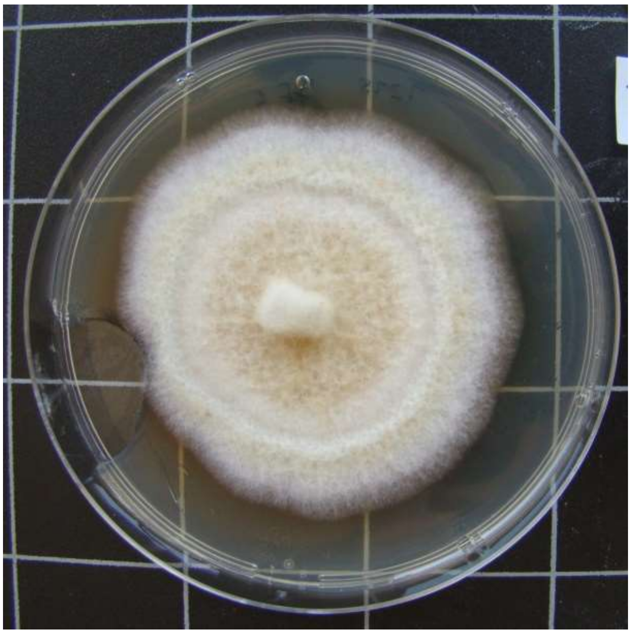

1215-TTF7.E-5

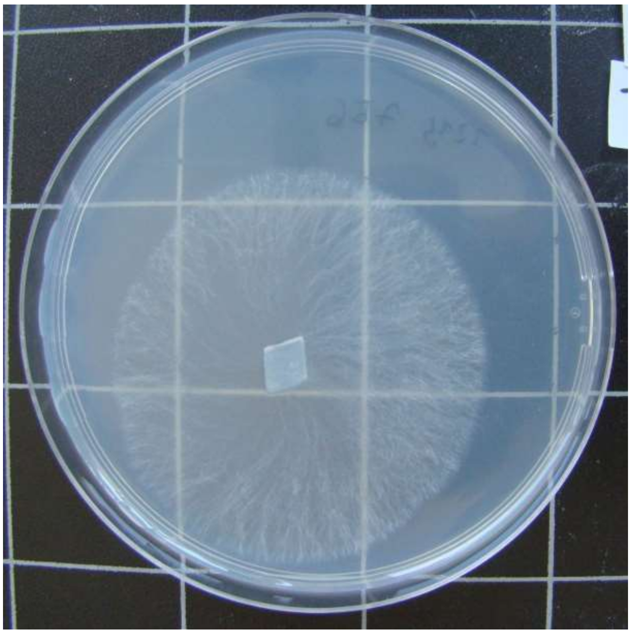

1215-TTF7.E-6

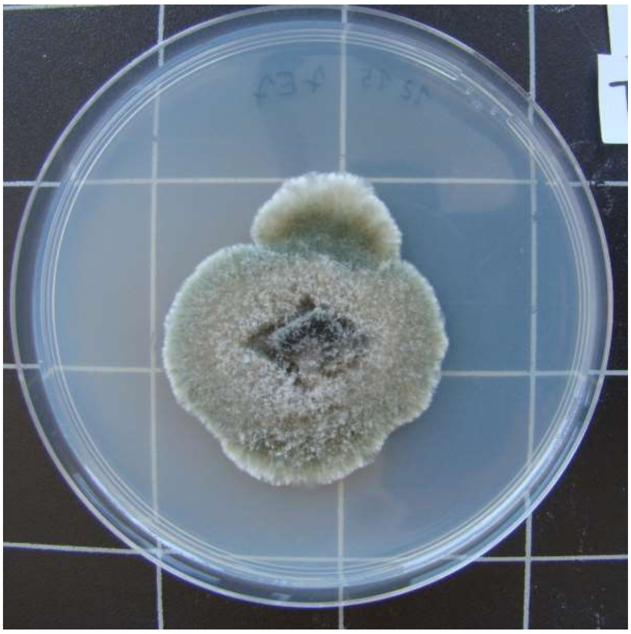

1215-TTF7.E-7

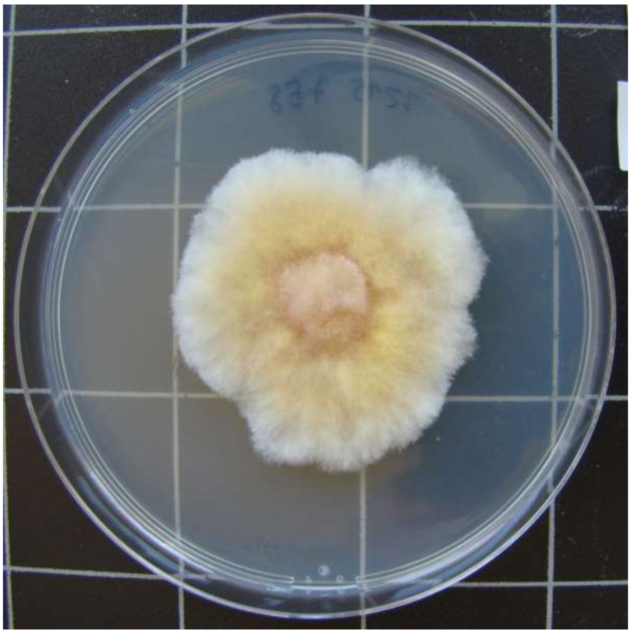

1215-TTF7.E-8

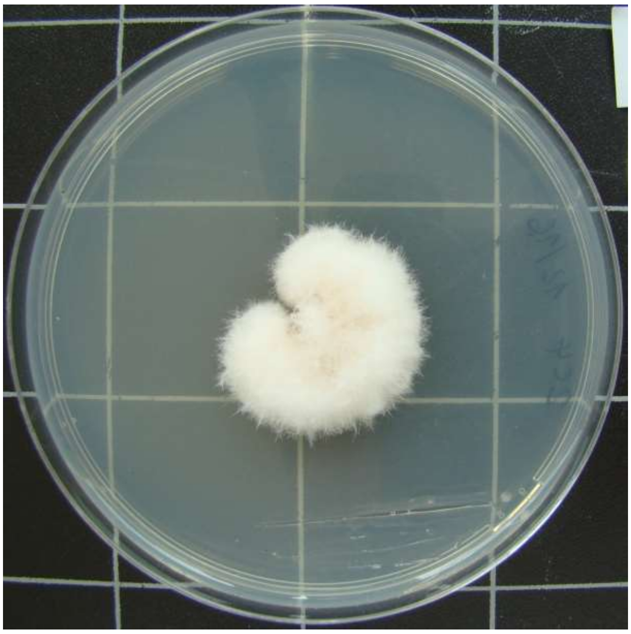

1216-HDG2.C-4

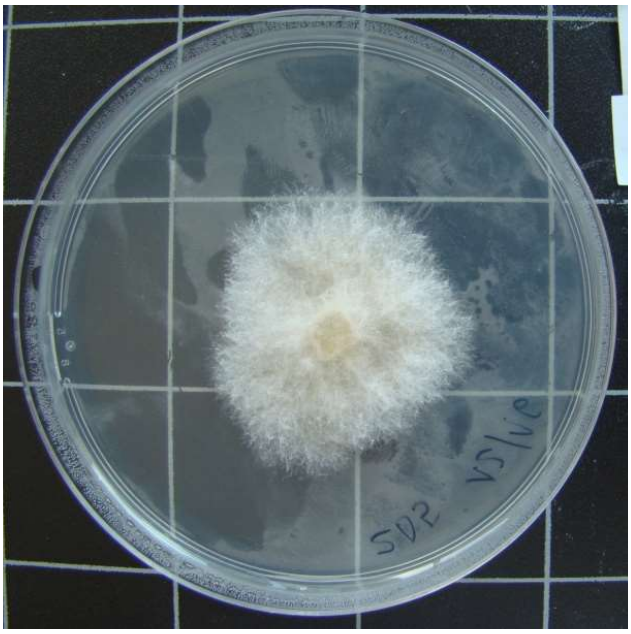

1216-HDG2.D-5

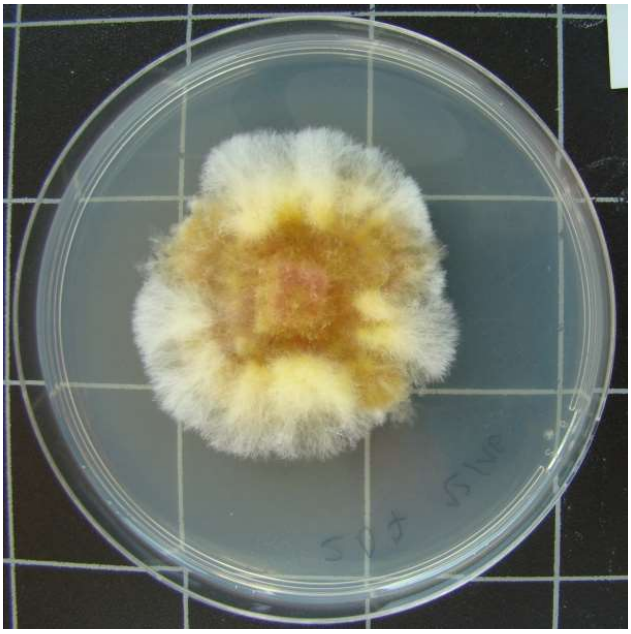

1216-HDG2.D-7

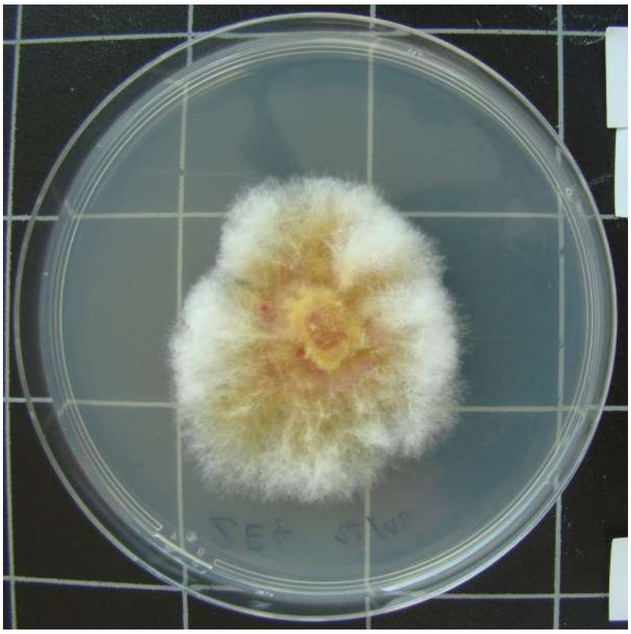

1216-HDG2.E-4

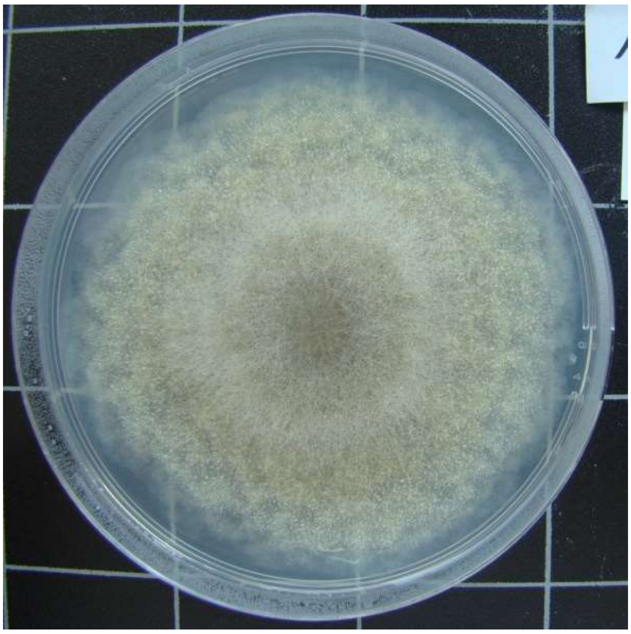

1216-HDG3.A-5

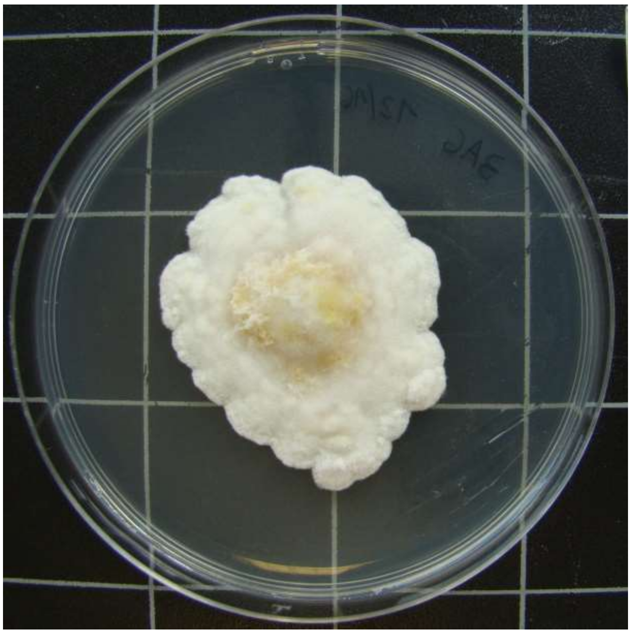

1216-HDG3.A-6

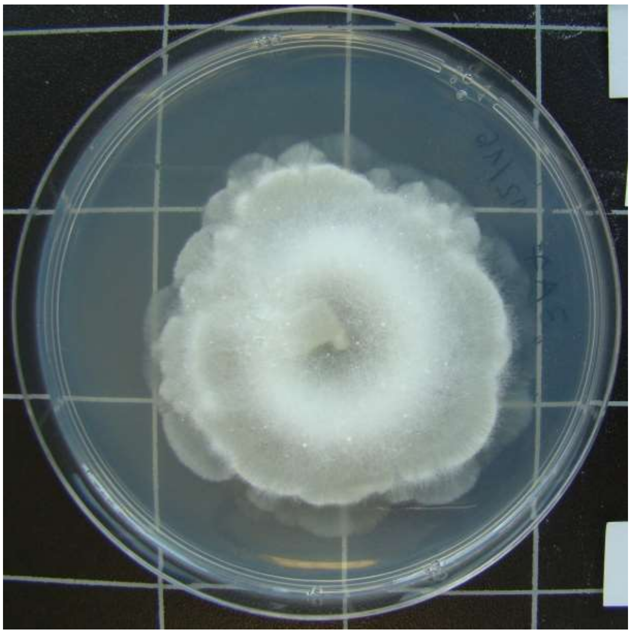

1216-HDG3.A-7

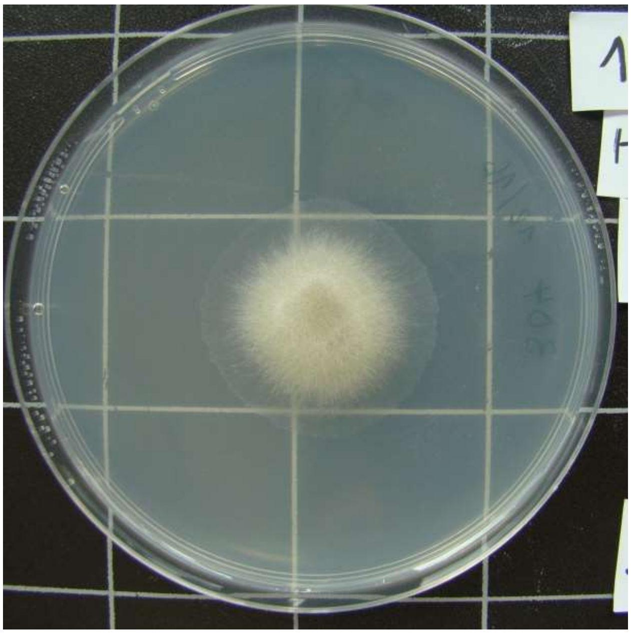

1216-HDG3.D-4

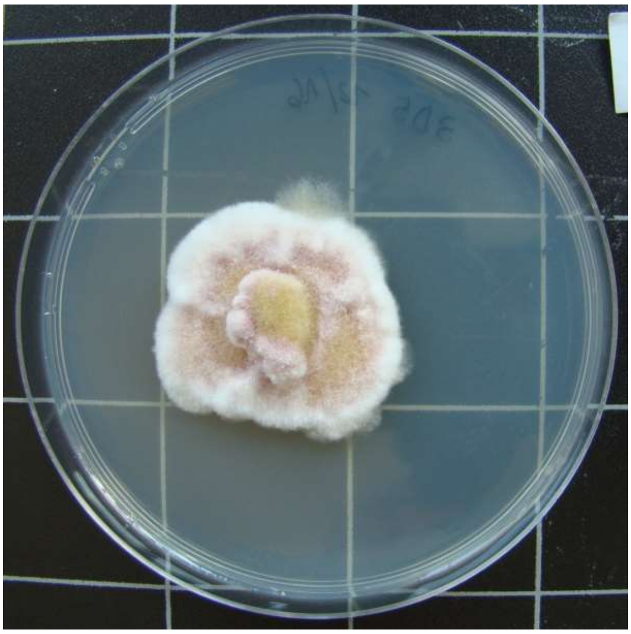

1216-HDG3.D-5

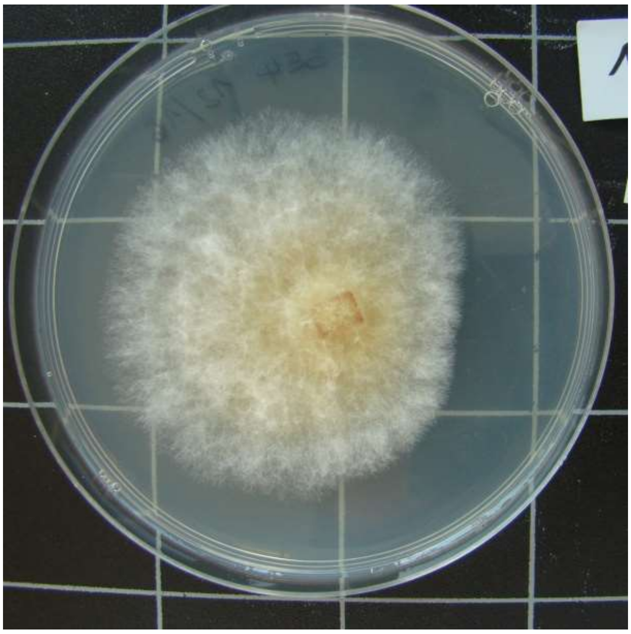

1216-HDG3.E-4

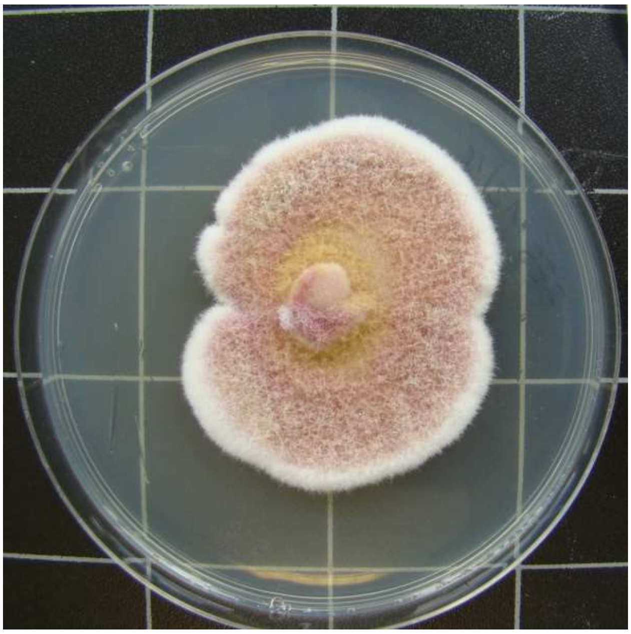

1216-HDG3.E-5

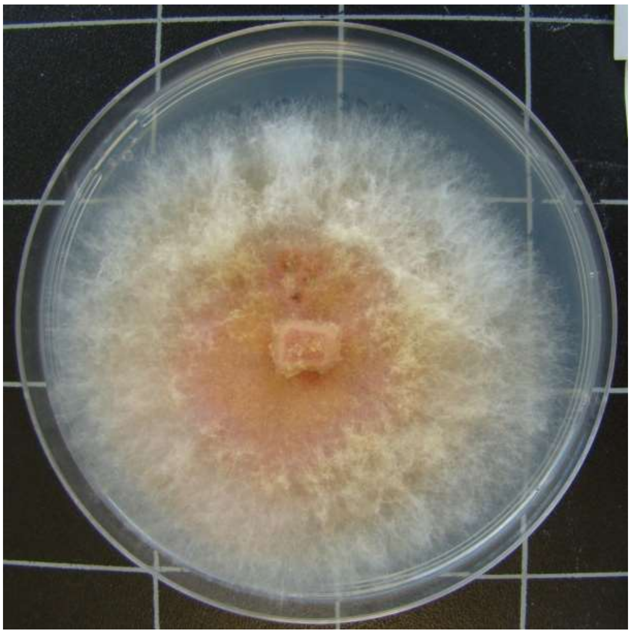

1216-TTF7.B-9

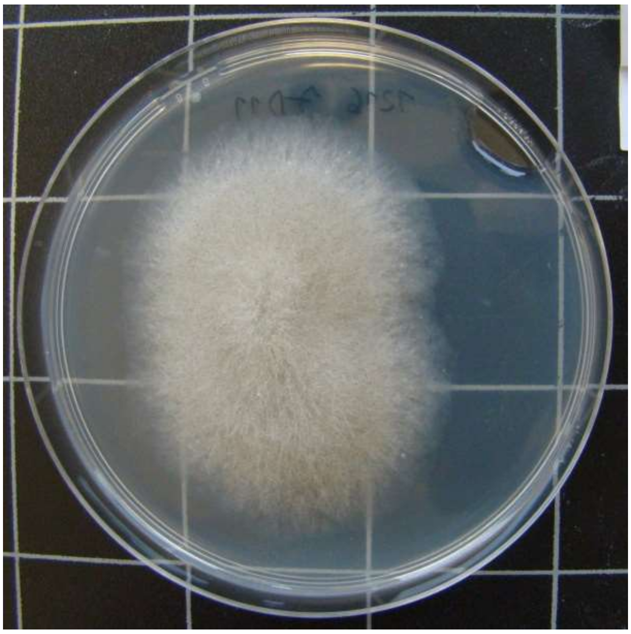

1216-TTF7.D-11
